# Dynamic spatiotemporal activation of a pervasive neurogenic competence in striatal astrocytes supports continuous neurogenesis following injury

**DOI:** 10.1101/2024.04.16.589779

**Authors:** Marco Fogli, Giulia Nato, Philip Greulich, Jacopo Pinto, Paolo Peretto, Annalisa Buffo, Federico Luzzati

## Abstract

Adult brain neural stem cells (NSCs) are conventionally regarded as rare cells restricted to two niches: the subventricular zone (SVZ) and the subgranular zone. Parenchymal astrocytes can also contribute to neurogenesis after injury, however the prevalence, distribution, and behaviour of these latent NSCs remained elusive. To tackle these issues, we reconstructed the spatiotemporal pattern of striatal astrocytes neurogenic activation after excitotoxic lesion in mice. Our results indicate that a neurogenic potential is broadly distributed throughout the striatum but is focally activated at the lesion border. In this region, similarly to canonical niches, steady state neurogenesis is ensured by the continuous stochastic activation of local astrocytes. Activated astrocytes quickly return to quiescence, while their progeny locally proliferate for about 10 days following a stochastic behaviour that features an acceleration in differentiation propensity. Notably, striatal astrocytes activation rate matches that of SVZ astrocytes indicating a comparable prevalence of NSC potential.

## INTRODUCTION

Stem cells are typically rare cells confined to specialised anatomical regions, called niches, that ensure their maintenance and regulate their activity^1^. In the adult mammalian brain, two such regions have been identified, the SubVentricular Zone (SVZ) and the SubGranular Zone (SGZ), where astroglial-like cells retaining the monolayer arrangement and apico-basal polarisation typical of embryonic radial glial progenitors, act as Neural Stem Cells (NSCs) throughout life^2–4^. These cells are mostly quiescent (qNSC) but sporadically activate generating transit amplifying progenitors (TAPs) that further divide before differentiating into neuroblasts (NBs)^5,6^. The stochastic activation of widespread qNSCs within the niche ensures continuous neuron production^7–11^. Outside these niches, the mature brain parenchyma has been traditionally considered non-permissive for neurogenesis due to a lack of neurogenic competence in parenchymal astrocytes and to the presence of gliogenic factors^12–15^. However, single cell RNAseq revealed striking similarities between parenchymal astrocytes and qNSC^16,17^ and all major elements maintaining stem cells in canonical niches are present in the parenchyma^14,18,19^. Parenchymal astrocytes may thus represent NSCs in a deep quiescent state. According to this notion, during early postnatal development^20^ or after injury subsets of parenchymal astrocytes can expand *in vitro* as neurospheres^21–24^ and become transcriptionally similar to NSC primed for activation^25^. Notably, in the mouse striatum, some astrocytes move beyond the primed state and express their neurogenic capacity *in vivo,* after stroke or quinolinic acid (QA) mediated excitotoxic lesion, supporting neurogenesis for several months^26–28^. However, the prevalence, spatial distribution and dynamics of these ectopic NSCs were not resolved. Consequently, how widespread is NSC potential among parenchymal astrocytes and to what extent the parenchyma is permissive for its maintenance and expression remain unclear. Striatal neurogenic astrocytes may simply represent a new rare NSCs population. Accordingly, the mainstream view in the field still adheres to the concept of the anatomical restriction of NSCs potential^29^. To address these issues, we investigated the spatiotemporal dynamics of neurogenic activity and lineage progression of striatal astrocytes after QA lesion^26^.

To reach this goal, we built upon our previous demonstration that in this model new striatal neurons originate exclusively from striatal astrocytes through neurogenic foci scattered around the lesion border^26^. These foci are clusters of proliferating Ki67^+^ cells (Ki67^+^clusters), including TAPs-like cells and proliferating NBs (pNBs), along with early post-mitotic NBs^26^. The current study demonstrates that, similarly to canonical niches, these neurogenic foci are transient structures continuously generated by the stochastic activation within a widespread population of quiescent astrocytes residing in a globally permissive environment. Collectively our results reveal an unprecedented neurogenic potential within the adult brain parenchyma, comparable to that found in neurogenic niches.

## RESULTS

### Neurogenic foci organise in a 3D germinal matrix centred around the lesion border

To analyse the spatial distribution of the Ki67^+^clusters, defined as groups of at least 4 cells in direct contact^26^, we 3D reconstructed the striatum of 7 specimens at 5 weeks post lesion (wpl). As we previously described^26^, QA lesions resulted in the loss of striatal neurons in a large dorsolateral domain that was filled by densely arranged GFAP^+^ reactive astrocytes (Figures 1A-1A’). Ki67^+^clusters organised in a 3D germinal matrix centred around the rostro-medial part of the lesion border in both lesioned and spared tissue (Figures 1A’’-1C and S1A-S1B; Video S1).

**Figure 1.**
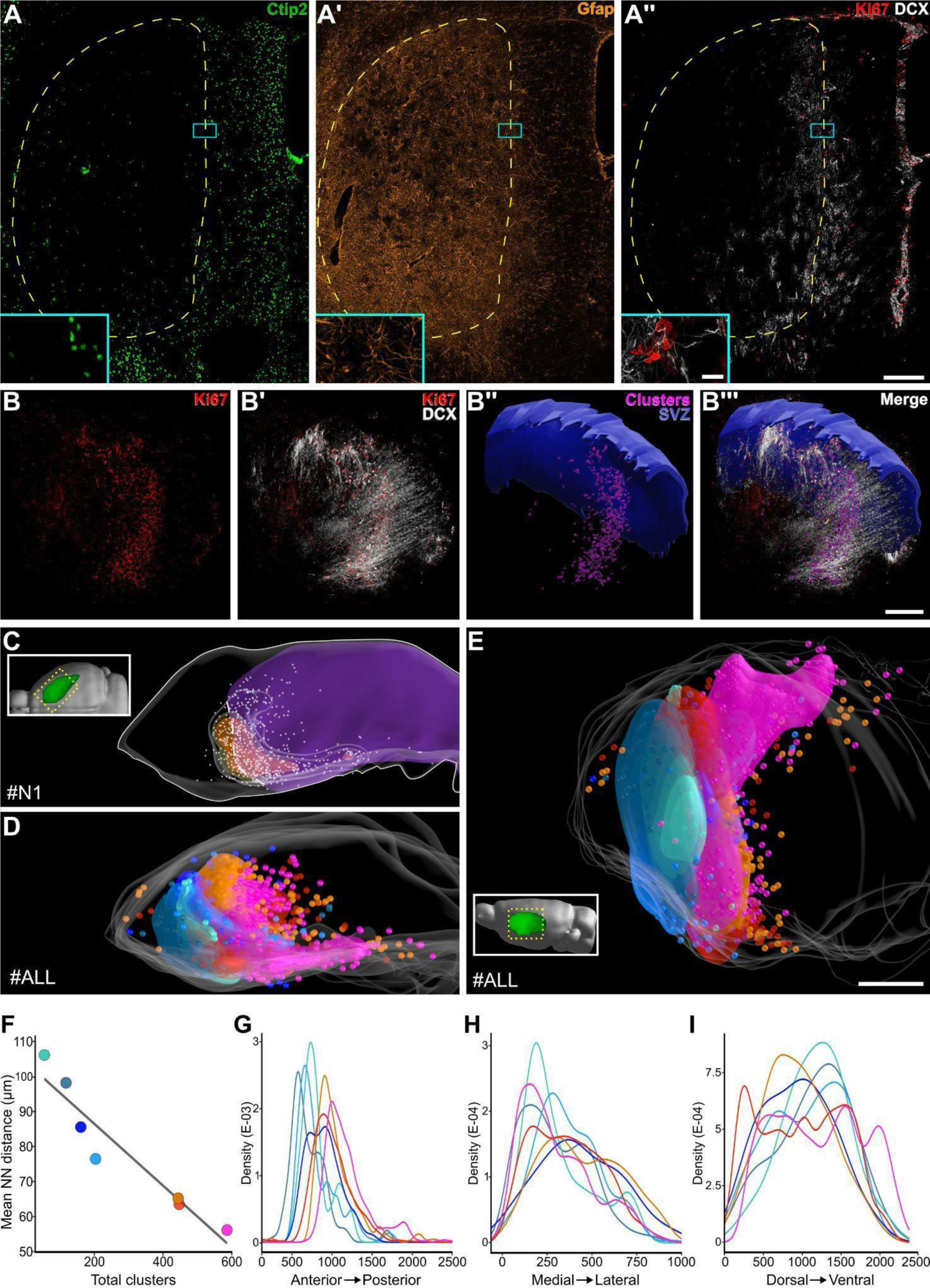
Neurogenic foci form a 3D germinal matrix. (A-A’’) Ctip2, GFAP, Ki67 and DCX stained section. Dashed yellow line: lesioned area. Inset: high magnification of a Ki67^+^cluster. (B-B’’’) 3D reconstruction of 40 serial sections stained with Ki67 and DCX (specimen #N1; voxel size 0.76 x 0.76 x 3.78 µm; see also Video S1). SVZ DCX^+^ and Ki67^+^ cells are hidden. In (B’’-B’’’) segmented striatal Ki67^+^clusters are in magenta. (C) Dorsal view of #N1 striatum 3D reconstruction. The lesion is in purple. White dashed line: the lesion border in the neurogenic area. Grey dots: Ki67^+^clusters (25 µm diameter). Increasing relative cluster density is rendered as transparent, yellow and orange volumes. (D,E) 3D reconstructions of 7 specimens registered to a common coordinate frame. The Ki67^+^clusters (100 µm radius) and the volume rendering of cluster density are coloured by specimen as in Figures S1A-S1B and in (F,G,H,I). (F) Correlation between the number of clusters and mean nearest-neighbour distance among them. Black line: linear regression. (G,H,I) Relative Ki67^+^cluster distribution along the antero-posterior (G), medio-lateral (H) and dorso-ventral (I) axis. Scale: (A) 200 µm; (inset A) 20 µm; (B-B’’’), (C), (D) and (E) 500 µm.

The DCX^+^ postmitotic NBs generated by these neurogenic foci further extended rostrally and caudally, partly inside white matter tracts (Figures 1B’ and 1B’’’; Video S1). The peak density of Ki67^+^clusters was always well distanced from the SVZ, in line with the striatal origin of these structures (Figures 1C and S1A). The number of Ki67^+^clusters varied greatly, ranging from 55 to 586 (mean±SD= 288±202) and correlated with their mean nearest neighbour distance (Figure 1F; Table S1, *p<0.001*), indicating that stronger neurogenic responses resulted in increased foci density within similar areas. The position of the neurogenic areas varied based on the lesion border. When all specimens were registered together, these areas collectively occupied most of the rostral and medial striatum (Figures 1D-1E, 1G-1I and S1A-S1B; Video S1). In particular, the neurogenic foci were preferentially distributed in the medial striatum (Figure 1H), an associative functional domain corresponding to the caudate nucleus of other mammals^30,31^ (Figures S1A’ and S1B’). By contrast, these structures were more sparse in the lateral somato-motor striatum and completely absent in the caudal multi-modal domain^30,31^ (Figure S1A).

In summary, striatal neurogenic foci organise in a complex 3D germinal matrix whose spatial disposition is contingent on the lesion border. The potential of neurogenic foci induction is widespread in the striatum with a higher probability in its medial domain.

### Neurogenic foci exist along a continuum of maturation profiles

Neurogenic foci have been described in different models of striatal neurogenesis, but their individual cellular composition has never been resolved^26,27,32,33^. The size and fraction of TAPs and pNBs in these foci could (i) vary along a continuum of maturation profiles as in adult neurogenic niches, or (ii) could be more invariant as in stem cell systems, like the skin, where continuous activity at fixed locations maintains a stationary proportion of maturation stages^34^. To distinguish between these models, here we evaluated the TAPs (DCX^-^Ki67^+^) and pNBs (DCX^+^Ki67^+^) content of 430 Ki67^+^clusters 3D reconstructed from serial sections (n=8 mice; Figures 2A and S2A-S2E).

**Figure 2.**
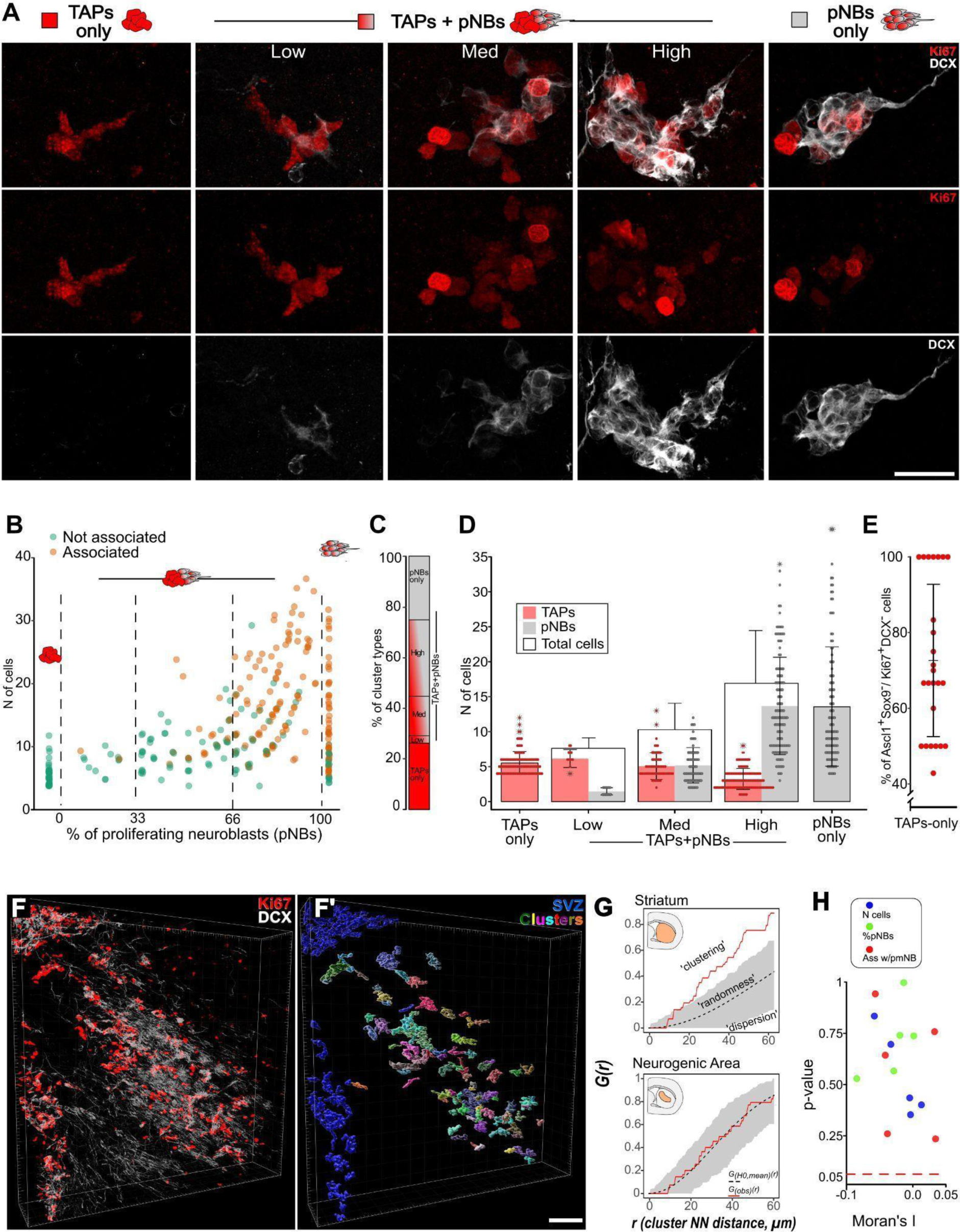
Neurogenic foci exhibit heterogeneous cellular composition and are spatially independent. (A) Ki67^+^clusters stained for DCX and Ki67. From left to right: TAPs-only, TAPs+pNBS_Low; TAPs+pNBs_Med; TAPs+pNBs_High and pNBs-only clusters (see also Figure S2A-S2E). (B) Percentage of pNBs vs number of cells per cluster; clusters associated with postmitotic NBs are in orange. (C) Proportions of cluster types among reconstructed clusters. (D) Mean±SD number of TAPs (red) and pNBs (gray) per cluster type (see also Table S1). (E) Percentage of TAPs expressing Ascl1 in the 25 TAPs-only clusters analysed. (F,F’) Perspective view of a 3D reconstructed portion of a neurogenic area. (F’) shows Ki67^+^clusters volume segmentation in random colours (see also Video S2). (G,H) Representative G functions of Ki67^+^cluster distribution with respect to the striatum or the neurogenic area of the specimen in (F). Grey areas: pointwise envelopes of 999 simulations of complete spatial randomness. *G_(obs)_(r)* (red line): experimental value; *G_(H0,mean)_(r)* (black dashed line): mean of the simulations (see also Figure S3A-S3B for formal analyses). (H) Moran’s I index and associated p-value for each tested specimen (n=5) and variable (n=3) (see Methods details and Table S1). Scale: (A) 25 µm (F) 100 µm.

The Ki67^+^clusters varied greatly in size, ranging from 4 to 38 cells (mean±SD: 11.8±7.6 cells), and also in the proportion of TAPs and pNBs (Figures 2A-2B). About half of the Ki67^+^clusters were composed either of only TAPs (“TAPs-only clusters”) or only pNBs (“pNBs-only clusters”) while the other half included a mixed population of the two cell types (“TAPs+pNBs clusters”; Figures 2A-2C). We further subdivided TAPs+pNBs clusters into TAPs+pNBs_Low, TAPs+pNBs_Med and TAPs+pNBs_High according to the pNB fraction (Figures 2A-2C). Both the cluster size and the number of pNBs increased with the pNBs fraction, progressing from TAPs-only to TAPs+pNBs_High clusters (Figure 2D; Table S1, size: *p<0.001*; pNBs: *p<0.001*). Conversely, TAPs exhibited an opposite trend (Figure 2D; Table S1, *p<0.001*). The increase in pNBs content and size further correlated with a higher probability of postmitotic NBs (DCX^+^Ki67^-^) being part of the neurogenic foci (Figure 2B; Table S1, % of pNBs: *p<0.001*; size: *p<0.001*). TAPs-only clusters were never associated with postmitotic NBs, while the probability of this association progressively rose from TAPs+pNBs_Low to pNBs-only clusters (Figure S2F; Table S1, *p<0.001*). Of note, in all the TAPs-only clusters most of the cells expressed the TAPs marker Ascl1^15,35^ but not the astrocyte marker SOX9^36,37^ (Figures 2E and S2G-S2G’), confirming the neuronal commitment of these structures.

These data show that striatal neurogenic foci are highly diverse in their maturation profiles. This heterogeneity takes the form of a continuous spectrum of transitions, suggesting that different Ki67^+^cluster types represent sequential stages of a common developmental process. These structures may be initially composed only of TAPs and progressively accumulate NBs that at first proliferate and gradually become postmitotic, eventually depleting the proliferating pool. Alternatively, this variability may be contingent to spatial or temporal factors.

### Neurogenic foci are spatially independent

Striatal neurogenic foci heterogeneity could result at least in part from regional differences in lineage progression. To verify this possibility we performed a spatial analysis of 318 3D reconstructed Ki67^+^clusters (n=5 mice; Figure 2F-2F’; Video S2). Spatial point pattern analyses using the G function^38^ detected significant clustering of Ki67^+^clusters along the lesion border relative to the entire striatum, according to previous observations (Figures 2G and S3A; *pooled p<0.001*). On the contrary, within the neurogenic area, striatal clusters were randomly distributed (Figures 2G and S3B; *pooled p=0.229*). We next employed a measure of global spatial autocorrelation, the Moran’s I index, to understand if the striatal cluster size, pNBs content and association with postmitotic NBs are distributed following specific spatial patterns (Figures S3C-S3C’’). For all the specimens the spatial distribution of these features did not deviate significantly from simulations of complete spatial randomness (Figure 2H; Table S1). Hence, we conclude that Ki67^+^clusters are randomly distributed in the neurogenic area and that their maturation profiles are spatially independent.

### Neurogenic foci number, distribution and cellular composition are stable after neurogenesis onset

To ascertain the temporal dynamics of striatal neurogenesis, Ki67^+^cluster number, distribution, and composition were analysed at 3, 4, 5, or 8 wpl. As observed at 5wpl, Ki67^+^clusters were always centred around the lesion border and their overall distribution did not vary with time (Figure S4A). Their total number varied among specimens, but, on the whole, it remained stable in time (Figures 3B-3E and S4A; Table S1, *p=0.157*). Of note, all the cluster types described by the 3D reconstruction analysis (Figures 2A-2C) were identified in all specimens and time-points (Figures S4B) and both their relative proportions and size were constant (Figures S4C-S4D; Table S1). These data show that shortly after their appearance, between 2 and 3 wpl, the number, distribution and composition of striatal neurogenic foci reach a stable level that is maintained up to 8 wpl. The stable heterogeneity of the Ki67+clusters strongly suggests they undergo continuous turnover.

**Figure 3.**
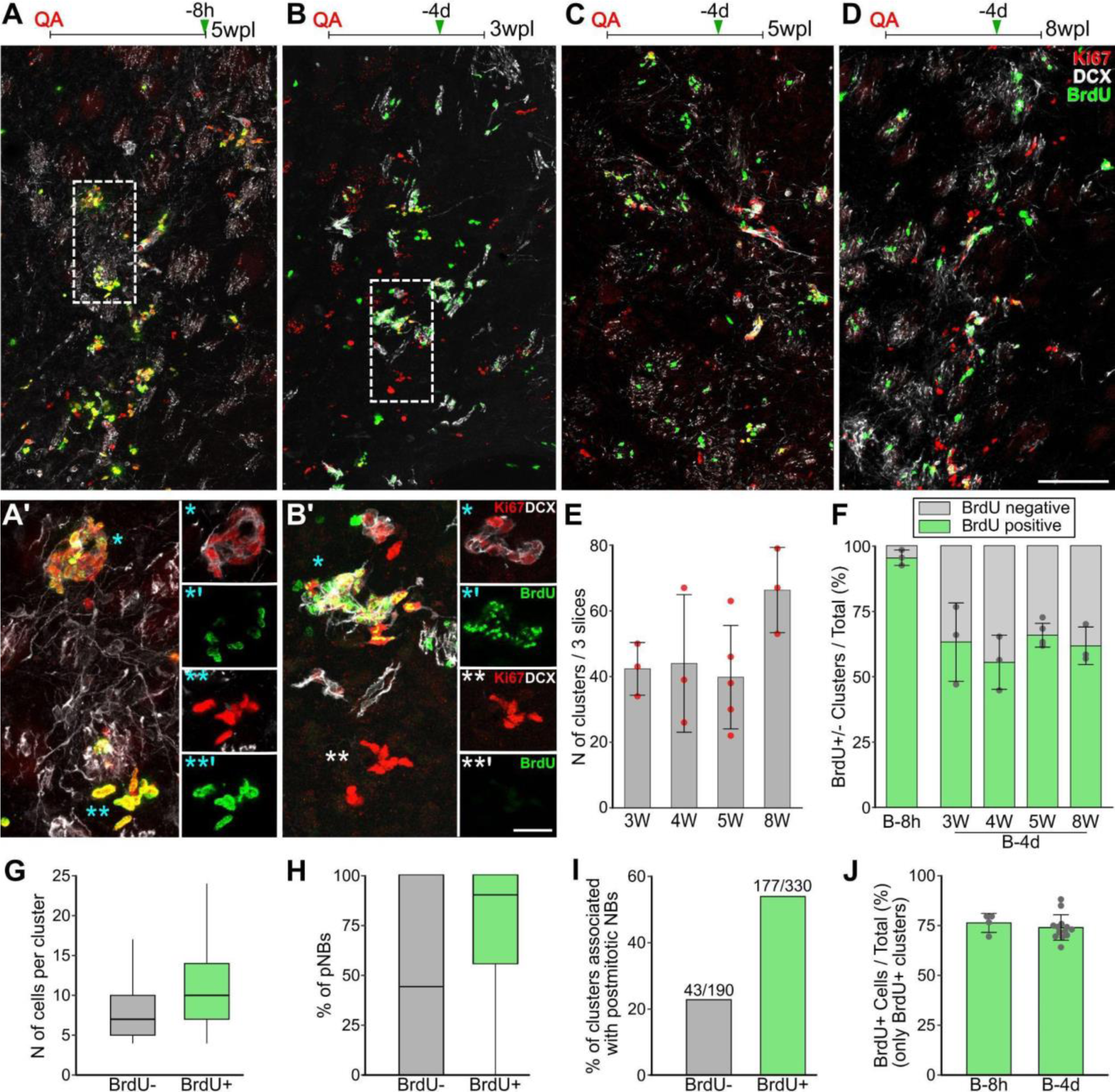
Neurogenic foci undergo a constant turnover. (A-D) BrdU, Ki67 and DCX labelling from a B-8h (A) and B-4d specimens at 3wpl (B), 5wpl (C) and 8wpl (D) in the neurogenic core (see also Figures S4A-S4D). (A’-B’) Higher magnification of A and B. MAX projections and single focal planes respectively in the left and right panels. * indicate more mature and ** more immature clusters. (E) Number of Ki67^+^clusters in three non-consecutive slices. (F) Percentage of BrdU^+^ clusters at B-8h and B-4d (see also Figures S5A-S5F). (G-I) Comparison (G, H: box-plot; I: bar-plot) of the number of cells per cluster (G), % of pNBs per cluster (H) and the fraction of Ki67^+^clusters associated with postmitotic NBs (I) between the BrdU^-^ and the BrdU^+^ clusters at B-4d. (J) Mean±SD fraction of BrdU+ cells in BrdU+ clusters in B-8h and B-4d specimens. See Table S1 for statistical analyses of (E-J). Scale: (A-D) 100 µm; (A’) and (B’) 20 µm.

### Neurogenic foci undergo continuous turnover

To assess the turnover rates of Ki67^+^clusters we first established a saturating BrdU protocol. Two BrdU injections at 8 and 2 hours before sacrifice (“B-8h” group) labelled 97±2% of the striatal clusters (Figures 3A and 3F) and 74±6% of their cells (Figures 3A’ and S5A) indicating that virtually all striatal clusters are made of actively dividing cells. As in SVZ^5^, TAPs had a slightly higher proliferation rate than pNBs (Figure S5B; Table S1, *p=0.040*), independent of the cluster maturation profile (Figure S5B; Table S1, *TAPs: p=0.125; pNBs: p=0.132*).

We then analysed the fraction of BrdU^+^ clusters after a four-day chase at 3, 4, 5 or 8 wpl (“B-4d” groups; Figures 3B-3D). If new Ki67^+^clusters were continuously generated, we expected that those established after BrdU administration would lack BrdU^+^ cells and exhibit an immature profile. According to this hypothesis, at all time points about 40% of the clusters were devoid of BrdU^+^ cells (Figure 3F). The fraction of BrdU^+^ cells and the frequency distribution of BrdU^+^ cells per cluster were stable in time (Figures S5C-S5D; Table S1, *p=0.512 and p=0.356 respectively*). Compared to BrdU^+^ clusters, the BrdU^-^ ones exhibited smaller sizes, lower fractions of pNBs, and less association with postmitotic NBs (Figures 3B’ and 3G-3I; Table S1, *p<0.001 in each case*), further supporting that they were established after BrdU injection.

However, BrdU^-^ clusters could also emerge at least in part by the loss of BrdU^+^ cells due to BrdU dilution under detection level or cellular turnover. Yet, when only the BrdU^+^ clusters were considered, neither the percentage of BrdU^+^ cells (Figure 3J, *p=0.319*) nor the frequency distribution of the BrdU^+^ cells per cluster varied between B-8h and B-4d (Figure S5E-S5F; Table S1, all clusters: *p<0.001*; BrdU^+^ clusters: *p=0.247*). Thus, neither BrdU dilution under detection levels nor cellular turnover occurred in the four-day chase, indicating that BrdU^-^ clusters were newly formed.

To sum up, for at least 5 weeks new neurogenic foci are continually established and undergo progressive maturation in the striatal parenchyma. Since their total number remains stable (Figure 3E), the genesis of new Ki67^+^clusters must be counterbalanced by the terminal differentiation of previously active ones. Neurogenic foci are thus transient structures for which we calculated from the proportion of BrdU^-^ clusters a lifetime of 10.6±2.1 days and a turnover rate of 9.4%/day (Supplementary Text 1, section 5).

Together with the lack of spatial patterns in cluster organisation, these data suggest that in the neurogenic area cluster initiation continuously occurs at random locations within a globally permissive environment.

### Clonal expansion of striatal astrocytes generates individual Ki67^+^clusters

Astrocytes could support the establishment of these transient clusters in several ways: each neurogenic foci could arise from individual or multiple astrocytes. In parallel, each astrocyte could expand locally to generate an individual clone or produce multiple migrating progenitors that secondarily expand, similar to metastatic tumour growth. To distinguish among these possibilities we conducted a multicolour clonal analysis of striatal astrocytes.

Glast^CreERT2/+^ mice expressing the inducible form of CRE recombinase under the pan astrocytic promoter Glast^39^, were crossed with R26R-Confetti reporter mice^40^ in which Tamoxifen administration leads to the stochastic and inheritable non-combinatorial expression of 1 out of 4 fluorescent proteins (Figures 4A and S6A). At 5 wpl, 15 days after Tamoxifen administration, 2.4±0.7%, 1.2±0.7% and 1.7±1.4% of the SOX9^+^ GFAP^+^ striatal astrocytes expressed RFP, mCFP and cYFP respectively (Figures S6B-S6E). In line with previous reports, we never observed the nGFP reporter^41^ (n= 1597 astrocytes, 4 mice). Consistent with the higher efficiency of confetti allele recombination in non-neural tissues^40^ we observed numerous recombined microglial cells that also express Glast. However, these cells were easily recognized by their morphology (Figures S6I-S6J) and excluded from our analyses.

**Figure 4.**
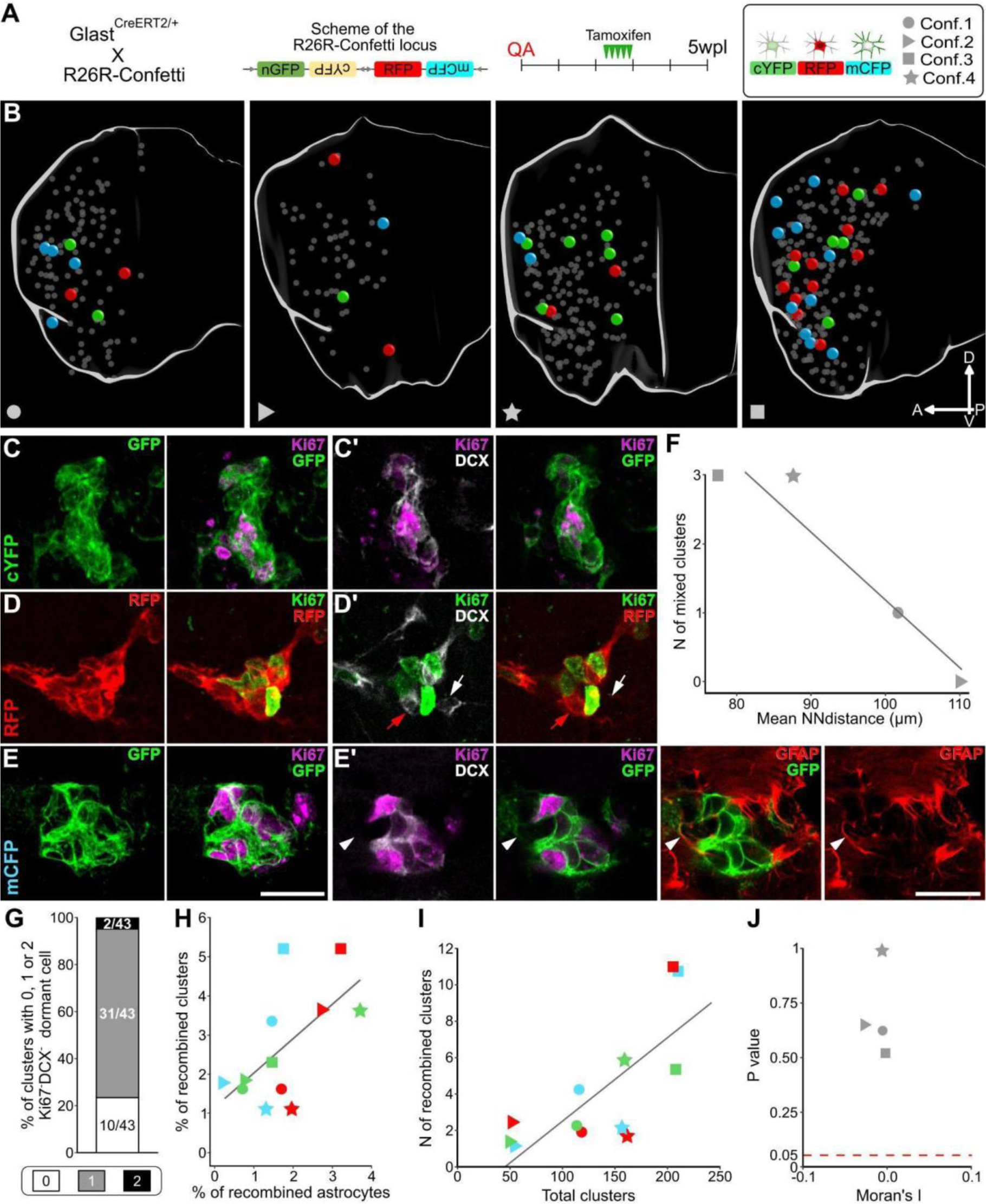
Clonal expansion of striatal astrocytes generates individual Ki67^+^clusters. (A) R26R-Confetti locus and experimental timeline. Right panel: reporter colour and specimen symbols legend. (B) Medial view of 3D reconstructions of the striatum (white contour) of 4 analysed specimens. Confetti clusters are depicted as bigger spheres coloured based on the reporter expression. Reporter^-^ clusters are smaller and grey. (C-E) MAX projection of clusters in which all the Ki67^+^ cells express the same reporter: cYFP, RFP and mCFP respectively. (C’-E’) Single focal planes of C, D and E respectively. Red and white arrows in (D’): colour-matching and unlabelled postmitotic NB, respectively. Arrowhead in (E’): dormant GFAP^+^ astrocyte. (F) Correlation between the mean nearest-neighbour distances among Ki67^+^clusters and the number of mixed clusters. (G) Percentage of Confetti clusters containing 0, 1 or 2 dormant Ki67^-^DCX^-^ cell. (H) Correlation between the total Ki67^+^clusters and the number of clusters expressing each reporter. (I) Moran’s I index and associated p-value calculated for each specimen (see Methods details and Table S1). Black Lines in (F) and (H) represent Linear Regression models. Scale: (C-E’) 20 µm.

Whole striatum 3D reconstruction of four specimens (Figure 4B) led to the identification of 537 Ki67^+^clusters (mean±SD= 134±63), 49 of which showed reporter expression (“Confetti clusters”; 8.4±3.3%; Figures 4C-4E’). The relative proportion of maturation profiles of Confetti clusters matched those previously described at 5wpl (Figure S6K, Table S1, *TAPs-only: p= 1; TAPs+pNBs: p= 0.454; pNBs-only: p=0.301*). Among Confetti clusters 42/49 were clones of cells expressing the same reporter (87±11%; Figures 4C-4E’) and thus originated from the same astrocyte. Non-recombined cells represented only 2% of all Confetti cluster cells (25/1048 of total counted cells) and were confined to 7 clusters. These few mixed clusters were found in the specimens with the lowest mean cluster nearest-neighbour distance (Figure 4F; Table S1, *p=0.042*), suggesting that increased density favoured the rare fusion of neurogenic foci.

Besides Ki67^+^ cells, recombined neurogenic foci with advanced maturation profiles included colour-matched postmitotic NBs (Figure 4D’). Interestingly, about 75% of Confetti clusters also included a clonally related Ki67^-^DCX^-^ cell (Figure 4G) that, in 70% of the cases, we could confirm being a GFAP^+^ astrocyte by performing quintuple immunostaining (Figure 4E’; 10/14 tested cells). Overall, these results demonstrate that striatal Ki67^+^clusters derive almost exclusively from the clonal expansion of a single astrocytic progenitor and suggest that this cell is maintained in a dormant state closely associated with its daughter cells.

Further, we addressed if a single astrocyte can give rise to multiple clusters. The number of clusters sharing the same reporter in each specimen was extremely low, ranging from 1 to 11 (mean±SD=4.1±3.6). When more than one cluster expressed the same colour, they were still within the expected values given the total number of clusters in that specimen (Figures 4H and S6F-S6H; Table S1, *p=0.004*). Moreover, Confetti clusters sharing the same reporter did not show any evident spatial association (Figure 4B), as confirmed statistically by Moran’s I and by permutation tests against simulations of complete spatial randomness (Figure 4I, Table S1). Sister Ki67^+^clusters originating from a common source are expected to exhibit closer proximity, while the absence of such spatial association indicates they are clonally independent structures deriving from distinct astrocytes. The presence of a dormant astrocyte within the neurogenic foci (Figures 4E’ and 4G) further supports this conclusion and indicates that these cells locally expand. Striatal neurogenic activity thus closely resembles that of canonical niches but distributed over a much wider 3D environment, thus leading to a negligible intermixing of different astrocyte progeny.

### Striatal astrocytes activate at a constant rate

Our results indicated that striatal astrocytes locally activate, generating individual Ki67^+^clusters that transiently expand. BrdU analysis did not detect any cellular turnover in neurogenic foci, strongly suggesting that astrocyte activation initiates new clusters that subsequently autonomously expand and mature while the primary progenitor returns to quiescence.

To fully demonstrate this model we used lineage tracing by dating back the astrocyte activation events contributing cells to 5wpl clusters. Different cohorts of Glast^CreERT2/+^xR26R-YFP animals were sacrificed at 5 wpl and received Tamoxifen 4, 7 or 14 days before sacrifice (T-4, T-7 and T-14; Figure 5A). At each time-point we measured the cluster labelling index, that is the fraction of clusters that received at least one YFP+ cell from their astrocytic progenitor (“YFP^+^ clusters”; Figures 5B-5C). As a reference for the maximum possible labelling index, we analysed specimens receiving Tamoxifen before the lesion (T-bQA; Figure 5A).

**Figure 5.**
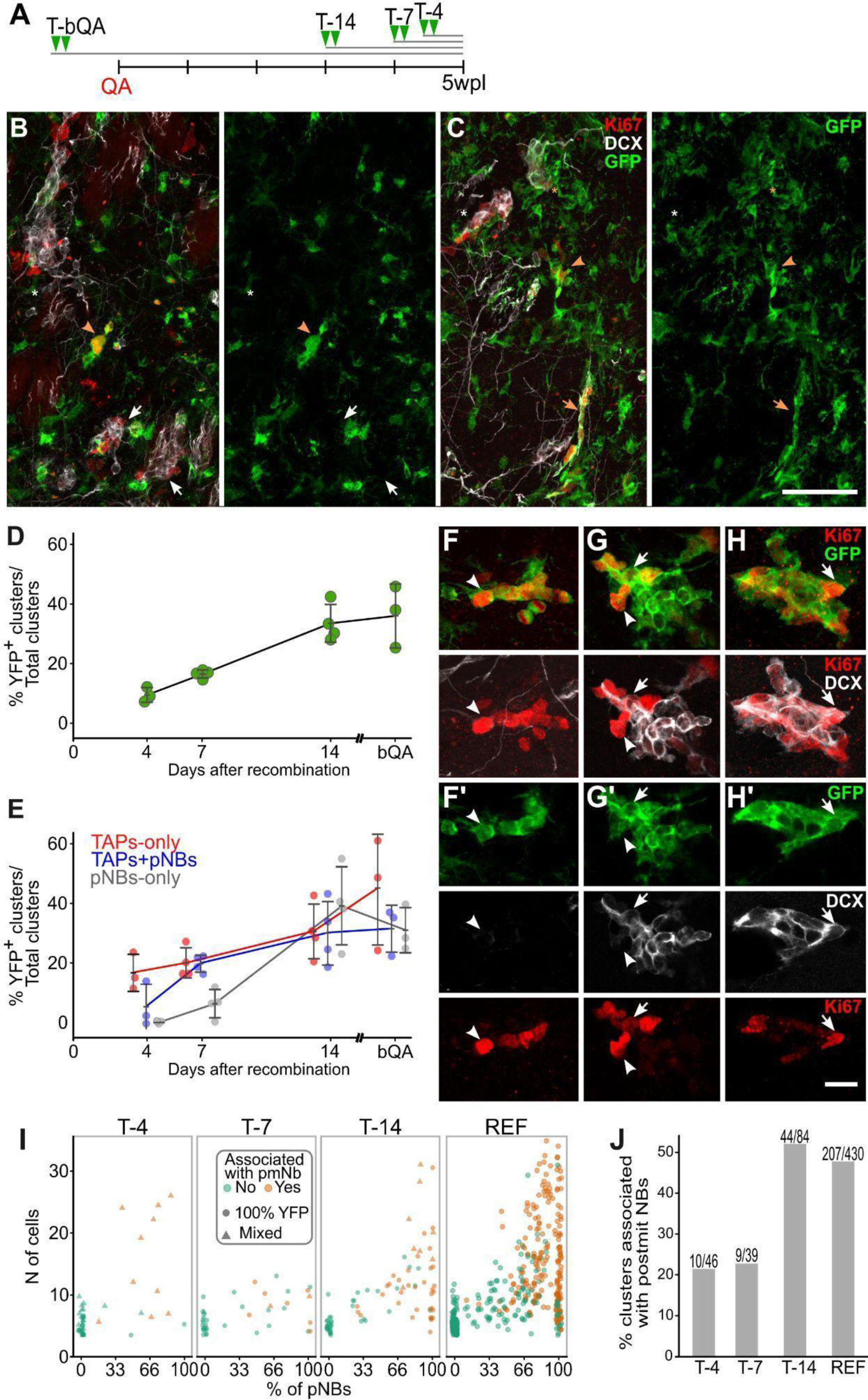
Constant astrocyte activation generates TAPs-only clusters that autonomously mature overtime. (A) Timeline of Tamoxifen (T) administration in Glast^CreERT2/+^xR26R-YFP mice. (B,C) YFP^+^ and YFP^-^ clusters in T-4 (B) and T-14 (C). Arrowheads: TAPs-only clusters; asterisks: TAPs+pNBs clusters; arrows: pNBs-only clusters. White and orange: YFP^-^ and YFP^+^ clusters, respectively. (D) Ki67^+^cluster labelling index measured as the percentage of clusters containing at least 1 YFP+ cell. (E) TAPs-only (red), TAPs+pNBs (blue) and pNBs-only (grey) clusters labelling index. (F-H) MAX projections of completely YFP^+^ TAPs-only (F), TAPs+pNBs (G) and pNBs-only (H) clusters. (F’-H’) Single focal planes of (F-H), respectively. Arrowheads: TAPs; arrows: pNBs. (I) Percentage of pNBs vs the total number of cells per cluster at different time-points (see also Figures S6A-S6B). Clusters associated with postmitotic NBs are in orange. REF as in Figure 2B. Circles: fully recombined clusters; triangles: mixed YFP^+^ and YFP^-^ clusters. (J) Fraction of clusters associated with postmitotic NBs at different time points and in the REF (see also Table S1). Data in (D) and (E) are shown as mean±SD at each timepoint (see also Table S1). Scale: (B-C) 50 µm; (F-H’) 15 µm.

The cluster labelling index linearly increased from 9.4±2.5% in T-4 to 33.5±6.3% in T-14 and remained stable in T-bQA (36.1±10.9%; Figure 5D; Table S1, *p<0.001; T-4 vs T-14: p=0.002; T-14 vs T-bQA: p=0.938*). This linear increase in the YFP+ cluster proportion means that astrocyte activation occurs in a staggered manner at a constant rate (see mathematical analysis below) until it reaches a plateau: around 14 days are needed for all the Ki67^+^clusters that are present at 5 wpl to have experienced at least one astrocyte activation event.

### Each astrocyte neurogenic activation event initiates a new Ki67^+^cluster that progressively matures

To explore if astrocyte activation preferentially occurs in specific phases of the cluster life we evaluated the labelling index of the main Ki67^+^clusters maturation profiles across time points (Figures 5E-5H’). The labelling index of TAPs-only clusters was already at the plateau at T-4 (Figures 5E-5F’; Table S1, p=0.359), showing that TAPs-only clusters are on average within four days from an astrocyte activation event. By contrast, only the 6±8% of TAPs+pNBs clusters contained YFP+ cells at T-4 and their labelling index significantly increased to 31±11% at T-14 (Figures 5E and 5G-5G’; Table S1, *p=0.006*), demonstrating that for this higher maturation profile, more time has passed since the last astrocytes activation. Finally, pNBs-only clusters almost never included cells derived from recent astrocyte activations, at T-4, and their YFP labelling index significantly increased between T-7 and T-14 (Figures 5E and 5H-5H’; Table S1, *p=0.001*), revealing that they are more distant in time from astrocyte activation than all other cluster types. In line with these observations, the number of Ki67^+^ cells, the fraction of pNBs as well as the association with post-mitotic NBs progressively increased with time (Figures 5I, S7A-S7B and 5J; Table S1, *p<0.001 in each case*) and became indistinguishable from the reference population by T-14 (Figures 5I, S7A-S7B and 5J; Table S1, T-14 vs REF: *p=0.249, p= 0.549 and p= 1, respectively*). Thus, striatal astrocyte activation predominantly occurs in the initial phases of cluster life, giving rise to TAPs-only clusters which then gradually mature. To further validate that astrocytes activate only at cluster initiation we examined the prevalence of YFP labelling in YFP^+^ clusters (Figure 6A). At T-14 and T-7 most YFP^+^ clusters were entirely YFP^+^, indicating that they originated from the clonal expansion of a single astrocyte (Figures 6A and S7A-S7B; Table S1). By contrast, at T-4 only 40% of the YFP^+^ clusters were 100% recombined (18/46; Figure 6A). These clusters were almost exclusively small TAPs-only clusters (Figure S7D). Interestingly, with the exception of 5 putative fused clusters (Figures S7C-S7C) most of the mixed clusters (including YFP^+^ and YFP^-^ cells) had only slightly more mature profiles suggesting they have a similar age (Figures 6B, S7A-S7B, S7E-S7G and 5I, Table S1). As most mixed clusters contained about 30-60% of YFP^+^ cells (Figure 6C), we can infer that Glast^CreERT2/+^ driven recombination occurred at 2-or 3-cell stage in astrocytes or their earliest progeny. Altogether these results indicate that astrocytes activate virtually exclusively to initiate a new cluster. As observed in Confetti mice, at all time-points after tamoxifen administration a Ki67^-^DCX^-^ dormant cell, in rare cases two, clustered with the 75% of YFP^+^ Ki67^+^clusters (Figures 6D-6E). The dormant cells turned out to be GFAP^+^ astrocytes in 26 out of 27 tested cases (Figure 6D) supporting that striatal astrocytes rapidly come back to quiescence after division and their reactivation is extremely rare, at least within established neurogenic foci.

**Figure 6.**
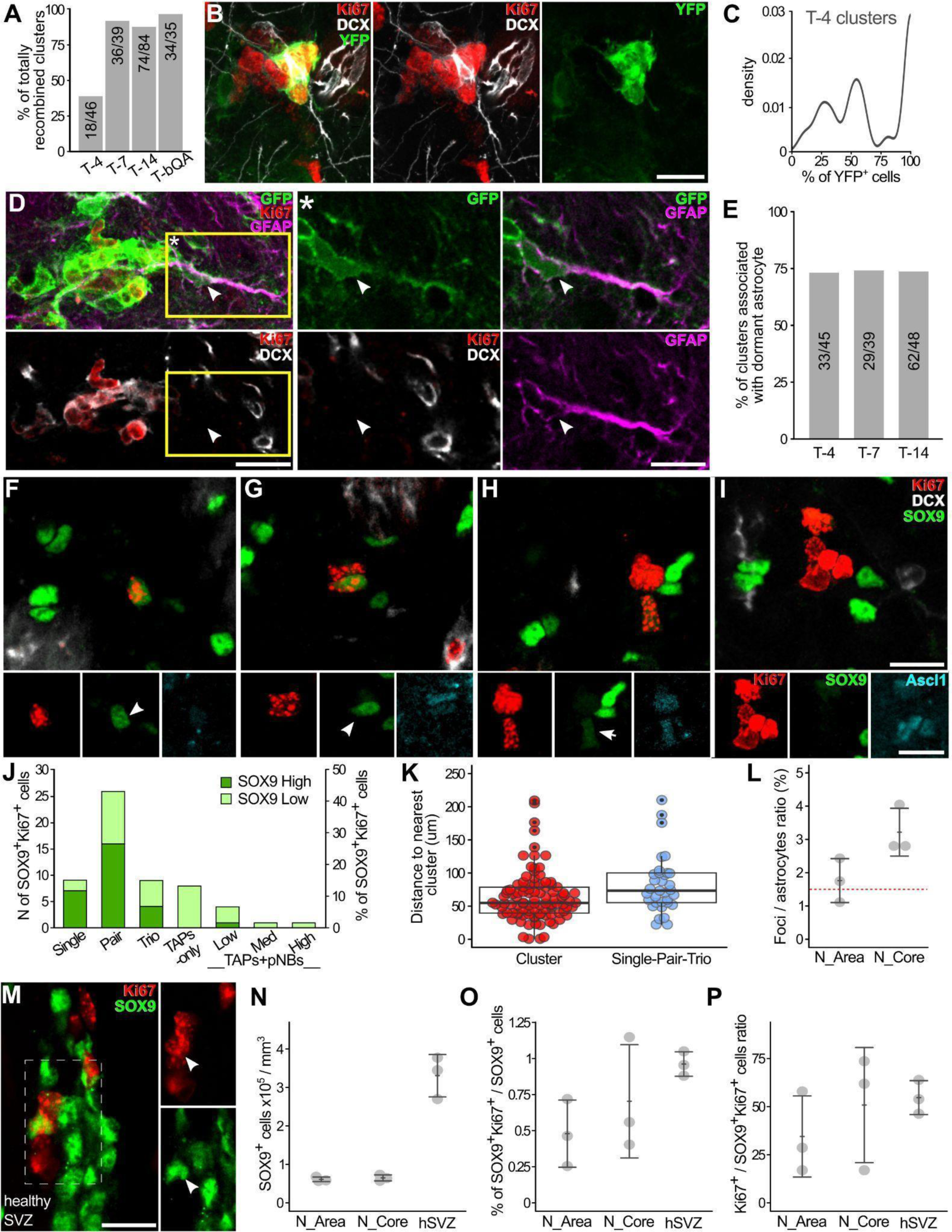
Astrocytes activate in cluster-free areas and shortly come back to quiescence. (A) Fraction of YFP^+^ clusters entirely composed of YFP^+^ cells over total YFP^+^ clusters. (B) Immature mixed cluster at T-4 (see also Figures S7C-S7G). (C) Relative distribution of the fraction of YFP^+^ cells per cluster in the T-4 YFP^+^ clusters. (D) YFP^+^ cluster containing a non-proliferating GFAP^+^ astrocyte. (*) single focal plane at a higher magnification of the yellow box in (D). (E) Percentage of Ki67^+^clusters containing at least one dormant Ki67^-^DCX^-^ cell overtime. (F-I) Single (F), pair (G), trio (H) or clustered (I) Ki67^+^ cells in which at least 1 proliferating cell is a SOX9^+^ astrocyte. The lower panels show SOX9 and Ascl1 expression in Ki67^+^ cells. Arrowheads: “SOX9^High^” cells; arrows “SOX9^Low^” cells. (J) Number (left) and percentage (right) of Ki67^+^SOX9^High^ and SOX9^Low^ cells in Ki67^+^ groups (see also Figure S7H). (K) Comparison of the distance to the nearest cluster between clusters (red) and single/pair/trio (grey) (see also Figure S7I; Table S1). (L) Foci/astrocytes ratio (%) in the neurogenic area (N_Area) and in its core (N_Core) (see also Figure S7I). Red dashed line: astrocytes activation rate in the SVZ^41^. (M) A SOX9^High^Ki67^+^ cell (arrowhead) in the SVZ of a healthy mouse. Left panel: maximum intensity projection. Right panel: single focal plane of the dashed rectangle. (N-P) Comparison of the astrocyte density (N), the % of proliferating astrocyte (O) and the ratio between TAPs/pNBs and proliferating astrocytes, among striatal neurogenic areas and healthy SVZ (see also Table S1). Scale: (B), (D),(F-I) and (M) 15 µm.

A mathematical analysis of cluster initiation dynamics (Supplementary Text 1, section 1) shows that the initial slope of the increase of the cluster labelling index over time corresponds to the cluster initiation rate (Figure 5D), which also applies to the astrocyte activation rate. Given that this slope is approximately linear, we can conclude that the astrocyte initiation rate is approximately constant. When approaching the cluster lifetime, the cluster labelling index is expected to saturate to a plateau (Supplementary Text 1, section 1). This occurs at around 10-14 days, consistent with the previously estimated cluster lifetime of 10.6 days. These results thus conclusively demonstrate that clusters undergo steady-state turnover and provide direct proof that astrocytes initiate new clusters that progressively mature before undergoing exhaustion.

### Astrocytes activate in neurogenic foci-free areas

The above data predict that astrocytes activate in cluster-free areas. To directly verify this possibility, we analysed the organisation and distribution of proliferating astrocytes in the neurogenic area in pairs of subsequent sections stained for Ki67, DCX, the early neuronal commitment marker Ascl1 and the astrocyte marker SOX9 at 5wpl (Figures 6F-6I and S2G-S2G’). We distinguished between cells expressing SOX9 at similar or slightly lower levels as the non-proliferating astrocytes (SOX9^High^) from those showing substantially lower levels (SOX9^Low^; Figures 6F-6I) that may represent TAPs^36^. Most Ki67^+^SOX9^+^ cells were in groups smaller than four cells with a strong prevalence for pairs (Figures 6G and 6J). The fraction of SOX9^High^ cells among Ki67^+^SOX9^+^ cells progressively decreased from 78% in individual cells to 62% in pairs, 44% in trios and 7% in Ki67^+^clusters (Figures 6F-6I). The decrease in Sox9 expression was paralleled by an increase in Ascl1 expression (Figure S7H; Table S1, *p<0.001* in both cases) confirming that early neurogenic foci cells lose their astrocytic identity before reaching the 4-cell stage.

Interestingly, the groups of less than four Ki67^+^ cells containing activated astrocytes never contacted postmitotic NBs and their nearest Ki67^+^cluster was on average farther away than the mean nearest-neighbour distance between Ki67^+^clusters (Figures 6K and S7I; Table S1, *p=0.013*). These results show that striatal astrocytes preferentially activate in areas where there were no pre-existing clusters and also far from previous activation events.

### Striatal and SVZ astrocytes activates at similar rates

Neurogenic foci represent activation events that occurred over the last 10 days. To evaluate striatal astrocytes activation rate we thus evaluated the ratio between their number and that of all astrocytes in the neurogenic area or its core, respectively comprising 95% or 25% of the foci. The astrocyte density did not differ between these areas and although it tended to be higher than in the rest of the striatum (Figure S7I) it was still about 5 times lower than in the SVZ of healthy mice (“hSVZ”; Figures 6M-6N; Table S1, N_Area vs hSVZ: p= 0.0001, N_Area vs hSVZ: p= 0.0001). The foci/astrocytes ratio ranges between 1% and 4%, corresponding on average to 0.18% activations per day in the neurogenic area, a value that is strikingly similar to that calculated for SVZ astrocytes (0.15% activations per day) (Figure 6L)^41^. In support of this observation, the fraction of proliferating astrocytes (Ki67^+^Sox9^+^ /Sox9+ cells) as well as the ratio between TAPs/pNBs and the latter cells in SVZ and striatum were also similar (Figures 6O-6P, Table S1, *p= 0.168*, *p= 0.52)*. These results indicate that striatal astrocytes share with SVZ astrocytes not only the activity pattern but also the activation rate, implying a comparable prevalence of NSC potential.

### Cell fate choice timings in the turnover of neurogenic foci

As astrocytes divide mainly once per cluster, most of the neurogenic production is due to intermediate progenitor divisions. The autonomous growth and steady-state turnover of striatal neurogenic clones allowed us to exploit our rich collection of reconstructed clusters (Figure 2) to model TAPs and NBs expansion and fate choices. Intermediate progenitors are usually thought to expand stereotypically, however their behaviour is still poorly characterised^42^. We implemented a stochastic mathematical model^43^ (Supplementary Text 1) to discern (1) how the differentiation propensity (TAP→pNB; pNB→NB transitions) varies over time and whether it depends on cluster ages. (2) Whether the differentiation of TAPs and NBs is coupled to cell division via asymmetric and symmetric divisions, or whether it can occur independently of cell divisions.

From the comparison of model predictions with the cluster size distribution (Supplementary Text 1, section 2), we could clearly discard models in which TAPs differentiation occurs synchronously after a fixed number of cell divisions or after a fixed amount of time past astrocyte activation (Supplementary Text 1, section 4.1.1). Conversely, we found an excellent fit for the distribution of TAPs per cluster when allowing cell fate choices to be stochastic, yet only if the differentiation propensity increases in an accelerated way over time (Figures 7A-7B; Supplementary Text 1, section 4.1.3). The best fit is achieved if we assume that differentiation can occur independently of cell division, which is thus the most likely scenario. We can further exclude any significant contribution from asymmetric divisions of NBs although we cannot strictly exclude the option that differentiation is coupled to symmetric cell division (Supplementary Text 1, sections 4.1.2 and 4.1.3).

**Figure 7.**
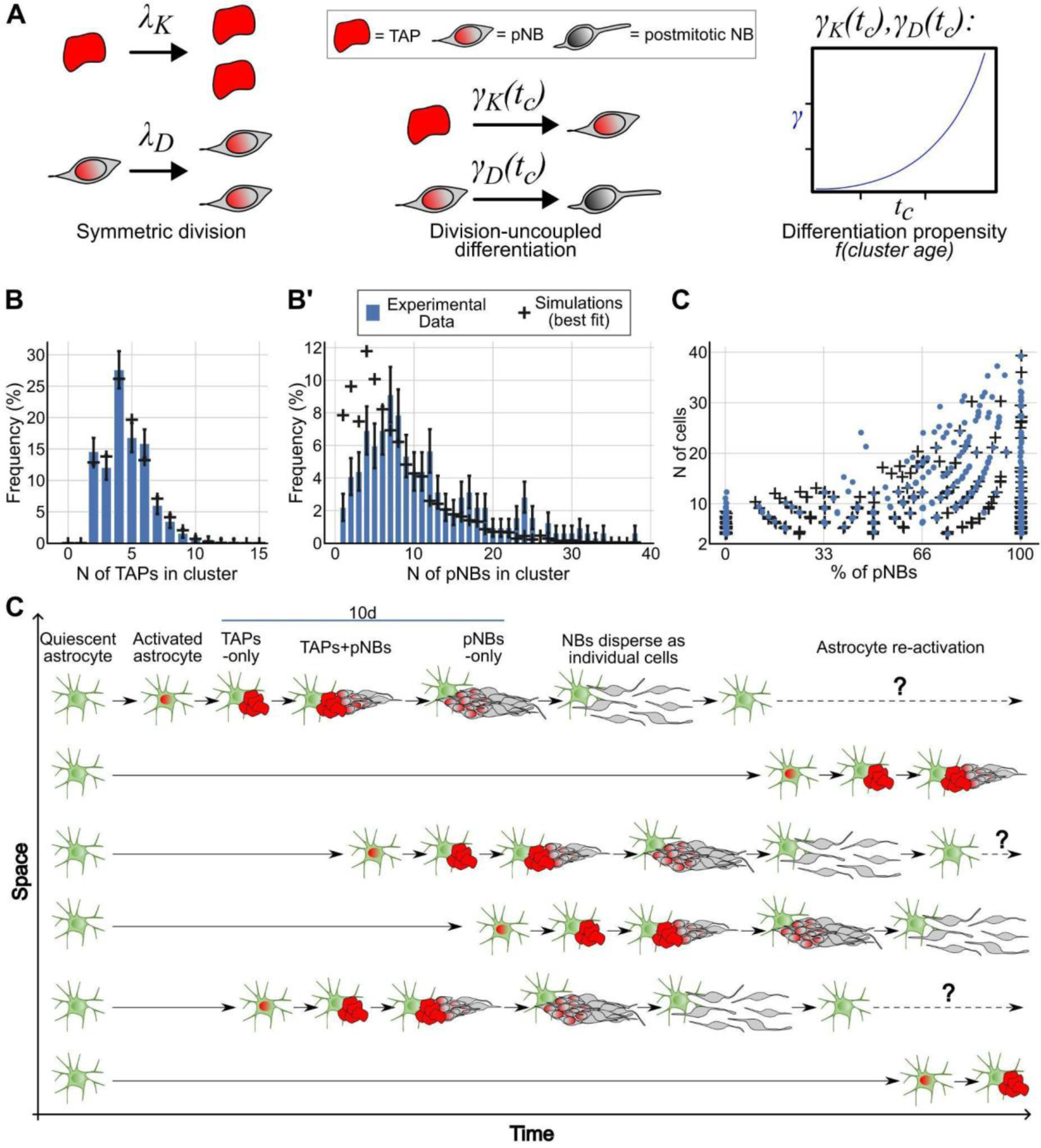
Mathematical modelling of cell fate choices in Ki67^+^clusters. (A) Schematics of the best fitting model features. TAPs and pNBs mostly undergo symmetric duplicating division with *λ_K_* and *λ_D_* rates, respectively. The differentiation propensities of TAPs and pNBs (*γ_K_* and *γ_D_*, respectively) change as a power law function over the cluster age (see also Data Supplementary text 1). (B,B’) Best model fitting of TAPs (B) and pNBs (B’) frequency distributions in clusters. Bars are measured frequencies, black crosses represent simulation output, obtained from the best fitting model (see also Data Supplementary text 1). (C) Scatterplot showing the percentage of pNBs vs number of cells per cluster. Similar to Figure 2B, with the addition of simulated clusters obtained from the best fitting model (data not used for fitting the model). (D) General scheme of the spatiotemporal dynamics of striatal astrocytes activation and lineage progression, that ultimately result in a continuous turnover of neurogenic foci.

The modelling of pNBs to NBs transition yielded very similar results. As for TAPs, only if the differentiation propensity increases over time in an accelerated way, do we obtain a reasonable model fit (Figures 7A and 7B’; Supplementary Text 1, section 4.2). However, for pNBs we can exclude that differentiation is coupled to cell division; thus, the exit of the cell cycle occurs independent of the previous division. We note that some deviations between data and model predictions remain even for the best fit, for very small and very large cell numbers, respectively over and underrepresented in the model (Figure 7B’). This is likely due to the dispersion of the last pNBs within clustered NB at cluster end (very small) and cluster fusions (very large).

Neurogenic foci cells thus show features of both stochastic and deterministic fate choices that ultimately result in their invariant exit from the cell cycle but with consistent variability. Hence, these data fully validate the presented model of neurogenic foci initiation, maturation and dynamic turnover (Figures 7C-7D) and mechanistically confirm the intermediate progenitor nature of the Ki67^+^cluster cells.

## DISCUSSION

Here, we demonstrated that the spatiotemporal patterns of striatal astrocyte neurogenic activity closely resemble those of classic adult vertebrate brain niches. However, these patterns emerged within the 3D lattice of parenchymal astrocytes, distinguishing them from conventional NSC monolayers^4^. Striatal astrocytes were mostly quiescent and sporadically activated, generating transient progenitors that autonomously expanded. Within the neurogenic core, astrocyte activation events were constant in time and random in space resulting in a steady state of continuous and widespread neuronal production. Although a large part of the striatal parenchyma had the potential to host neurogenic activation events, their frequency was contingent on the lesion border with a preference for the medial striatum functional domain. In this region, the rate of activity of striatal astrocytes was comparable to that of SVZ astrocytes. The mouse striatum can thus be considered as a dormant neurogenic niche hosting a widespread population of qNSC that can be focally activated.

### NSCs activation dynamics in the striatal neurogenic niche

Spatiotemporal dynamics of an entire NSCs population have so far been determined only in zebrafish through live imaging^11,44,45^, a technique that however is not currently available for such deep and large 3D volumes as the striatal parenchyma^46^. Nonetheless, an advantage of the striatal neurogenic system emerged from the local expansion and minimum overlap of astrocytes proliferating progeny. This allowed us to use neurogenic foci as a proxy for the location of astrocyte activation events over the last ∼10 days. By integrating two complementary dynamic approaches, we unveiled that astrocytes neurogenically activate at a constant rate. In parallel, the neurogenic foci spatial independence revealed that activation events are randomly distributed. Moreover, these events preferentially occur in new locations, far from pre-existing foci, overall demonstrating the widespread presence of neurogenically competent astrocytes within the striatum. A similar random pattern of NSC activation inhibited by previous events was described in zebrafish^44^. In that case, inhibition was proposed to be mediated by NBs, which inhibit NSC activation also in mammalian niches through GABA release or cell-cell contacts^44,47,48^. Supporting this possibility, proliferating striatal astrocytes were virtually never in direct contact with NBs.

Following activation, in both SVZ and DG, NSCs undergo consuming neurogenic divisions with high probability^2,41,49^, while asymmetric self-renewing divisions characterise neurogenesis during ageing^50–53^. This latter mode of division is preferred also by striatal astrocytes, ensuring maintenance of the population. Of note, a recent study suggests that in the SVZ NSCs divide always symmetrically and the fate of daughter cells is regulated locally by competition for the occupancy of “a restricted niche”^8^. According to this model, the lower astrocyte density in the parenchyma may increase the probability that at least one cell will return to occupy the niche. The division of TAPs and NBs were regulated by stochastic processes although an accelerating differentiation propensity introduces a deterministic factor constraining their expansion. Similar stochastic behaviour of TAPs has been described in the DG by *in vivo* imaging^6,42,49^. As intermediate progenitors can respond to physiologic and pathologic stimuli^54,55^, it would be interesting to explore if extrinsic cues contribute to their varied local behaviour in the striatum.

### Neurogenic permissiveness of the striatal parenchyma

Our results demonstrated that the striatum acts as a neurogenic niche by constitutively hosting a widespread population of dormant NSCs. According with our results, all major known NSCs quiescence stimulating pathways such as Notch^56,57^, BMP^58,59^, β1 integrin^60,61^, S1P^62^ or GABA^47,63^ are active in parenchymal astrocytes^58,64–67^. Unlike canonical niches, in the parenchyma inhibition of these pathways alone fails to induce neurogenic activation^58,64,65^. Nonetheless, at least for Notch inhibition, single cell RNAseq indicates induction of a primed NSC state^25,65^. Acute lesions can also reduce NSCs quiescence in both canonical niches^68,69^, and parenchyma^21,22,26,27,70^ through loss of contact with neurons^27^ and/or inflammatory stimuli^16,71,72^. Only in some cases however, these state transitions progress to neurogenic activation *in vivo*, suggesting that additional signals are needed to regulate this switch. Few Notch-inhibited astrocytes became neurogenically active in the medial striatum, leading to the hypothesis that the nearby SVZ may be a source of such signals^27,65^. QA-induced neurogenic activation events also show a bias for the medial striatum, however their pattern was not shaped as an SVZ centred gradient but peaked more laterally. This suggests the involvement of local signals preferentially associated with the lesion border and the medial striatum. The nature of these stimuli remains to be established. Nonetheless, similar neurogenic foci in normal rabbits^32^, under progressive degeneration^33^ and stroke^27^, were also preferentially distributed in the medial striatum, indicating that progenitors residing in this functional domain can activate in multiple contexts, including physiological conditions. As after stroke^27^, the QA-induced neurogenic response shows delayed activation, during the third week after lesion, which may be consistent with the tissue remodelling phase^73,74^. In all models of striatal neurogenesis neuroblasts live transiently and do not replace lost striatal neurons^23,32,33,69^. The identity and function of these cells are still unknown, but depleting or silencing similar neuroblasts worsens cortical stroke recovery^75^. These observations introduce the intriguing possibility that striatal neuroblasts support a conserved form of plasticity, frequently associated with the medial striatum, that contributes to the remodelling of damaged striatal circuits.

### How widespread is the neurogenic potential among astrocytes?

Notch pathway blockage causes widespread transition of cortical and striatal astrocytes to acquire the transcriptional signature of primed NSC-like state^25,65,76^, leading the authors to speculate that all astrocytes possess a neurogenic potential. Here, we demonstrate that naive striatal astrocytes do have such widespread potential. Establishing whether all striatal astrocytes harbour NSC potential is challenging, a question that is unresolved even for the canonical NSC^2,4^. The close similarity in both the pattern and rate of neurogenic activation indicate that these two populations have comparable prevalence of NSCs^41^. As for canonical niches,the exact fraction of astrocytes that is recruited to striatal neurogenesis cannot be estimated precisely because we ignore their reactivation rate. Here we assessed that astrocytes do not reactivate at least over the Ki67^+^cluster lifetime (Figure S7D), but their quiescence likely lasts longer because: (i) clusters of postmitotic NBs remain for some time after proliferative pool exhaustion and (ii) astrocytes activate far from pre-existing clusters. Moreover, the local activation density may be much higher than average, as suggested by cases of cluster fusion. Thus, the maximum density of recruited astrocytes, a proxy of the neurogenic potential per unit area, is likely much higher than our measure over 10 days (up to 1/25 astrocytes), potentially including all astrocytes. In non-mammalian vertebrates, radial glia persist into adulthood where they act both as NSC and in supporting neuron homeostasis and function^77–79^. In mammals, it was thought that these two functions were fulfilled by distinct cell types and that astrocytic differentiation implies a permanent loss of NSC potential. However, our results definitely demonstrate that this potential can be maintained in parenchymal astrocytes. Whether all astrocyte subtypes/states have the same probability to activate a NSC potential, both between and within regions, remains to be established. NSCs are a heterogeneous population of progenitors committed to generate specific neuron types and, at least in adults, are regulated by distinct conditions and neuronal circuits^80–85^. Within the mammalian parenchyma, astrocytes, while specialising in local circuit-specific control, may have also acquired a circuit-specific regulation of their neurogenic capacity. In this perspective, as for canonical niches, network activity represents a major candidate regulator of the parenchymal neurogenic potential.

We already showed activation of a dormant parenchymal niche in the ventral striatum around weaning in guinea pigs, but similarly to the dorsal SVZ this niche was confined to the narrow pallial-subpallial-boundary^23^. Our new study extends the complexity of adult NSC spatial heterogeneity to the 3D lattice of parenchymal astrocytes. This radically challenges a traditional view of neurogenic niches as unique domains for stem cell maintenance acting as barriers to parenchymal differentiation signals. The brain may rather have a widespread quiescent neurogenic potential organised as a mosaic of progenitor domains regulated by specific activation signals. In canonical niches these signals are constitutively active, while in other regions they may be triggered only in specific conditions. The size, distribution, regulation and cell fate potential of these progenitor domains await further analysis, thereby paving the way for a deeper understanding of the full adult brain neurogenic potential.

### Limitations

We could not precisely measure how many astrocytes harbours a NSC potential, yet distinguishing between reversible cell state transitions and stable intrinsic identities is a major unsolved issue in astrocytes and NSC biology. In addition, as the lateral striatum was always more severely damaged, the lower density of neurogenic activation events in that area may represent a peculiar feature of our lesioning paradigm.

## Supporting information

Supplementary Text 1 - Mathematical modeling

Table S1 - Statistical analyses

## ACKNOWLEDGMENTS

The authors would like to thank Sara Trova and Alessia Caramello for preliminary work, Gregorio Valsania for help in 3D reconstructions, Valentina Cerrato, Stefano Zucca and Ilaria Bertocchi for critically reviewing the manuscript. This research was supported by: Fondazione Cecilia Gilardi, Fondazione Umberto Veronesi, funding from PNRR MUR – M4C2 – Investimento 1.4 to PP, the European Union’s Horizon 2020 research and innovation programme under grant agreement No 874758 to AB and funds of the University of Turin and Compagnia di San Paolo (S1618 Grant) to AB and FL. This study was also supported by the Italian Ministero dell’Istruzione, dell’Università e della Ricerca—MIUR project “Dipartimenti di Eccellenza 2018–2022 and 2023-2027 to Dept. of Neuroscience “Rita Levi Montalcini”.

## AUTHOR CONTRIBUTIONS

Conceptualization, M.F., G.N. and F.L.; Software - 3D reconstructions, F.L.; Investigation, M.F. and G.N.; Formal Analysis, M.F, G.N. and F.L.; Spatial Analyses, M.F.; Mathematical Modelling, P.G.; Formal Analyses - BrdU Time course, J.P.; Writing - Original Draft, M.F., G.N., P.G. and F.L.; Writing - Review & Editing, M.F., G.N., P.G., P.P., A.B. and F.L.; Visualization, M.F, G.N. and F.L.; Supervision, P.P., A.B. and F.L.; Funding Acquisition, P.P., A.B. and F.L.

## SUPPLEMENTARY ITEMS TITLES AND LEGENDS

**Supplementary Figures.**

**Supplementary Text 1. Detailed description of mathematical modelling campaign.**

**Table S1. Results of all statistical analyses presented in the study.**

**Video S1. 3D reconstruction of seven striata at 5wpl, related to Figures 1 and S1**

The 528 focal planes of the 40 sequential sections 3D reconstructed from specimen #N1 are shown from the most anterior to the posterior, and then are presented all together as a MAX projection. After a 360° rotation of the whole reconstruction, a couple of isolated Ki67^+^clusters are shown. Ki67 staining is presented alone and the striatal cluster segmentation (magenta) is overlaid. Note the sparse Ki67^+^ cells that are excluded from the segmentation as they may represent new astrocyte activation events or other proliferating cell types. Clustered Ki67^+^ cells in the SVZ are shown in blue. By min 1:14 blender renderings of surfaces and manually annotated Ki67^+^clusters, represented as balls, are shown. A measure of relative Ki67^+^cluster density is shown as an object of different colours, with the hottest representing the highest value. The Ki67^+^clusters distribute along the lesion (purple) border, mainly at its rostro-medial border, the ventricle is in blue and the striatum is transparent. From 1:25 onward each of the seven reconstructed specimens are shown one at a time with their lesion in purple, while their Ki67^+^clusters and cluster densities are shown with the same colours used in Figures 1 and S1. Specimens are observed from the rostral part rotating alternatively from medial to lateral and lateral to medial as shown by the reference striatum in the inset. Specimens are ordered from the one having the most posterior neurogenic response to the one displaying the most anterior one. From 1:53 all specimens are overlaid together highlighting that, on the whole, Ki67^+^clusters cover most of the medial and rostral portion of the striatum. In both Imaris and Blender, the 3D views are in orthographic mode.

**Video S2. High resolution 3D reconstruction of a portion of the neurogenic area at 5wpl, related to Figure 2 and S2**

A portion of the neurogenic area of specimen #G14.3 is shown in this video. SVZ Ki67^+^ cell segmentation is shown in blue, while striatal Ki67^+^cluster segmentation is shown in random colours to highlight that each of them is an independent structure. Starting from 24 seconds the random colour segmentations are shown as transparent objects allowing to appreciate individual Ki67^+^ and/ or Ki67^+^DCX^+^ cells within each structure. Ki67^+^clusters at different maturation stages are zoomed in at different timepoints: a TAPs-only cluster at 10 seconds, a TAPs+pNBs cluster at 25 seconds; a pNBs-only cluster at 35 seconds.

## EXPERIMENTAL PROCEDURES

### Resource availability

#### Lead contact

Further information and requests for resources and reagents should be directed to and will be fulfilled by the lead contact, Federico Luzzati.

#### Materials availability

This study did not generate any unique reagents.

#### Data and code availability

Primary data are available from the lead contact upon request.

Codes for performing 3D reconstructions are available at: https://drive.google.com/drive/folders/1MHtB4hBN5qYblCSePC5VvXo5FfCUQrxt?us p=drive_link.

3D reconstructions of individual specimens are available at: (we will activate a link if the paper will be accepted).

Any additional information required to reanalyse the data reported in this paper is available from the lead contact upon request.

### Experimental model and subject details

#### Animal procedures

The experimental plan was designed according to the guidelines of the NIH, the European Communities Council (2010/63/EU), and the Italian Law for Care and Use of Experimental Animals (DL26/2014). It was also approved by the Italian Ministry of Health (authorization 327/2020-PR) and the Bioethical Committee of the University of Turin. The study was conducted according to the Arrive guidelines.

#### Mouse lines

Experiments were performed on 8-12 weeks old animals of the following mouse lines: C57BL/6J mice (Harlan laboratories; n= 18 males), Glast^CreERT239^, R26R-YFP^86^ and R26R-Confetti^40^. Glast^CreERT2^ mice were crossed to R26R-YFP or R26R-Confetti mice to produce mice heterozygous for Glast^CreERT2^ and homozygous for R26R-YFP (Glast^CreERT2/+^xR26R-YFP, n = 11 males and 3 females) or heterozygous for R26R-Confetti (Glast^CreERT2/+^xR26R-Confetti, n= 4 males).

### Method details

#### Stereotaxic injections

Mice were anesthetised with 0.3 ml/kg ketamine (Ketavet, Gellini) and 0.2 ml/kg xylazine (Rompun, Bayer), positioned in a stereotaxic apparatus (Stoelting) and injected with a pneumatic pressure injection apparatus (Picospritzer II, General Valve Corporation). Injection coordinates: quinolinic acid (1 µl; diluted to 120 mM in 0.1 M PB), +0.1 mm AP, −2.1 mm ML and −2.6 mm DV.

#### BrdU pulse labelling

C57BL/6J lesioned mice received two intraperitoneal injections of 5-bromo-2-deoxyuridine (BrdU, Sigma-Aldrich; 50 mg/kg in 0.1 M Tris pH 7.4) 6 hours apart and were sacrificed either 2h (B-8h) or 4d (B-4d) after the last injection. Animals of the B-8h group were sacrificed at 5wpl while B-4d at 3, 4, 5 or 8wpl.

#### Tamoxifen administration

Tamoxifen (Sigma-Aldrich) was dissolved in corn oil (Sigma-Aldrich) and was administered by forced feeding (oral gavage) at a dose of 2.5mg per administration with a 24-hour interval. Glast^CreERT2/+^xR26R-YFP mice received two tamoxifen administrations, while Glast^CreERT2/+^xR26R-Confetti mice received five administrations.

#### Histology

Animals were anesthetised with a ketamine (100 mg/kg) and xylazine (33 mg/kg) solution and perfused with a solution of 4% paraformaldehyde (PFA, Sigma-Aldrich) and 2% picric acid (AnalytiCals) in 0.1 M sodium phosphate buffer (PB) pH 7.4. Brains were then post-fixed for 5 hours, cryoprotected in 30% sucrose (Sigma-Aldrich) in 0.1M PB pH 7.4, embedded at −80°C in Killik/OCT (Bio-Optica), and cryostat sectioned in series of 50 µm-thick coronal sections.

#### Immunofluorescence staining

Sections were incubated for 48 h at 4°C in 0.01 M PBS pH 7.4 containing 2% Triton X-100, 1:100 normal donkey serum and primary antibodies (see Key resources table). For BrdU staining, sections were pre-incubated in 2M HCl for 30 min at 37°C and then rinsed in 0.1 M borate buffer pH 8.5. Sections were incubated overnight with appropriate secondary antibodies (see Key resources table) and coverslipped with antifade mounting medium Mowiol (4-88 reagent, Calbiochem). For Mash1 staining, sections were pre-incubated for 5 min at 95°C in an antigen retrieval solution consisting of Citrate buffer (20mM citric acid, 0.05% Tween, pH=6).

One of the three series of the Confetti sections was stained for five antigens: Ki67, DCX, GFAP, GFP and RFP. Since the anti-Ki67 and anti-RFP antibodies were both raised in rabbits, slices were initially stained only for anti-Ki67, anti-DCX, anti-GFP, and anti-GFAP primary antibodies. Subsequently, sections were incubated with anti-rabbit Monovalent Fab Fragments (see Key resources table) to saturate the binding sites recognized by the anti-rabbit antibody. Finally, the sections were treated with rabbit anti-RFP antibodies followed by anti-rabbit secondary antibodies.

#### Confocal imaging

All the fluorescent images presented in this study were acquired on a Leica SP5 confocal microscope (Leica Microsystems) equipped with 40x and 63x objectives (HCX PL APO lambda blue: 40x, NA 1.25; 63x, NA 1.4). For the whole striata 3D reconstructions (Figures 1, S1 and 4) images were acquired at a resolution of 0.76 µm/pixel with an optical sectioning in Z every 2.5 µm. For all the quantifications of Ki67^+^cluster number and cellular composition, as well as for the quantification of striatal astrocytes recombination efficiency in the Confetti experiment, images were taken at a resolution of 0.38 µm/pixel with optical sectioning in Z every 1.5 µm. For the quantifications of Confetti cluster cellular composition and of Gfap expression in dormant cells, images were acquired with the 63x objective at a resolution of 0.24 µm/pixel with optical sectioning in Z every 1 µm. For acquisition tiles, stitching was done automatically by the LasAF software after imaging.

#### Epifluorescence imaging

Whole-section images, used as anatomical references for the high resolution 3D reconstructions (see below), were acquired automatically using a slide scanner Axio Scan (.Z1, Zeiss) equipped with a 10x objective (Plan-Apochromat 10x, NA 0.45 M27). Images of a single focal plane per section were acquired at a resolution of 0.65 µm/pixel.

#### 3D reconstructions

Confocal microscopy serial section 3D reconstructions were performed by modifying a previous method^87^. All the following image processing and alignment procedures were performed with Fiji using a set of original scripts/custom-made scripts available at the following link: (We will activate a link if the paper is accepted). The 3D reconstructions of all the specimens will be available at the following link: (We will activate a link if the paper will be accepted). Whole-section epifluorescence acquisitions (Axio Scan) or cryotomography images served as a backbone anatomical reference for the 3D reconstructions of high resolution confocal acquisitions.

##### Low resolution whole sections registration

Epifluorescence images were extracted from the original Carl Zeiss Image (.czi) file format using the script *“Export tif from czi.ijm”*. This script allows saving as .tif files the images of each section splitted by channel, assigning a prefix with the channel name and the desired LUTs. Additionally, it includes a “zN” tag codifying the position of each section relative to the entire brain, allowing a straightforward image alignment (see below).

Epifluorescence images were then imported in the Fiji plugin TrakEM2^88^ using the script *“Import patches z names.py”* which imports the images into the appropriate layer based on their “zN” tag. The script also scales each image based on its pixel size to reach a final resolution of 1 µm/pixel. Thus, all the images that are imported with this script can be easily overlaid even though they were acquired at different resolutions. Sequential sections were manually aligned with a landmark based registration method to obtain a registration of all the sections that will work as the anatomical reference of the 3D reconstruction; we only used rigid transformations, i.e. translation and rotation. To ease this initial alignment phase, in some specimens, we performed cryotomography imaging during the cryostat cutting procedure. Briefly, images were acquired with a customised system composed of a Raspberry Pi camera (V2 8MP) with additional 16mm M12 objective mounted through a customised support on the cryostat anti-roll glass, which was controlled through the script *“CRIOpi Capture.py”*. Images were then calibrated and processed using the script *“CRIOpi Processing.ijm”* and imported as the first images in the TrakEM2 project using the same importing script. In those cases, epifluorescence images were simply overlaid with the cryotomography images and minimal adjustments to the overall image alignment were needed.

##### Confocal high-resolution images

The multi-channel z-stacks confocal acquisitions of the striatum (“z-stacks”) were named including the same “zN” tags of corresponding epifluorescence/cryotomography images. The z-stacks were initially batch-converted from the Leica image file (.lif) format into .tif files with the script *“Export tif from lif.ijm”*. Then, the z-stacks of each specimen were processed separately using the script *“Split tif to MAX and Sequence.ijm”* to obtain: i) the maximal intensity projections (“MAX projections”) of each channel and section; ii) the sequence of the single focal planes for each channel and section (“sequences”); iii) a .csv file containing the information for importing the sequences (see below). Similarly to what is done with the epifluorescence images, the script allows to apply the desired LUT to each channel and adds a prefix with the marker name (e.g. “Ki67^-^”).

Then, the MAX projections of each confocal section were imported using the same importing script and manually overlaid to the corresponding epifluorescence images. Finally, the sequences of each section were imported at the right positions with the “import from text file” TrakEM2 function, using the .csv file previously generated by the script “*Split tif to MAX and Sequence.ijm”*. As a last step, the same transformation previously applied on each MAX projection was applied to its corresponding sequences to register all the z-stacks.

For each layer in TrakEM2 are now available separate image files for each channel that could be visualised as a merged image using the script *“Set patch composite mode.py”*. For inspection and manual counting, the script *“Set visibility toggle channels.py’’* enables the use of customizable keyboard shortcuts for changing the visibility of each channel.

##### Annotations, region drawing and calibration

The Ki67^+^clusters were manually annotated as TrakEM2 “ball objects”. The areas of the dorsal striatum were manually drawn according to the Allen Brain Atlas subdivisions. The lesioned areas were manually drawn based on Gfap expression. To correct for the section shrinking along the z-axis and adjust the voxel size accordingly, the z-spacing of each focal plane was calculated by dividing the total depth of the reconstructed volume, according to cryostat sectioning, by the total number of focal planes. Of note, this is a fundamental requirement for obtaining proper distance measurements and performing 3D spatial analyses.

##### 3D models alignment and visualisation in Blender

Surface areas drawn in TrakEM2 of the striatum, lesioned area and lateral ventricle, were exported as .obj files from TrakEM2 using the script *“Export Arealists.py”* and imported in Blender (v2.79) with the script “*Blender_Import Surfaces and Reference points.py*”. Cluster coordinates were exported using the script *“Export Balls.py”* or *“Export Balls Confetti.py”* and imported in Blender as spheres using the script “*Blender_Import_Coordinates_as_Balls.py*”. Each specimen was imported in a different layer, after assigning materials, renderings were performed exclusively as orthographic views. All surfaces have been smoothed with a *smooth* modifier and additional imperfections were manually smoothed in sculpt mode. Reconstructions of the striatal projection domains of the anterior cingulate cortex (ACA) and the primary somatosensory areas(SSp) were obtained by aligning and manually drawing in TrakEM2 the segmentations of anterograde tracers injections obtained from (http://www.mouseconnectome.org/CorticalMap/page/map/5)^30^ and overlaid to the Allen reference atlas CCF V3. Reference samples were the SW110323-03 and SW110321-04 for ACA and SW120525-02, SW110418-01, SW110516-02, SW110418-02, SW110419-02 for SSp. Surfaces were imported in Blender as described. The registration of the different specimens to a common reference space has been performed manually based on the overall 3D shape of the striatum.

##### Imaris visualisation

Video S1, Video S2, Figures 1B-1B’’’ and 2F-2F’ were prepared in Imaris by importing the Ki67 and DCX channels of aligned #N1 and #G14.3 specimens in Imaris (v9.7.2). After adjusting brightness and contrast, the SVZ was manually drawn in order to exclude the fluorescence in that area and consider only the striatal neurogenic system. Ki67^+^clusters in Video S1 and Video S2, Figures 1B’-1B’’’’ and 2F’ were segmented semi-automatically based on fluorescence intensity, object size and striatal location. The video editing of Video S1 has been performed in Blender Video Editor 3.9.

#### Data visualisation

All the graphs were generated with ggplot289, except for the G function analyses (Figures 2G and S3A-B) whose plots were produced with a dedicated function of the spatstat package38.

The relative distributions reported in Figures 1F-1I, 7C and S4D-S4F actually represent the kernel density estimates of such distributions: the area under each curve is 1 and represents 100% of the observations. Although the term “relative” is not completely correct from a mathematical point of view, it accurately describes what is shown in the graph in a more understandable manner.

The volume rendering of the cluster densities reported in Figure 1C-1E, S1A and S1B were obtained by computing 3D kernel density estimates of the Ki67^+^clusters in each specimen, by using the R package ks^90^. Increasing cluster density is shown with three objects of different colours: transparent (white boundary), yellow and orange. The obtained 3D objects were visualised and exported as .obj files with the R package rgl (Murdoch and Adler, 2023).

### STATISTICAL ANALYSIS

#### Cellular composition of Ki67+clusters at 5wpl

The cellular composition of Ki67^+^clusters at 5 wpl (Figure 2) was analysed on entirely reconstructed clusters (n= 430) from 3-5 consecutive 3D reconstructed sections per mouse (n= 8 mice). Ki67^+^ clustered cells were manually counted using the MultiPoint tool in Fiji. As previously described^26^, clusters were defined as groups of at least 4 Ki67^+^ cells showing direct contact among their cell bodies. Of note, due to this stringent criterion used to define clustered cells, when pNBs were sparse in clusters of postmitotic NBs not showing direct contact among their cell bodies that cluster was excluded from the analysis. As a consequence, our analyses did not include the oldest cluster population in which the proliferative potential is fading off (see also Data Supplementary text 1, section 4.2).

#### Time course analyses of Ki67^+^cluster distribution, number, cellular composition and turnover

Time course analyses of Ki67^+^cluster (Figures 3, S4 and S5) were performed on three non-consecutive 400µm-spaced sections for each specimen. It is to be noted that the 63±9% of Ki67^+^clusters were partially included in the analysed sections (Figure S4B). However, the fraction of incomplete clusters was constant among specimens (*One-way ANOVA, F^(3,9)^=1.53, p=0.271*).

#### Clonal analyses of striatal astrocyte progeny

To visualize different reporters, we used anti-GFP antibody to recognize GFP, YFP, and CFP and anti-RFP antibody for RFP. GFP^+^ cells were discriminated according to the localization of the green staining: nuclear (corresponding to “nGFP”), cytoplasmic (corresponding to “cYFP”), or membrane-associated (corresponding to “CFP”).

For each Confetti specimen (Figures 4 and S6), all the consecutive sections containing striatal newborn cells were 3D reconstructed as described in the 3D reconstruction section (#C1: n=26 slices; #C2: n= 24 slices; #C3: n= 32 slices; #C4: n= 29 slices). The spatial localization and the type of each Ki67^+^cluster were manually annotated in TrakEM2, allowing to map the Ki67^+^clusters containing at least 1 Confetti Reporter^+^ cell in 3D space. Cluster coordinates were exported from TrakEM2 using the script *“EXPORT_Balls_3.3.py”* for further spatial statistical analyses.

The cellular composition and reporter expression of these Confetti clusters was manually evaluated on 3D reconstructions from high resolution confocal acquisitions (see the “Confocal imaging” section). The same stringent criterion used to define clustered cells was applied for considering dormant astrocytes as part of the cluster - and thus clonally related to the other cells it gave rise to. The same criteria were used for the quantification of clonally-related dormant astrocytes in subsequent fate-mapping analyses with Glast^CreERT2/+^xR26R-YFP mice.

Striatal astrocytes recombination efficiency was measured as the fraction of GFAP^+^SOX9+ cells expressing each Confetti reporter (Figures 4 and S6), in four fields of view for each specimen.

#### Genetic fate-mapping analysis of striatal astrocytes activation and cluster maturation

Fractions of YFP^+^ striatal clusters were counted over the entire thickness of 3 consecutive 3D reconstructed sections (Figure 5). The cellular composition of Ki67^+^clusters containing at least 1 YFP^+^ cell was evaluated only for Ki67^+^clusters entirely included in the volume (Figures 5 and 6). For the T-4 animals and two of the four T-7 animals that add less than 9 clusters within the first three sections, three or four additional slices were analysed.

#### Expression of Ascl1 and Sox9 in striatal Ki67^+^ cells

Ki67^+^ cells organized as single cells, pairs, trios or clusters were analysed on two consecutive reconstructed slices for each mouse (n=3; Figures 6 and S7). We distinguished between SOX9^High^ cells expressing this transcription factor at a similar level as the non-proliferating parenchymal astrocytes and SOX9^Low^ cells expressing it at barely detectable levels (Figure 6F-I).

#### Estimate of astrocyte activation rate

To estimate striatal astrocyte activation rate within the neurogenic area we segmented the SOX9^+^ nuclei using Ilastik^91^. Briefly, we trained a pixel classification model recognizing the nuclei centres, subsequently applied a watershed separation of individual objects using hysteresis thresholding and further trained an object classification model to recognise nuclei that were not separated, almost only couples of nuclei, or that were splitted, never more than in two pieces. The total number of nuclei was thus obtained by adding the number of individual nuclei, unsplitted nuclei*2 and splitted objects/2. The number of astrocytes and active Ki67^+^ structures were counted over the neurogenic area and its core, using the R packages spatstat^38^ and RImageJROI (https://github.com/davidcsterratt/RImageJROI). The neurogenic area and core were obtained by computing 3D kernel density estimates of the Ki67^+^clusters in each specimen with the R package ks^90^, and included respectively the 95% and 25% of clusters with increasing density (see also Figure S7I).

Due to the extremely high density of SOX9+ cells in the SVZ, quantifications in these region were performed manually, by counting the SOX9^+^Ki67^-^, the SOX9^+^Ki67^+^ and the SOX9^-^Ki67^+^ cells in the lateral wall of 3 SVZ from healthy mice. All counted objects, in both striatum and SVZ were counted stereologically by excluding those that touched one of the section surfaces.

#### Statistical tests

The results of all the analyses are reported in Table S1. We preferred to not overload the figures with the statistics to allow a better visualization of the data. We reported in the main text only the p-values of the comparisons we specifically cite there. All analyses were performed using R Statistical Software (R Core Team 2021).

#### Means comparisons

To select the proper statistical test we always checked if the data were normally distributed, using the Shapiro–Wilk test, and homoscedastic, using Levene’s test. Data that fulfilled those requirements were compared with the t-test in the case of two groups or with the ANOVA in case of three or more groups. ANOVA analyses that returned significant F values were followed by Tukey’s post hoc test. In the case of non-normally distributed and/or non-homoscedastic data, we used the nonparametric alternatives: the Mann–Whitney U test and the Kruskal-Wallis test to compare two, or more than two groups, respectively. Significant Kruskal-Wallis tests were followed by Dunn’s test for multiple comparisons (p-values were adjusted according to the Benjamini-Hochberg method). In case of paired samples, the Wilcoxon signed-rank test was used instead of the Mann-Whitney U test.

#### Comparison of proportions

Fisher’s Exact Test was used to compare proportions among count data with binary outcomes (e.g. fraction of Ki67^+^clusters associated or not associated with postmitotic NBs). We use this test instead of the more conventional Chi-square test as it is more suitable to compare groups with low numbers of observations. For comparisons among more than two groups, significant Fisher’s Exact Tests were followed by Pairwise Fisher’s Exact Tests in which p-values were adjusted according to the Holm-Sidak method.

#### Comparison of frequency distributions

The Kolmogorov-Smirnov test was used to compare two distributions, while the Anderson-Darling k-sample test to compare three or more distributions.

#### Spatial statistics (point pattern analysis)

##### Evaluation of Ki67^+^cluster distribution

To assess if Ki67^+^clusters distribute according to specific spatial patterns we used the G(*r*) function^38^. This summary function represents the cumulative frequency distribution of the first-order nearest neighbour distances (*r*) and it is one of the simplest techniques to find clustering or dispersion in a point pattern. Although a 3D version of this function is implemented in spatstat, we preferred to use its 2D counterpart as it allows, unlike the 3D version, the use of irregular polygonal areas describing the region from which the point pattern was retrieved. As shown in Figures 1 and S1, the striata and neurogenic areas are highly irregular and cannot be approximated to a parallelepiped. Further, since the analyses were performed on a few consecutive sections, cluster dispersion was much greater on the X and Y axis than in Z, suggesting the reliability of a 2D projection analysis.

The G function calculated from the data (*G_(obs)_(r)*) is visually inspected and compared to the theoretical one obtained from a homogeneous Poisson process (*G_(theo,H0)_*) that represents the null hypothesis (*H0*) of complete spatial randomness (CSR). If *G_(obs)_(r)* lies at the left or at the right of *G_(theo,H0)_(r)* it suggests clustering or dispersion, respectively. Yet, to avoid false-positive results, each *G_(obs)_(r)* was compared with 999 Monte Carlo simulations of *G_(theo,H0)_(r)*, and the graphs in Figure 2G and S3A-S3B report the mean of these simulations *G_(mean,H0)_(r)* and the simulation envelopes. For statistical comparisons (Figures S3A-S3B), we constructed simultaneous envelopes having a constant width around *G_(mean,H0)_(r).* The width is defined by finding the most extreme deviation, at any value of *r*, of the 999 simulated *G_(theo,H0)_(r)* from *G_(mean,H0)_(r)*. Thus, the visual interpretation of the graphs coincide with the results of Maximum Absolute Deviation Tests^38^. We rejected the null hypothesis of CSR if *G_(obs)_(r)* felt outside of the envelope at any value of *r*. As a complementary strategy to compare the experimental distributions with the simulations we used the more powerful Diggle-Cressie-Loosmore-Ford test^38^, that, instead of the maximum absolute deviation, uses the integrated squared deviation between *G_(obs)_(r)* and *G_(theo,H0)_(r)* over the the range of measured *r*. For representative purposes, in Figure 2G we constructed pointwise envelopes instead of simultaneous ones, as we thought they were simpler to visually interpret (e.g *G_(theo,H0)_(r) > 0* at every value of *r*). However, we did not use such envelopes for statistical purposes as they would be more suitable for testing if clustering/dispersion exist at specific values of *r*, which was not what we intended to test here^38^.

To perform these analyses, the R packages spatstat^38^ and RImageJROI (https://github.com/davidcsterratt/RImageJROI) were used. The pooled p values shown in the main text were obtained with the R package poolr^92^ by using the Fisher’s method on the p values reported in Figure S3A-S3B.

##### Global spatial autocorrelation

To analyse the 3D distribution of Ki67^+^cluster features (cellular composition in Figure 2 and Confetti reporter expression in Figure 4I) we used the Moran’s I index of global spatial autocorrelation^93^. It’s a widely used technique in spatial econometrics^94^ that, similarly to a correlation coefficient, provides an estimate of how closer objects are more similar than distant ones. Thus, it’s a powerful technique to search for spatial clustering of similar objects. It bears the great advantages of being a “distance-based” metric. So, unlike other summary functions such as the G function, there’s no need to specify the study area/volume from which the point pattern was retrieved. Further, no edge correction methods are needed, avoiding the possible introduction of biases by the selection of specific methods or areas. This greatly helps in assessing the distribution of objects embedded in highly irregular and variable environments like the lesioned striata.

A positive Moran’s I index suggests clustering of similar objects, while a negative one is indicative of homogeneous dispersion. Yet, the Index value depends on the number of data points and the bigger the sample the more it approaches 0, which represents the absence of spatial autocorrelation. Thus, the sole examination of the Moran’s I Index is difficult to interpret. To understand if the observed data distribution was significantly different from complete spatial randomness, we performed a permutation test based on Monte Carlo simulations. The simulations randomly permuted the cluster features over the same point pattern distribution and every time the resulting Moran’s I Index was measured. Finally, a p-value was calculated based on the proportion of simulations giving a statistics that is as or more extreme than the observed data. To perform this analysis, the R package ade4^95^ was used.

### Key resources table

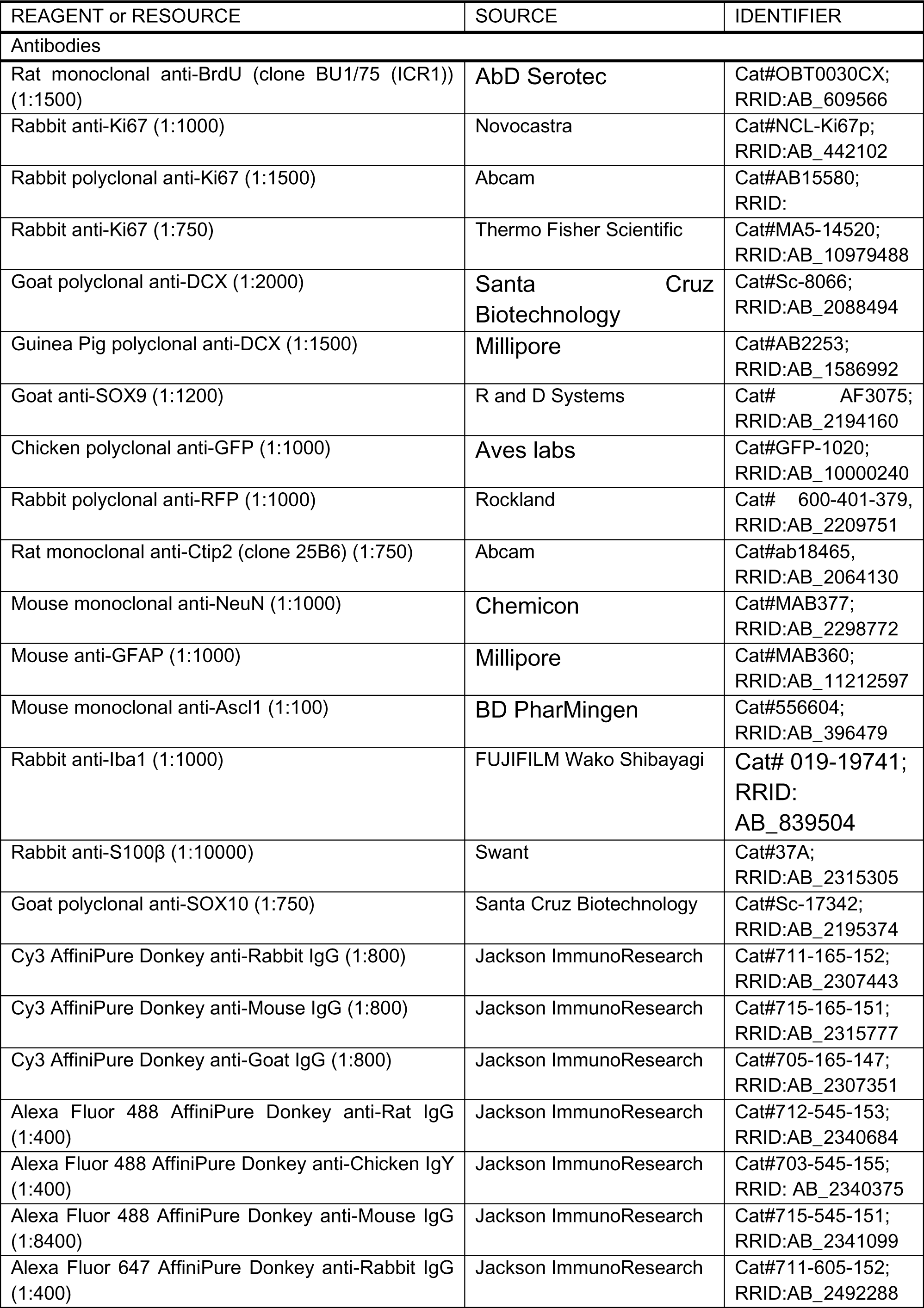
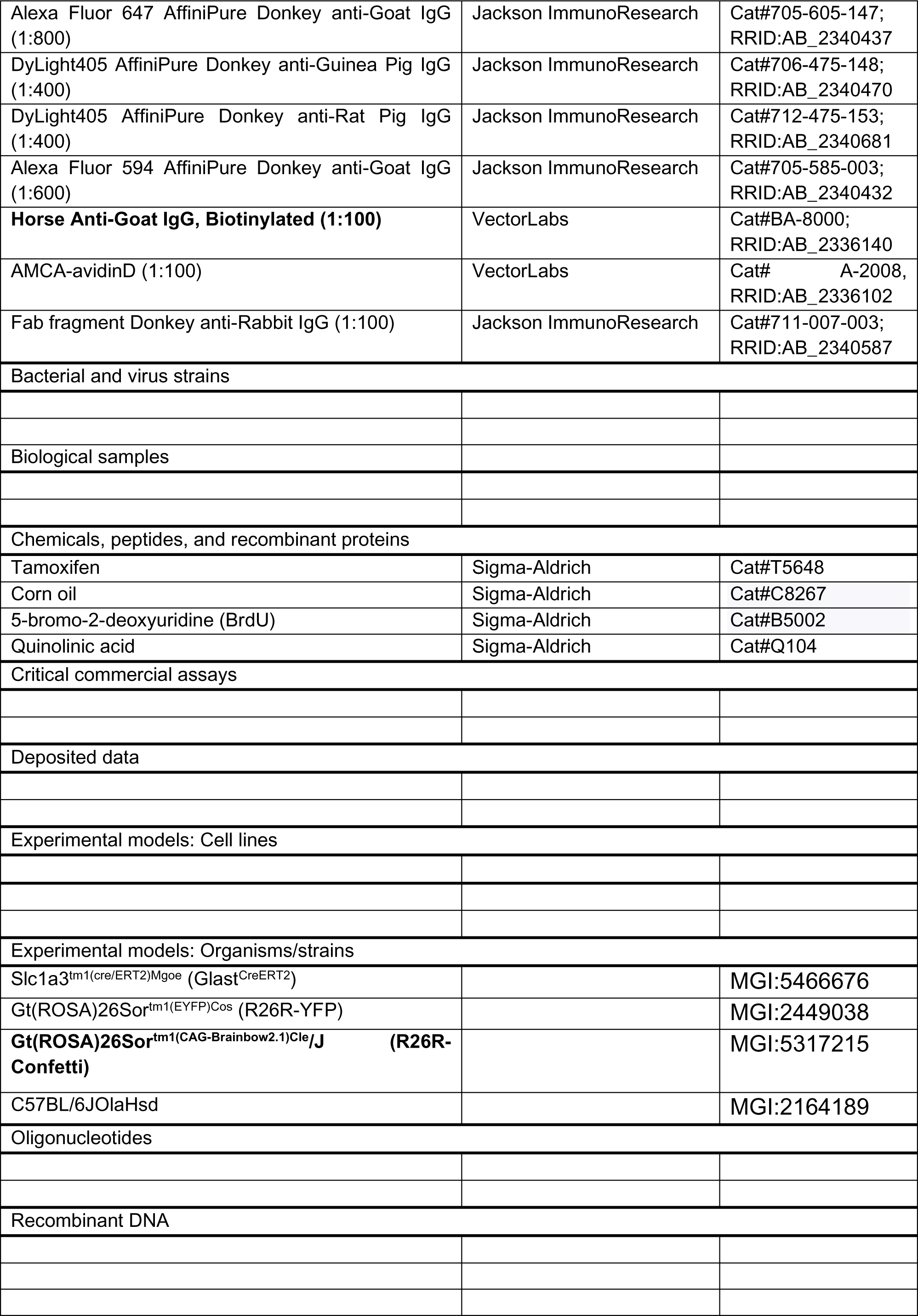
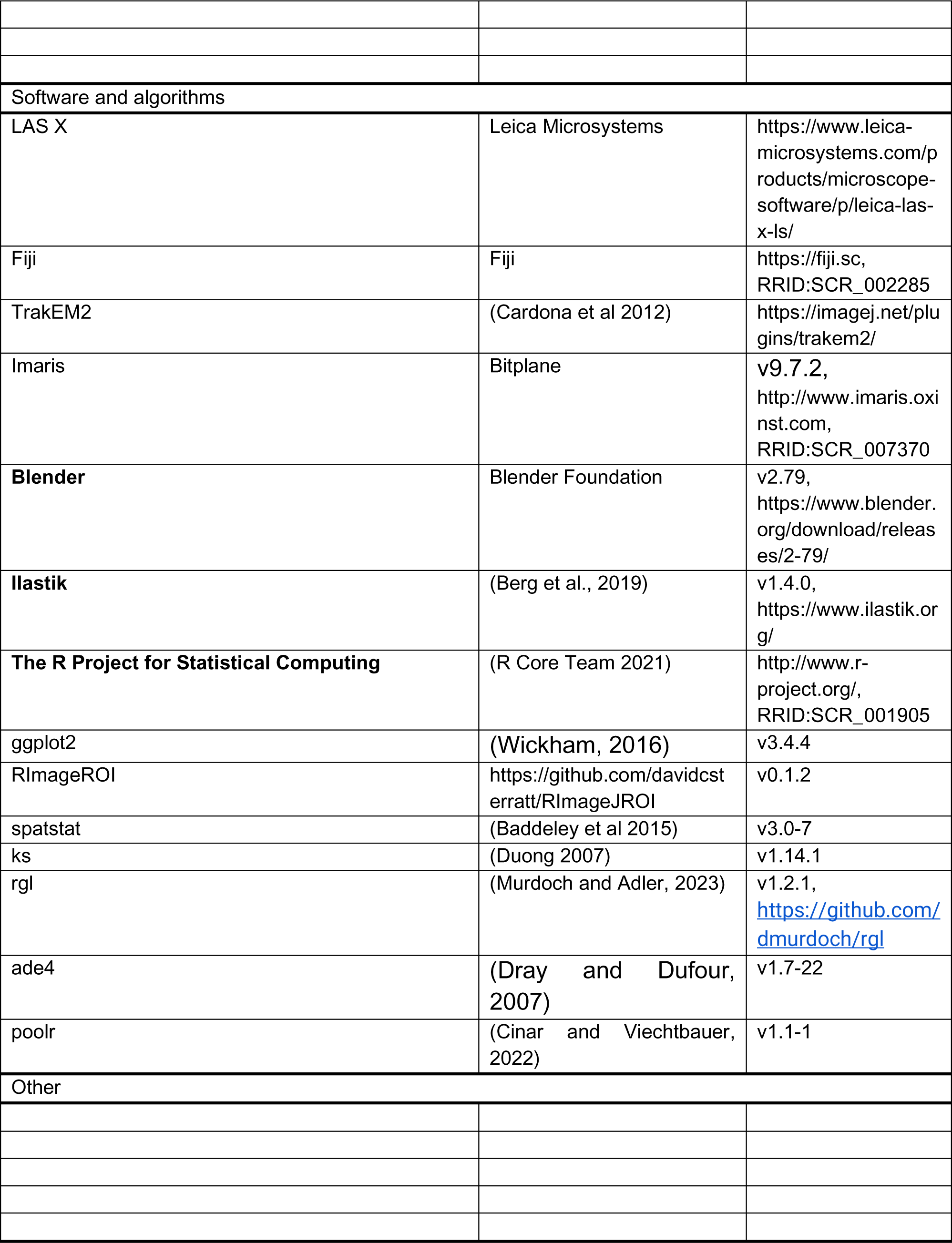

