## Supplementary Figures for "Dynamic spatiotemporal activation of a pervasive neurogenic competence in striatal astrocytes supports continuous neurogenesis following injury"

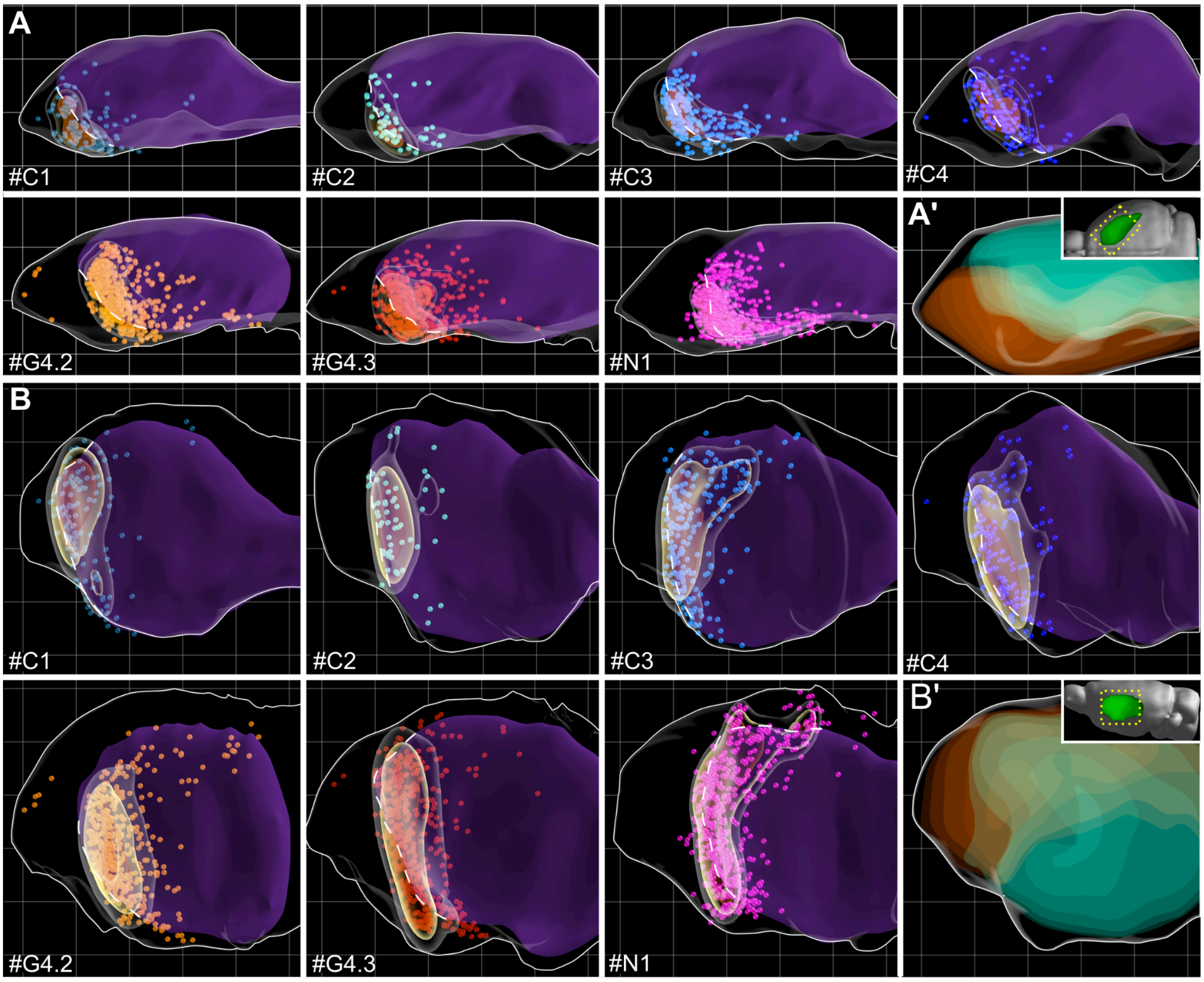


***Figure S1. Neurogenic foci distribution in individual specimens, related to Figure 1***

(A) Dorsal and (B) Medial views of the 3D reconstructions of the Ki67$^+$ clusters over the entire striatum of 7 specimens registered to a common reference space ( see also Figure 1D and Video S1). The lesion is in purple. A white dashed line shows the lesion border in the neurogenic area. Dots: Ki67$^+$ clusters (25 um diameter) colour-coded as in Figures 1D-1E. Increasing cluster density is rendered as transparent, yellow and orange volumes. (A') Dorsal and (B') medial view of a reference striatum with the associative functional domain of the striatum, receiving projections from the anterior cingulate cortex, in orange and the somato-motor domain, receiving afferents from the somatosensory cortex, in cyan. It highlights the medial and lateral striatum, respectively (3D reconstructed from Hintiran et al. 2016, see Method details). Grid lines are 500 μm spaced.

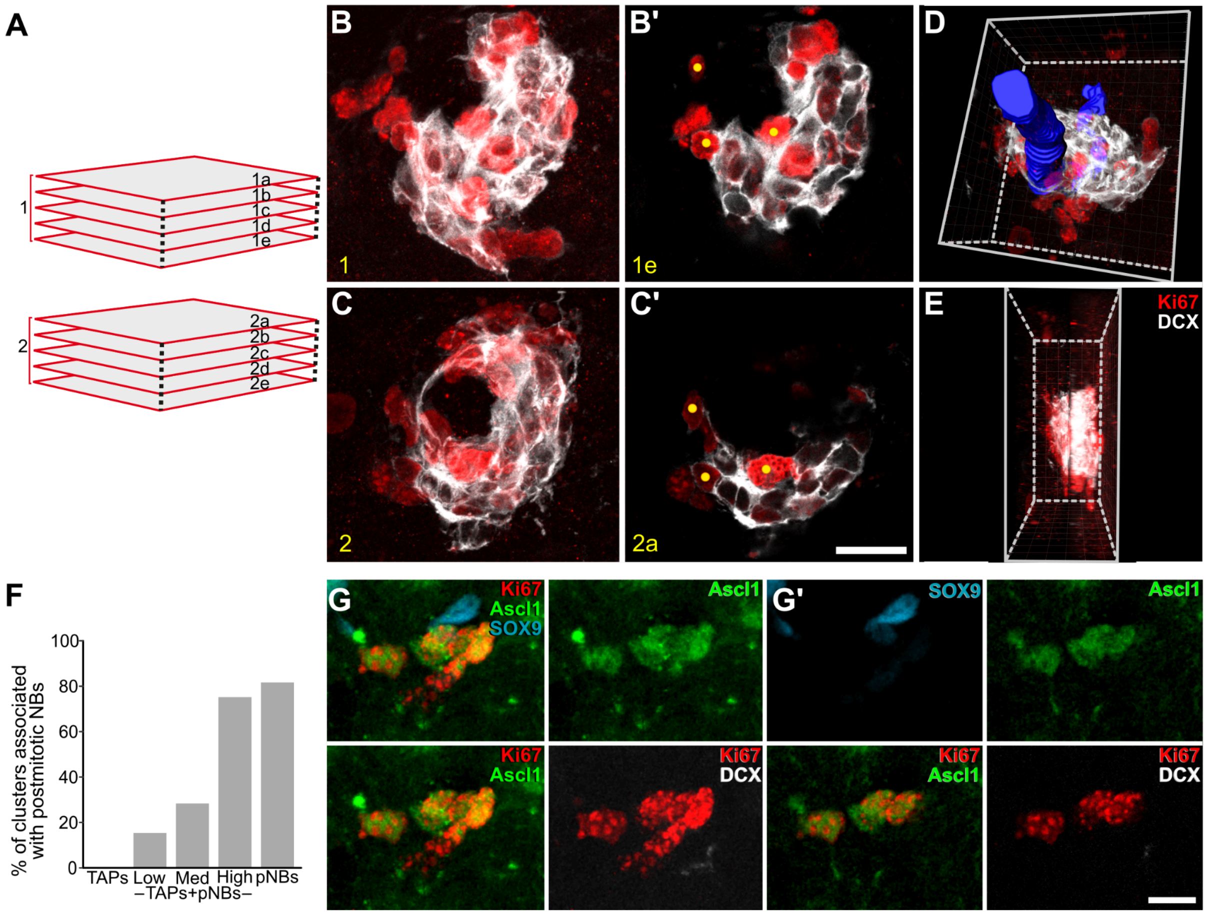


***Figure S2. Method of high resolution 3D reconstruction and cellular composition of Ki67⁺ clusters, related to Figure 2***

(A) Schematic view of the focal planes of two confocal stacks acquired from two consecutive sections. (B,C) Maximal intensity projection of sections 1 and 2 showing a Ki67$^+$ cluster splitted between the two sections. (B', C') The last intact focal plane (the section surface is usually not perfectly flat) of section 1 and the first one of section 2, respectively. Yellow dots highlight corresponding Ki67$^+$ cells in the two sections. (D) Imaris 3D rendering of the reconstructed cluster. A blood vessel was manually segmented and shown in blue. (E) Side view of the reconstructed cluster. (F) Percentage of Ki67$^+$ clusters associated with postmitotic NB by cluster types. (G,G') TAPs-only cluster stained for Ascl1 and Sox9. (G) represents a maximal intensity (MAX) projection, (G') a single focal plane.

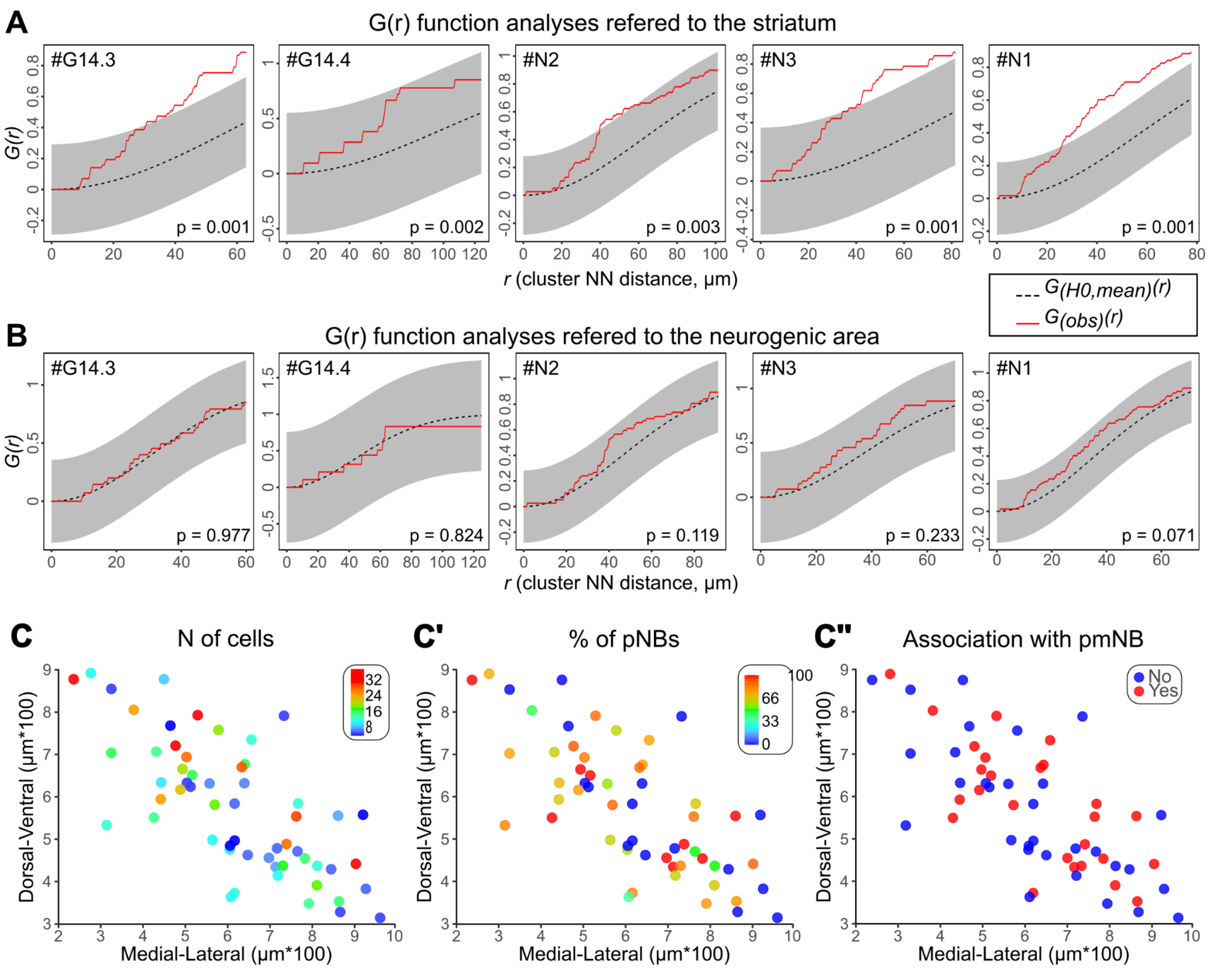


***Figure S3. Spatial analysis of Ki67$^{+}$ cluster distribution, related to Figure 2***

(A,B) G function analyses of Ki67$^{+}$ clusters distribution in respect of the striatum (A) or the neurogenic area (B) in five specimens. Red lines: $G_{(obs)}(r)$, functions calculated from the experimental data. Black dashed lines: $G_{(H0,mean)}(r)$, mean functions of the simulations of complete spatial randomness (*H0* = homogeneous Poisson process). Grey areas: 95% confidence envelopes. Unlike in Figures 2G-2H, here simultaneous - instead of pointwise - envelopes were constructed so that we could reject the null hypothesis if the $G_{(obs)}(r)$ function lied outside the envelope at any value of *r.* As an additional approach for statistical comparison the Diggle-Cressie-Loosmore-Ford (DCLF) test was performed and the resulting p-values are reported for each specimen and area (see also Method details). (C-C'') X-Y projection of clusters shown in Figure 2F coloured by size, percentage (%) of pNBs and association with postmitotic NBs.

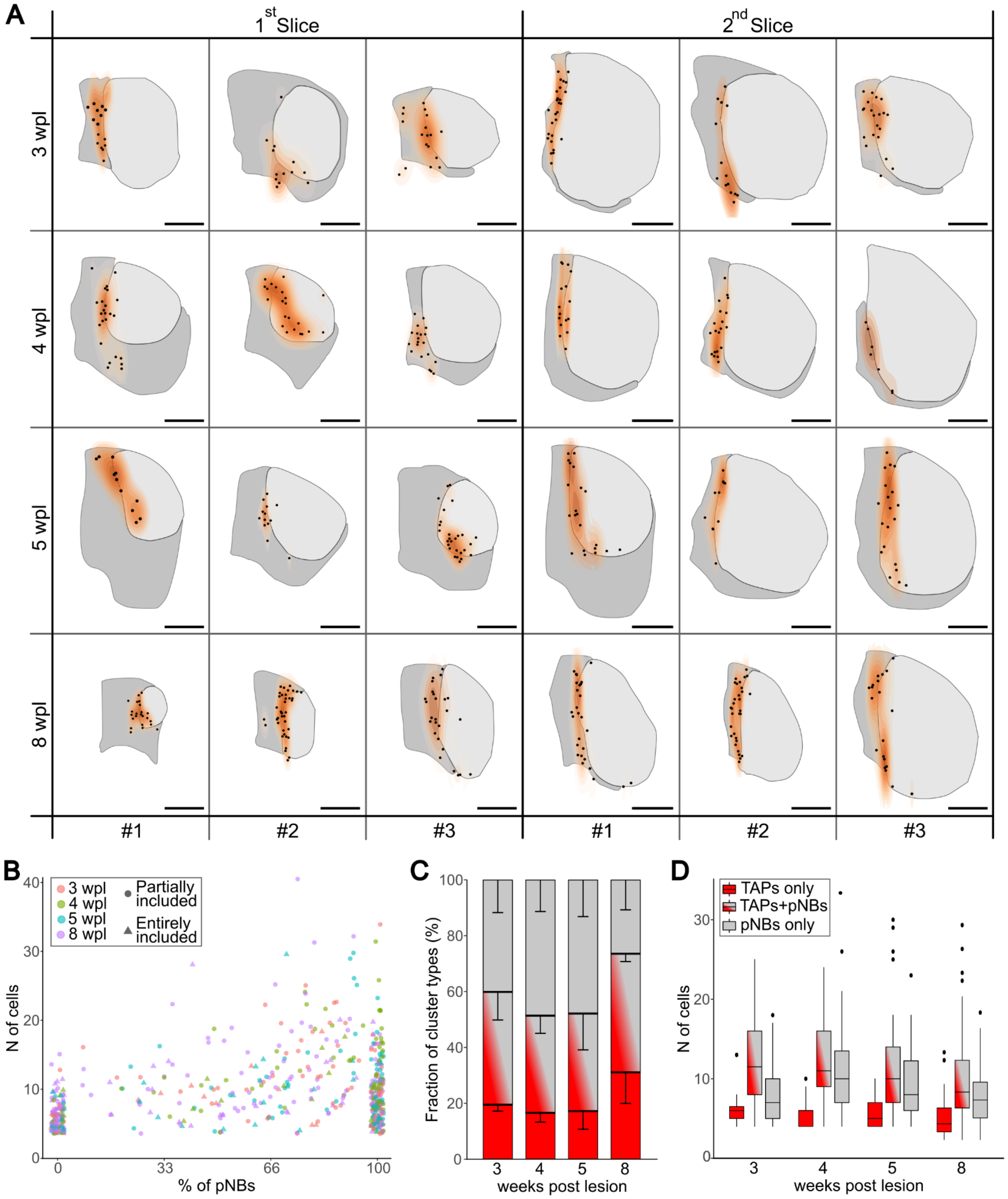


***Figure S4. Ki67⁺ cluster distribution and cellular composition do not change overtime, related to Figure 3***

(A) Schematic view of the Ki67⁺ cluster distribution in the two more anterior sections of the three analysed. Only three specimens (#) per group were included in this representation. Each cluster is shown as a dot and the kernel density estimate of Ki67⁺ cluster distribution is shown on a graded orange scale. Higher cluster density corresponds to higher orange intensity. Of note the highest cluster density is always observed at the border between the lesioned (light grey) and spared (darker grey) striatal tissue. (B) Percentage of pNBs vs number of cells in each Ki67⁺ cluster. The colour of the dots indicate the group from which those Ki67$^+$ clusters belong, while the shape indicates if the cluster was partially (circle) or entirely (triangle) included in the analysed sections. (C) Relative fraction of cluster types at each time point (Table S1, TAPs-only: *p=0.049*, TAPs+pNBs: *p=0.873*, pNBs-only: *p=0.213*). Only concerning TAPs-only clusters we found a significant difference overtime. Yet, post hoc analyses revealed a barely significant difference only when comparing 5wpl with 8wpl (Table s1, *p= 0.047)*. The low number of specimens and the high variability may explain this small difference. (D) Number of cells per cluster at different time points. Data were splitted into the three main cluster types and shown as box and whisker plots to highlight the data distribution (Table S1, TAPs-only: *p=0.468*, TAPs+pNBs: *p=0.530*, pNBs-only: *p=0.073*). Outliers are shown as black dots. Scale: (A) 500 μm.

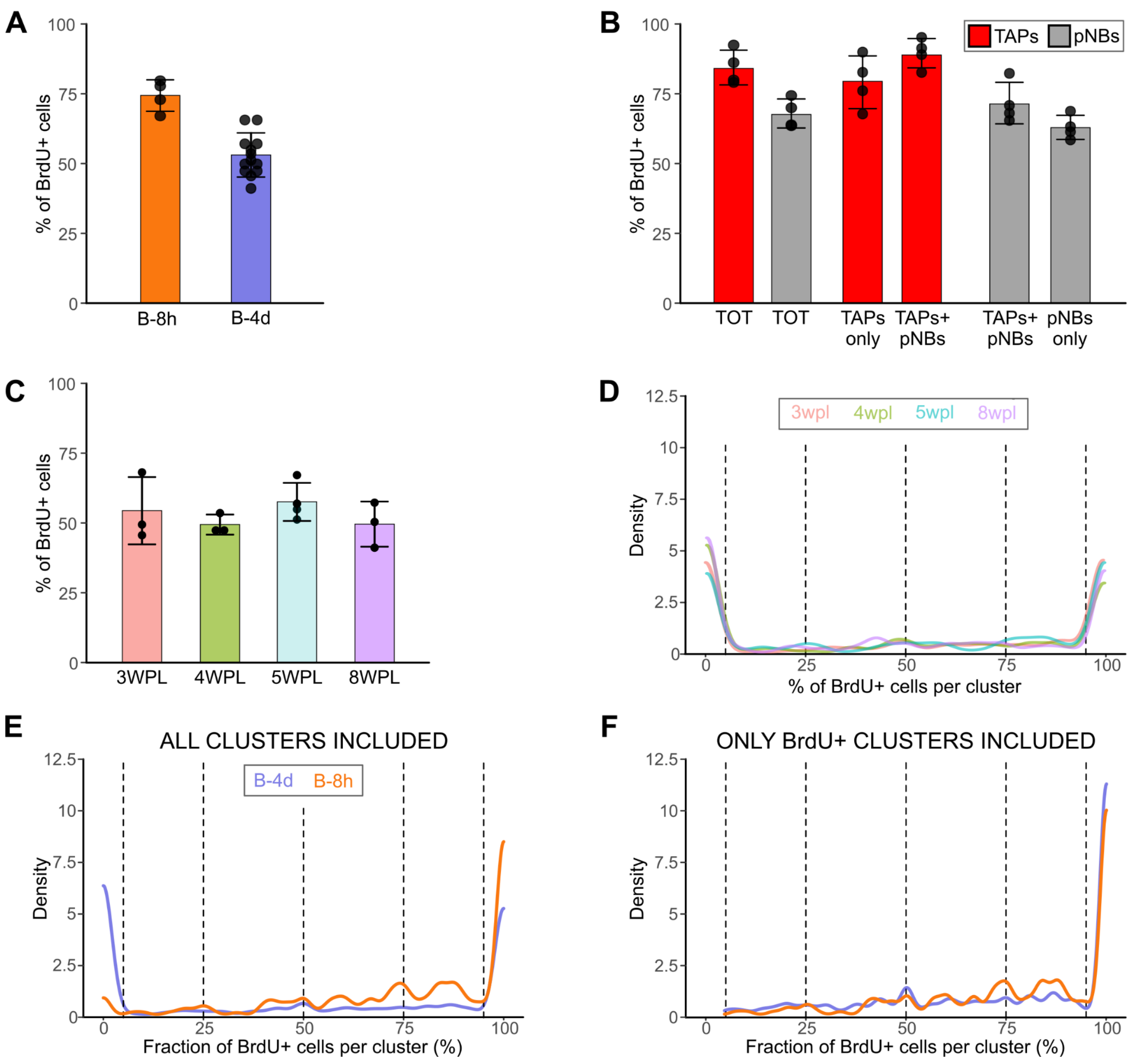


***Figure S5. No cellular turnover occurs within individual Ki67⁺ clusters, related to Figure 3***

(A) Percentage of BrdU⁺ cells among all the Ki67⁺ clustered cells in B-8h and B-4d groups. (B) Percentage of BrdU+ TAPs (red) or pNBs (grey) in the whole population (TOT) or by cluster type (TAPs-only, TAPs+pNBs, pNBs-only) in the B-8h group. (C) The percentage of BrdU⁺ cells among all the Ki67⁺ clustered cells is constant among B-4d groups. (D) The relative distribution of the % of BrdU+ cells per cluster is similar among the B-4d group. (E,F) Same as D, but all the clusters of B-4d (violet) and B-8h (orange) were pooled together, respectively. In (E) all clusters while in (F) only the BrdU⁺ ones are considered. If BrdU dilution under detection level or cellular turnover occurred at substantial levels in Ki67⁺ clusters, we would expect a progressive leftward shifting of the orange curve. The distributions of B-4d and B-8h including all clusters are statistically different (Table S1, *p<0.001*), but this is due to a net peak at 0% of BrdU⁺ cells (i.e. BrdU⁻ clusters) for B-4d clusters. Indeed, when these clusters are removed in (F) the cluster populations at the two time-points showed highly similar distributions (Table S1, *p=0.247*). These analyses strongly support that neither BrdU dilution under detection levels, nor cellular turnover occurred within Ki67+ clusters. Thus, BrdU⁻ clusters observed at each time point are newly formed structures. Data in (A), (B) and (C) are reported as: barplot= mean±SD, dots= individual specimens.

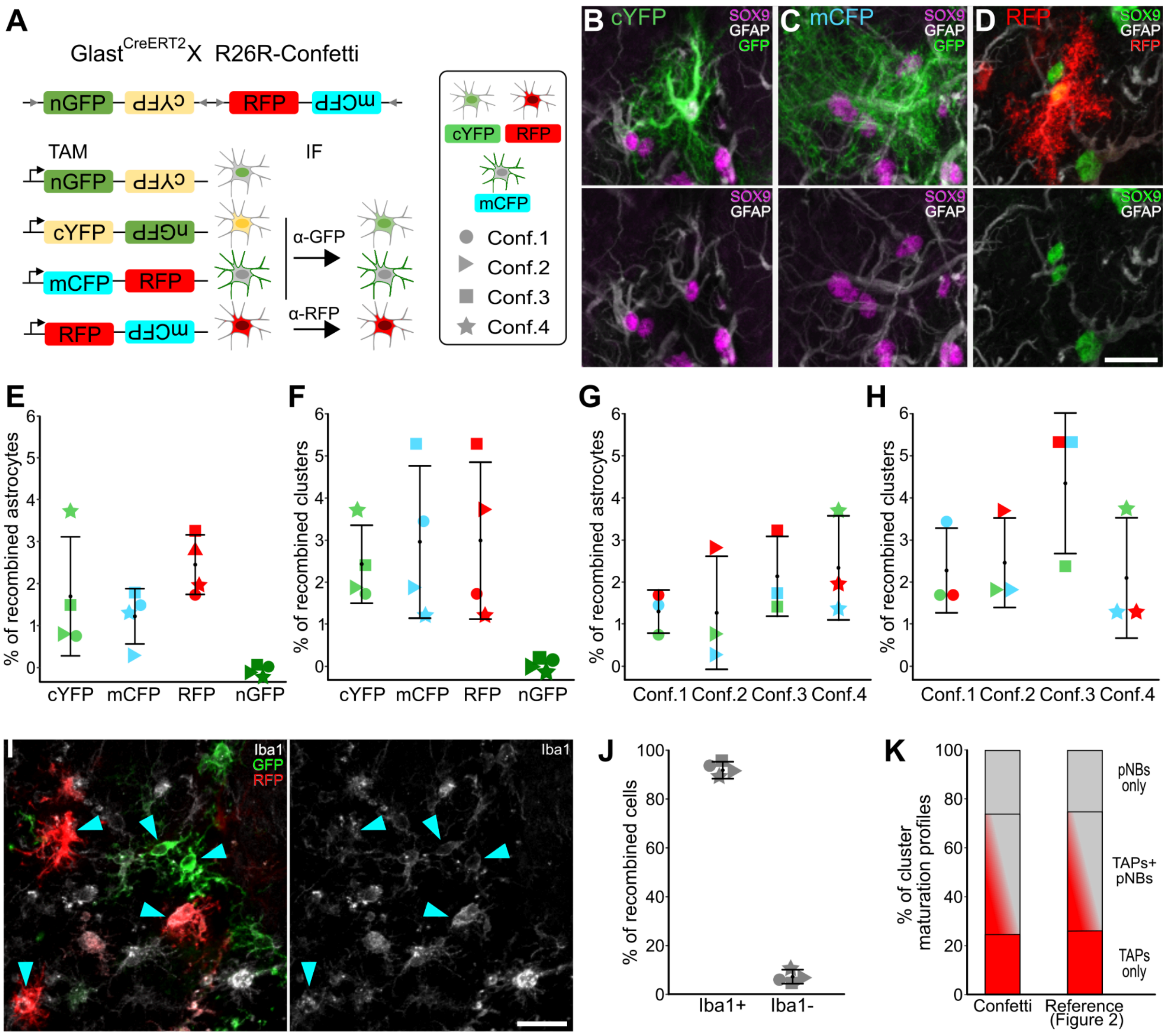


***Figure S6. Confetti reporters are expressed homogeneously among striatal astrocytes, Ki67⁺ clusters and microglial cells of different specimens, related to Figure 4***

A) Schematics of the R26R-Confetti locus. Tamoxifen (TAM) administration induces the stochastic and non-combinatorial expression of 1 of the 4 reporters. After immunostaining with α-GFP and α-RFP antibodies recombined cells showed either a cytoplasmic GFP staining (cYFP), a membrane-localised GFP staining (mCFP) or an RFP staining (RFP). The nGFP was never observed. Each specimen is represented with a different symbol throughout the figure. (B-D) Representative images of recombined $SOX9^+Gfap^+$ striatal astrocytes. (E) Fraction of $SOX9^+Gfap^+$ cells or (F) Ki67+ clusters expressing the cYFP, mCFP or RFP. (G,H) Same as (E,F) but different specimens are on the x-axis to show that they have similar recombination efficiency of striatal astrocytes and $Ki67^+$ clusters (see also Table S1). The representations from (E) to (H) shows that the fraction of striatal astrocytes and Ki67+ clusters expressing the different Confetti reporters were similar between reporter and specimens. (I) Representative image of recombined Iba1+ microglial cells. (J) Fraction of recombined cells ($GFP^+$ and $RFP^+$ pooled together) expressing the microglial marker Iba1. (K) Relative frequencies of $Ki67^+$ cluster maturation profiles in the Confetti clusters and the reference population of clusters analysed in Figure 2. Scale (B-D) and (I) 20 µm.

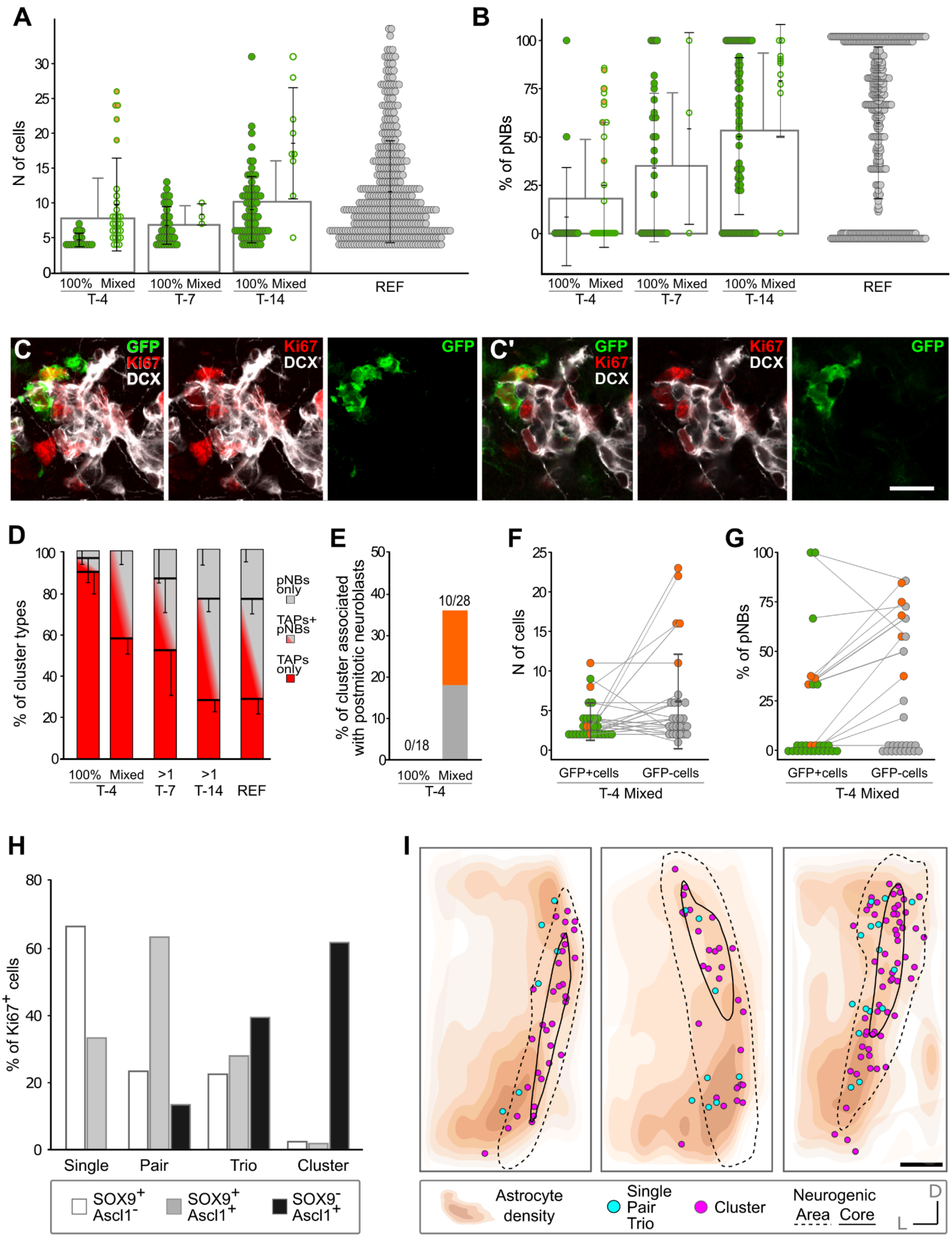


***Figure S7. Striatal astrocytes activate only at the beginning of neurogenic foci life, related to Figures 5 and 6***

(A,B) Number of Ki67$^{+}$ cells and % of pNBs per cluster. The data of the lineage tracing groups (T-4, T-7 and T-14) were compared with the data from the reference population of striatal clusters analysed in Figure 2 ("REF", see also Tables S1). At each time-point, data from different specimens were pooled together and shown as mean (white box with grey contour) ±SD. The data are also reported splitted between clusters entirely composed of $YFP^+$ cells ("100%") and those composed of a mixture of $YFP^+$ and $YFP^-$ cells ("Mixed"). Like in Confetti mice, all the Mixed clusters in T-7 and T-14 likely represent cluster fusion as they displayed a more advanced maturation profile than their 100% counterparts and their number correlated with the mean nearest neighbour distance among clusters, a measure of cluster density (Regression analysis with exponential decay fit: *p=0.004*). Concerning T-4 Mixed clusters, only 5 could be derived from cluster fusion for the above mentioned criteria (the orange ones, as in Figures S6E-S6G; see also Figure 6I). All others displayed only slightly more advanced maturation profiles than the 100% $YFP^+$ ones. Further, their number did not correlate with the mean nearest neighbour distance among clusters (Regression analysis with exponential decay fit: *p=0.957*) suggesting that fusion due to high $Ki67^+$ cluster density cannot explain all the T-4 mixed clusters. (C,C') Representative image of one of the five T-4 clusters that likely resulted from cluster fusion. (C) is a MAX projection including the entire cluster while (C') a single focal plane in which it is easier to see which cell is $YFP^+$ and which one is not. (D) Relative proportion of different cluster types among $YFP^+$ clusters. T-4 clusters were splitted into 100% $YFP^+$ and Mixed clusters, while in other groups all clusters with at least 1 $YFP^+$ cell (>1) were considered together. Data are shown as mean±SD among specimens. The 100% T-4 clusters are the most immature ones, while the T-4 Mixed resemble the T-7 but are much more immature than T-14 and REF clusters. (E) Fraction of $Ki67^+$ clusters associated with postmitotic NBs among 100% $YFP^+$ and Mixed clusters of T-4. (F) Number of cells and % of pNBs (G) among the $YFP^-$ and the $YFP^+$ cells of T-4 Mixed clusters. Each dot represents a cluster. Of note, when the five putatively fused clusters were excluded from the analyses, we did not observe any difference in the number of cells and in the % of pNBs among the $YFP^+$ and $YFP^-$ compartments of T-4 Mixed clusters (Table S1). These data indicate that both groups of cells originated almost simultaneously at the beginning of the $Ki67^+$ cluster life. This is in line with the hypothesis that striatal astrocytes activate only at the beginning of $Ki67^+$ cluster life and with the possibility that $Glast^{CreERT2}$ recombination occurs up to the 2- or 3-cell stage. (H) Percentage of $Ki67^+$ cells organised as single, pair, trio or clusters that express only Sox9 (white), Sox9 and Ascl1 (grey) or only Ascl1 (black). (I) Astrocyte density map (increasing orange intensity) and spatial distribution of $Ki67^+$ clusters (magenta) and single/pair/trios (light blue) in two consecutive 3D-reconstructed slices (n= 3 mice). The neurogenic area and its core used to quantify cell densities in Figures 6L and 6N-SP are depicted as black lines (see also Method details). Quantifications in Figures 6K, 6L and 6N-6P were taken from these distributions, also considering the Z dimension. Scale: (C-C') 15 μm; (I) 250 μm.
