## Supplementary Text 1 - Mathematical modeling for "Dynamic spatiotemporal activation of a pervasive neurogenic competence in striatal astrocytes supports continuous neurogenesis following injury"

In this supplementary text we will explain the modelling approach chosen to support our findings and to verify the underlying hypotheses on cluster turnover dynamics and associated cell fate choices. First, in section 1 we will argue the dynamics of initiation of proliferative clusters and general cluster turnover dynamics, while in sections 2 - 4 we will compare the predictions of different models for cluster turnover dynamics with the cluster data, to assess which models are consistent with the experimental data.

We will use a slightly different notation of cell types when modelling the cell fate dynamics. In particular, we identify cell types only by the markers Ki67 and DCX, and whether they appear within a cluster. Thus, we pool astrocytes and TAPs, both being DCX-negative cell types that are cycling, that is, expressing Ki67, as one cell type category, calling them **cK cells** (for **c**lustered **K**i67). pNB on the other hand are called **cKD** cells (**c**lustered **K**i67 and **D**CX), while postmitotic neuroblasts, which do not express Ki67 are called cD cells (**c**lustered **D**CX).

### 1. Cluster initiation timings

First we assess the timing of the initiation of cluster proliferation. To address this question we make use of the fact that GLAST labelling of cells only occurs in astrocytes, or only for a limited amount of time upon differentiation. In Figure 4 of the main text, we established that clusters originate from the activation and initial division of an astrocyte each, but cycling astrocytes are absent in aged clusters (see also Figure 6, main text). We can therefore assume that clusters can only be induced to express YFP – via Cre-recombination upon administration of tamoxifen – before the founding

*Preprint submitted to Stem Cell Reports* *March 22, 2024*

astrocyte has divided for the first time (*initiation*), and up to some defined time after that time point. This is confirmed by the finding that for most clusters which contain YFP+ cells, all cells are labelled, meaning that the cell of origin has been labelled initially. Hence, we assume that induction of cells in a proliferating cluster is not possible when a certain time has passed since initiation of the cluster. For convenience, we will in the following refer to the time point when a (previous) astrocyte cannot recombine anymore as "initiation", keeping in mind that the true initiation may have happened some time before that.

Let us now denote by $Y(t_0)$ the number of YFP+ clusters measured at the time of culling and harvesting of clusters, $t = 0$, which have been induced at time $t_0 < 0$ [1]. These are those clusters which have been initiated after the time point $t_0$, and which 'survive' until the time of culling, $t = 0$, that is, they remain proliferating. Hence, we can relate this to the number of clusters initiated between any two time points $t_1$ and $t_2$, $\Delta N$ by,

$$Y(t_2) - Y(t_1) = \Delta N \times p_{\text{surv}}(t_1, t_2) \tag{1}$$

where $p_{\text{surv}}(t_1, t_2)$ is the proportion of clusters initiated between $t_1$ and $t_2$ which survive until being harvested at $t = 0$.

Since we cannot directly compare $Y(t_1)$ and $Y(t_2)$ in the same animal, and since the total number of clusters may vary significantly between clusters it is helpful to normalise this quantity to allow a more accurate comparison. For that purpose, we divide by $\bar{N}_{\text{tot}}$, the mean total number of clusters over the different animals, and take the expected value (indicated by angular brackets),

$$\left\langle \frac{Y(t_2)}{\bar{N}_{\text{tot}}} \right\rangle - \left\langle \frac{Y(t_1)}{\bar{N}_{\text{tot}}} \right\rangle = \left\langle \frac{\Delta N p_{\text{surv}}(t_1, t_2)}{\bar{N}_{\text{tot}}} \right\rangle \tag{2}$$

The proportion of surviving clusters, $p_{\text{surv}}$ is a property of individual clusters and does not depend on the total number of clusters $N_{\text{tot}}$ nor the number of initiated clusters $\Delta N$, hence $\left\langle \frac{\Delta N p_{\text{surv}}(t_1,t_2)}{N_{\text{tot}}} \right\rangle = \left\langle \frac{\Delta N(t_1,t_2)}{\bar{N}_{\text{tot}}} \right\rangle \bar{p}_{\text{surv}}(t_1, t_2)$, where $\bar{p}_{\text{surv}} = \langle p_{\text{surv}} \rangle$ is the expected proportion of surviving clusters, that is, the probability of survival. Now we define $y(t) = \left\langle \frac{Y(t)}{\bar{N}_{\text{tot}}} \right\rangle$, and $\Delta n(t, t') =$

[1]Note that induction occurs by definition at negative times

$\left\langle \frac{\Delta N(t,t')}{\bar{N}_{\text{tot}}} \right\rangle$ is the proportion of clusters, taken over all animals, initiated between timepoints $t_1$ and $t_2$. With this we have,

$$y(t_2) - y(t_1) = \Delta n(t_1, t_2)\bar{p}_{\text{surv}} \tag{3}$$

Finally we divide by $(t_2 - t_1)$ and take the limit $t_2 \to t_1$ $(t_2 > t_1)$, to get,

$$r(t) = \frac{\frac{d}{dt}y(t)}{\bar{p}_{\text{surv}}(t)} \tag{4}$$

where we identified the time derivative $\frac{dy}{dt} = \lim_{\Delta t \to 0} \frac{y(t+\Delta t) - y(t)}{\Delta t}$ and the normalised cluster initiation rate $r(t) = \lim_{\Delta t \to 0} \frac{\Delta n(t,t+\Delta t)}{\Delta t} = \frac{dn_{\text{tot}}}{dt}$, the proportion of clusters initiated per time unit, with $n$ being the total normalised number of initiated clusters.

Hence, given the cluster survival probability, we can identify the cluster initiation rate by the slope of the function $y(t)$. The latter can be sampled by counting YFP+ clusters and total clusters at certain time points $t$ of clonal induction.

However, the cluster survival probability is a priori unknown, but we know that immediately after cluster initiation it is $p_{\text{surv}} = 1$ and it can only decrease with time. Thus, the slope of the function $y(t)$ does for small times $t$ correspond to the cluster initiation rate $r$, while it may saturate to a plateau when $p_{\text{surv}}$ drops to zero. Yet, $y(t)$ remains a lower bound for $r$.

The measured ratio $y_{\text{exp}} = Y/N$, where $Y$ is the number of YFP+ clusters (containing at least one YFP+ cell) among $N$ total clusters is shown in Figure 5D of the main text. This serves as an estimate for $y(t)$ and we notice that indeed this slope is positive for $t$ close to $t = 0$, meaning that the cluster initiation rate is non-zero. **This shows that clusters are initiated not at a given time point, but in a staggered manner, throughout time (at least close to the culling time point)**.

Further away from $t = 0$ the slope of $y(t)$ vanishes. We now need to discern whether this is due to cluster initiation rate being vanishing or due to $p_{\text{surv}}$ vanishing, i.e. clusters stopping to proliferate. In the following we will show that the cluster initiation rate is non-zero also far from $t = 0$ and most likely nearly constant over time.

Let us explore the possibility that $p_{\text{surv}}$ remains non-zero while $r(t) \to 0$. However, measurements from culling at earlier time points show that clusters are already present at 3 weeks post-lesion in similar numbers as at 5 weeks post-lesion (culling time point for clonal induction assays). This means not only that the cluster initiation rate must be non-zero at earlier time points (well before 3 weeks post-lesion), but also that cluster numbers stay the same over time, i.e. attain a steady state.

We now express the surviving cluster proportion $n_{\mathrm{tot}}(t)$ as,

$$n_{\mathrm{tot}}(t) = \int_{t'<t} r(t') p_{\mathrm{surv}}(t-t')\, dt' \tag{5}$$

The condition that this number does not change over time is expressed by the vanishing of the time-derivative,

$$0 = \frac{d}{dt} n_{\mathrm{tot}} = \frac{d}{dt} \int_{t'<t} r(t') p_{\mathrm{surv}}(t-t')\, dt' = r(t) p_{\mathrm{surv}}(0) + \int_{t'<t} r(t') \frac{d}{dt} p_{\mathrm{surv}}(t-t')\, dt' \tag{6}$$

Here we used Leibniz' integral derivation rule, $\frac{d}{dt}\left(\int_{s<t} f(s,t) ds\right) = f(t,t) + \int_{s<t} \frac{d}{dt} f(s,t)$.

Now we identify the extinction probability density function (the probability that a cluster ceases to proliferate), $\gamma(t) = -\frac{d}{dt} p_{\mathrm{surv}}$, and since $p_{\mathrm{surv}}(0) = 1$, we get the steady state condition

$$r(t) = \int_{t'<t} r(t') \gamma(t-t')\, dt' = \langle r(t-t') \rangle_{t'<t} \tag{7}$$

where $\langle ... \rangle_{t'<t}$ denotes the expected value with respect to the probability density $\gamma(t)$ restricted to $t' < t$. The right hand side is the total rate of clusters ceasing at time $t$, which is related to the mean initiation rate in the past. To illustrate the meaning of this function, let us consider the case that clusters have a fixed lifetime $t_l$. In that case, the lifetime distribution is $\gamma(t-t') = \delta(t-t')$ (the Dirac Delta-function), and thus,

$$r(t) = r(t - t_l) \tag{8}$$

Note that this must hold for all times $t$ during which the clusters are stationary, hence over this time period the cluster initiation rate must remain constant. This argument remains valid as long as $\gamma$ does not explicitly depend on time, since this means that the expected value in the past must be equal to the current value. Only if the form of the lifetime distribution changes over time in an exactly fine-tuned manner, to always compensate changes of initiation rates at the right point in the future, could the steady state be maintained, despite varying values of $r(t)$. This seems highly unlikely, given hat there is no known mechanism which could achieve this fine-tuning. On the other hand, if $r(t) = r_0$ is constant over time, then Eq. ((7)) is always fulfilled.

Thus we can conclude that **the cluster initiation rate is most likely constant over the time period of 3-5 weeks post-lesion**, and very possibly also beyond this time window.

### 2. The model for cluster turnover

We now establish the outlines for a model of cell proliferation and differentiation dynamics in proliferative clusters.

From Figures 3H,I and 5E, main text, we see that cluster composition changes from predominantly TAPs (cK cells) to pNBs (cKD cells) and then postmitotic NBs (cD cells) over time. Hence, we assume that the first cells after cluster initiation and division of the founding astrocyte are of cK type, which then differentiate to cKD cells and eventually to cD cells. We can thus establish the differentiation hierarchy: cK → cKD → cD.

Furthermore, both cK and cKD cells must be dividing ,since they are expressing Ki67. Our model, as the data on clusters does, will only consider cycling cells. Hence, we have the following possibilities for cell fate dynamics:

1. Differentiation may be coupled to cell division and thus cK cells choose their fate immediately upon division:

$$cK \xrightarrow{\lambda_K} \begin{cases} cK + cK & \text{with probability } r_K(1-\delta_K) \\ cK + cKD & \text{with probability } 1 - 2r_K \\ cKD + cKD & \text{with probability } r_K(1+\delta_K) \end{cases} \tag{9}$$

   where $\lambda_K$ is the division rate of cK cells (*rates* are generally defined as the inverse of the mean time between events). The cell fate probabilities are parametrised by $r_K$, the proportion of symmetric divisions, and $\delta_K$, the bias towards differentiation into cKD cells.
2. cK cells acquire DCX without cell division, differentiating into cKD cells:

$$cK \xrightarrow{\gamma_K} cKD \tag{10}$$

   where $\gamma_K$ is the differentiation rate of cK cells, that is, the rate at which they acquire DCX expression.

Within this model definition, we can identify two clear-cut cases:

- $\gamma_K = 0$ means that cell differentiation can only happen upon division, i.e. it is completely coupled to cell division.
- $\delta_K = -1, \gamma_K > 0$ means that cell differentiation is entirely uncoupled from cell division, since all divisions are symmetric and do not produce differentiated cells.

The same applies to cKD cells, which can differentiate and exit cell cycle to become cD cells:

$$cKD \xrightarrow{\lambda_D} \begin{cases} cKD + cKD & \text{with prob. } r_D(1-\delta_D) \\ cKD + cD & \text{with prob. } 1-2r_D \\ cD + cD & \text{with prob. } r_D(1+\delta_D) \end{cases} \tag{11}$$

and

$$cKD \xrightarrow{\gamma_D} cD \ . \tag{12}$$

As our model only considers the proliferating part of clusters, differentiation into cD cells, which are not cycling, is modelled as cell loss.

Finally, we consider that the propensity to differentiate may change with cluster age, i.e. the associated parameters either explicitly depend on the time since cluster initiation, $t_c$ (through an internal or external signalling clock), or on the number of undergone cell divisions $m$. This is expressed in terms of functions, $\delta(t_c, m), \gamma_K(t_c, m)$ and we will test various functional forms.

In the following, we will model those processes as a Markov process, that is, we assume that events occur stochastically, with rates only depending on the current configuration of the simulated cluster. This will be implemented and evaluated through stochastic computer simulations using a Gillespie algorithm [1] to predict cluster compositions, which we then compare to the experimental data. form the confetti data, Figure 4G, we see that 41 out of 43 clusters contained one or no astrocyte, and all astrocytes were dormant. This shows that in 95.3% of all clusters, astrocytes initially divide asymmetrically, producing one cK cell (pNB) and one astrocyte, which becomes dormant and either stays associated to the cluster (1 dormant astrocyte) or dissociates from it (no dormant astrocyte). We will thus simulate individual clusters, starting with one cK cell initially, which corresponds to the first cK cell **after** the first asymmetric division of an astrocyte, while the latter becomes dormant and thus lost from the model scope. By fitting and validating the model(s) on the data, we will first try to exclude most options and filter out the possible fitting model options. Note that by modelling the dynamics stochastically, we do not necessarily assume that the underlying biological processes are stochastic, but the probabilistic notion takes into account the uncertainty related to influences from the cellular environment that may affect cell fate, which we do not model explicitly. The biological processes may or may not be stochastic.

We note that in order to keep the number of parameters low enough to provide statistical meaningful fitting results and to avoid overfitting, the model makes certain simplifications. Most of those simplifications do not affect the the outcomes, while some deviations may occur, as discussed in the following:

- By assuming the model to be a Markov process, it is implicitly assumed that event timings follow an exponential probability distribution. While in reality, events such as cell division times do not follow such a distribution, we make use of the property of *weak convergence* (a generalisation of the central limit theorem) [2, 3], meaning that after multiple events have occurred, the detailed distribution of the individual event times does not matter, and only the mean event times, defined by the stochastic rates, matter to distinguish model outputs. This means the predictions of the here implemented Markov model will be identical, after long times, to a model with the real distribution and the same stochastic rates. Nonetheless, there may be deviations due to this assumption, at short simulation times, when not many events have occurred, and in the tails of of predicted outcome distributions.

- While we do not explicitly model the loss of cK cells – that is either by exiting the cell cycle, death, or dissociation from a cluster – these are indirectly taken into account via possible fast transitions cK $\rightarrow$ cKD $\rightarrow$ cD (whereby cD cells are considered as lost in this model) which can occur since the Markov property allows events to occur arbitrarily quickly (though with lower probability). Such events can be accommodated by inclusion in the stochastic rates $\gamma_{K,D}$, and by adjusting the distribution of event times. Nonetheless, as argued above, the latter does not matter in the long term for the bulk of the output distribution (that is, the distribution of cluster sizes), but may lead to deviations in its tails.

- The model does not explicitly account for cells that have exited the cell cycle to re-enter the cell cycle (temporary quiescence). Nonetheless, from a modelling standpoint, this simply means that the time period between cell division events (the last before exiting the cell cycle and the first before re-entering it) is significantly longer than the mean time between cell division events. This is in principle covered by the broad event-time distribution of a Markov process, while if such events occur in significant numbers it may result in an altered distribution of event times. However, as argued above, this does not affect model outcomes in the long term, but distributions at short times may be affected.

### 3. Fitting and validation procedure

To fit and validate the Markov model defined by (9) - (12), we compute the model likelihood with respect to the data: the statistical distribution of cluster sizes and their composition of cK and cKD cells. To that end, we run simulations of the Markov model defined above, following a kinetic Monte Carlo (Gillespie) algorithm [1]. The relative frequencies of simulated clusters with $n_K$ cK cells and $n_D$ cKD cells give an estimate for the probability $p_{n_K,n_D}$ to find simultaneously $n_K$ cK cells and $n_D$ cKD cells, as well as for the single cell-type probability distributions $p_{n_K}$ (to find $n_K$ cK cells irrespective of $n_D$) and $p_{n_D}$ (to find $n_D$ cKD cells, irrespective of $n_K$). We note that since for the data only clusters with 4 or more cells were counted, we do the same for the simulated clonal statistics. The simulation results are then compared with the data, the measured total counts $f_{n_K,n_D}$, as well as $f_{n_K}$ and $f_{n_D}$ (defined analogous to the respective probabilities). Since the outcomes of individual simulations – simulating a single cluster each – are independent of each other, the probability that the model with given probabilities $p_n$, where $n$ stands for $n_K, n_D$ or the combination $(n_K, n_D)$ respectively, to reproduce exactly the data $f_n$, is $P(D,\theta) \propto \prod_n [p_n(\theta)]^{f_n}$, where the sum and product go over all configurations of $n$. By normalising the probability distribution, one obtains a multinomial distribution for the likelihood,

$$L(\theta) = P(D,\theta) = \frac{(\sum_n f_n)!}{\prod_n f_n!} \prod_n [p_n(\theta)]^{f_n} \quad . \tag{13}$$

To get an estimate for the likelihood function we simulate various model versions for a wide range of parameter value configurations which form a tight mesh in parameter space, and determine the likelihood according to Eq. (13). We then plot the likelihood function. As explained in the following, we will reduce the model versions so far that we have only one or two fit parameters for each model fit, for which the likelihood function $L(\theta)$ can be visualised (for convenience, we will rather show the logarithm of the likelihood, the log-likelihood, *log-L* $= \ln(L)$, where ln is the natural logarithm).

*3.1. Finding the best model*

Usually, the model and parameters with the maximum likelihood are chosen as the best model/parameter combination. We would then accept or reject a model and parameters based on whether the deviation of the model prediction from the data is statistically significant. However, several aspects may confound such a straightforward approach:

- The prediction of the probability $p_n$ is stochastic itself due to the stochastic nature of the simulations. As such, the best likelihood estimate is prone to stochastic variations, leading to a very rugged likelihood landscape $L(\theta)$, with many local maxima. In particular, due to the stochasticity, the global maximum coming from simulations of a finite number of clusters may not coincide with the true global maximum, which one would get from an infinite number of simulated clusters.

- The best fit can often be achieved with more than one set of parameters. Hence, the likelihood maximum is often not unique, and this further increases the problem from the previous point.

- While the model can take into account simultaneous changes in division and differentiation rates of cells, through external factors and internal clocks, the model does not explicitly account for cooperative signalling interactions between cells that lead to simultaneous divisions, differentiation, or dissolution of clusters.

To address these issues,

- We choose to estimate the likelihood function $L(\theta)$ for a wide range of parameter set values rather than just a single maximum. Then, we consider a range of parameters $\theta$ with values $L(\theta)$ close to the maximum and sample parameters from this range to display the simulation results overlaid with the data. From this, the goodness of fit can be estimated.

- We do not reject models merely based on the statistical significance of deviations. Since such deviations may occur due to artefacts coming from the Markov approximation of the dynamics, or the lack of modelled cell-cooperative effects, even the best reasonable model (with our assumptions) may show such deviations. Instead, we inspect the structure of the predicted and measured cluster size distributions and see whether those are reasonably matched. Among the non-rejected models we then compare them based on numerical similarity between model and data and can thereby determine which model is more likely to be true, without strictly rejecting others.

- If discrepancies between model and data remain even for the best fit of the model, we will discuss what factors, not included in the model (such as the non-Markovian nature of the model and collective effects) can cause such discrepancies.

### 4. Model fitting and selection: results

The model as a whole contains at least 8 parameters, a number which further increases if cluster-age dependence is also considered. Since greedy optimisation algorithms, such as gradient descent, cannot be employed due to the potentially large number of local maxima of the likelihood function (see previous section), the number of parameters needs to be drastically reduced to avoid overfitting. To this end, we will use data from other experiments to fix some parameters before fitting on the cluster data, and then we will consider particular sub-versions of the model, with fewer (one or two) free parameters and will fit those on the data. By excluding most of these versions, we seek to arrive at a final version of a candidate model with few parameters which then can be fitted to the full set of data.

As a convention, we express all times in units of "average **c**ell **c**ycle times of cK cells" (cc) and associated rates as "per cell cycle" ($cc^{-1}$). This allows to eliminate one free fit parameter. We will later, in section 5, determine the unit "cc" in terms of common time units, from separate experimental data.

#### *4.1. Fitting cK-cells only*

Since, in the model, there are no transitions from cKD to cK cells, the cK-cell population in a cluster evolves independently from the cKD cell population. Thus, we first model only the cK population in a cluster and fit the model to the measured distribution of cK cells in clusters. We will then, in the subsequent section, use the best fit for cK cells to extend the model to include cKD cells and fit their distribution in clusters. For convenience, we will drop the subscripts of parameters $\delta_K, r_K, \gamma_K$ if it is clear from the context.

*4.1.1. Deterministic differentiation*

First, we consider a model version in which differentiation occurs precisely at a certain cluster age, that is, either at a specific time point or after a specified number of cell divisions after cluster initiation. Hence, we consider two model sub-variants:

$A.t$: Differentiation of cK cells occurs when a specified time $t_c = t_0$ has passed since the initiation of the cluster.

$A.m$: Differentiation occurs after a specified number of cell divisions $m = m_0$.

Thus, we assume that cK cells first only divide symmetrically, $cK \rightarrow cK + cK$ until time $t_0$ (version $A.t$) or for the first $m_0$ cell divisions (version $A.m$), upon which they differentiate. This can be implemented in our generic model by taking only processes (9) and (10) and assuming either (variant $A.t$):

$$\gamma = \begin{cases} 0 & \text{for } t < t_0 \\ \infty & \text{for } t \geq t_0 \end{cases} , \tag{14}$$

or (variant $A.m$):

$$\delta = \begin{cases} -1 & \text{for } m < m_0 \\ 1 & \text{for } m \geq m_0 \end{cases} . \tag{15}$$

We can easily see that model $A.t$ cannot be fitted to the data. Since, according to this model, all cK cells become cKD cells at the same time, clusters containing a mix of cK and cKD cells could not exist. Thus, the presence of such mixed clusters in the data tells us that model $A.t$ cannot be correct.

Hence, we only need to fit model $A.m$ quantitatively to the cK-cell distributions in clusters, whereby $m_0$ is the only free parameter. The fitting results are shown in Figure 1 in terms of the log-likelihood functions: The top panel shows the log-likelihood as function of the fitting parameter, $m_0$, the bottom

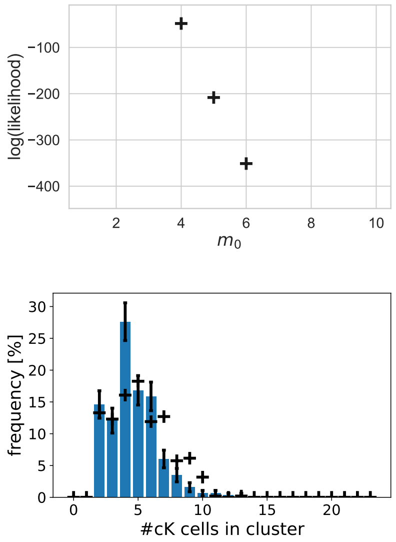


Figure 1: Fitting model $A.m$ to the distribution of cK cells in clusters. (Top row) log-likelihood for various values of parameters $m_0$. (Bottom row) Plots of cK-cell distributions in clusters for the value of $m_0 = 4$ with maximum likelihood. Crosses are simulation results and bars are experimental data. Corresponding figures for model $A.t$ are not shown since all likelihoods are zero.

panels show the predictions for the cK-cell distributions in clusters, according to the best fit parameter values, together with the experimental data.

We note that also for $A.m$, the best fit parameters do not produce a reasonably good match with the data. Therefore we reject both $A.t$ and $A.m$, and thus the hypothesis that differentiation occurs entirely deterministically.

##### *4.1.2. Constant differentiation rate (stochastic)*

The next model version we consider is when the time point of differentiation is uncertain, modelled as stochastic timing of events, while the propensities for doing so do not change over time. We consider two sub-variants:

$B.\delta$ Differentiation of cK-cells is coupled to cell division, i.e. $\gamma = 0$, while $\delta$ is a free parameter.

$B.\gamma$ Differentiation of cK-cells is *not* coupled to cell division, i.e. divisions simply duplicate cK cells, ensured by $\delta = -1$, while differentiation, $cK \to cKD$, occurs as separate event, whereby the differentiation rate $\gamma$ is a free parameter.

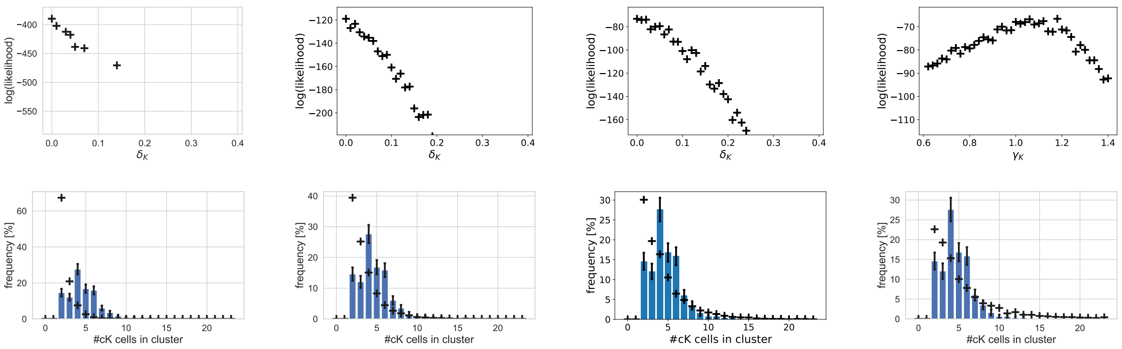


Figure 2: Fitting model $B$ to distributions of cK cells in clusters. (Top row) log-likelihood for various values of parameter $\delta_K$ for model $B.\delta$ and various values of parameter $\gamma$ for model $B.\gamma$ (parameters do not change with time). (Left column): model version $B.\delta$ with $r = 0.1$, (Left middle column): model version $B.\delta$ with $r = 0.3$, (Right middle column): model version $B.\delta$ with $r = 0.5$. (Right column): model version $B.\gamma$. (Bottom row) Plots of cK-cell distributions in clusters for the parameter values with maximum likelihood of corresponding plots above in the same column. Crosses are simulation results and bars are experimental data.

We note that also a mixture of both processes may be a possibility, however, since this means that in principle differentiation is not coupled to division, this would predict distributions very similar to variant $B.\gamma$.

In Figure 2 we show the fitting of the model variants. For variant $B.\delta$ we have free fit parameter $\delta$ for different fixed values of $r$, and $\gamma = 0$. For variant $B.\gamma$, we have free fit parameter $\gamma$ and $\delta = -1$ (in either cases, $\lambda$ is fixed as time unit). The top panels show the likelihoods as a function of $\delta$, for variant $B.\delta$ for cases $r = 0.1, 0.3, 0.4$ and for variant $B.\gamma$. The bottom panel shows the prediction for the distribution of cK cells for the parameter values with maximum likelihood (which is unique). As the models with best fit parameter for either model variant do not reproduce any reasonably good match with the data, we reject the hypothesis that differentiation rates are independent of cluster age.

#### *4.1.3. Cluster-age dependent differentiation rates*

In this section we study further model variants where the rates are not constant over time, but change with cluster age, be it with the number of cell divisions $m$ or explicitly as time $t_c$ passes after cluster initiation.

First, we assessed experimentally whether the cell division rate changes significantly over the course of maturation of a cluster. To that end we administered BrdU and harvested cells 2 hours later. BrdU labels cells in S-phase of mitosis, thus the labelling frequency from a single pulse of BrdU administration is approximately proportional to the cell division rate (given that the length of S-phase is not coupled to cell division rate and if labelling is not close to saturation). We compared the BrdU labelling efficiency in young, cK-only clusters, and more mature, mixed cK+cKD clusters. The result is shown in Table 1. Notably, the difference in BrdU labelling efficiency between cK cells in young and mature clusters is not significant, which led us to assume that the cell division rate does not change significantly over cluster age. Thus, we assume that the rate of division remains constant over cluster aging, while the proportion of differentiation events increases (in order be consistent with a finite cluster age, see section 5, the differentiation propensity cannot decrease substantially).

| cell types in cluster type | cK cells in cK clusters | cK cells in cK + cKD clusters |
|---|---|---|
| % BrdU+ | 52.9 | 43.1 |
| n | 81 | 47 |

Table 1: BrdU labelling frequency (% BrdU+) of cK cells in different cluster types, representing different stages of maturation (cK-only clusters are on average younger than mixed cK+cKD), together with sample sizes n. The p-value is $p = 0.117$ for BrdU percentage of cK cells in different cluster types. Thus the difference is not significant.

In the following, we model the increase of the differentiation propensity, $\gamma(X), \delta(X)$, respectively, where $X = t_c, m$, as either an exponential function,

$$\gamma(X) = \beta e^{\alpha X}, \qquad \delta(X) = \beta e^{\alpha X} - 1 \ , \tag{16}$$

or a power law,

$$\gamma(X) = \beta X^{\alpha}, \qquad \delta(X) = \beta X^{\alpha} - 1 \ . \tag{17}$$

Here, the parameters $\alpha$ and $\beta$ are the free fit parameters of this model version. The term "-1" for $\delta$ ensures that $\delta(X = 0) = -1$. Furthermore, $\delta$ is capped at $\delta = 1$ since this is associated with maximal symmetric differentiation probability equal to one. Note that the power law function includes the case of a linear increase for $\alpha = 1$.

We can implement different versions of the model of increasing differentiation propensity:

- As before, we implement model sub-versions when differentiation is coupled to cell division and when not. In the former case, we set $\gamma = 0$ and increase $\delta$ with cluster age, in the latter, we set $\delta = -1$ and increase $\gamma$ with cluster age. We will keep $r$ constant over cluster age, since it does not affect the propensity of differentiation (it is unbiased with respect to proliferation vs. differentiation).

- We can apply an exponential function (16) or power law (17) to model the variation over cluster age.

- We can have the parameters depend explicitly on the time $t_c$ since the cluster has been initiated, or on the number of undergone cell divisions $m$.

We denote model versions as follows: the first letter is the primary version ($C$ for main version C, i.e. cluster-age dependent differentiation propensity), the second symbol denotes which parameter varies with cluster age ("$\delta$" means that parameter $\delta$ increases with cluster age, starting from $\delta = -1$ while $\gamma = 0$; "$\gamma$" means that $\gamma$ varies with cluster age, starting from $\gamma = 0$, while $\delta = -1$), the third string of characters denotes the used function (*exp* for an exponential function according to (16), *pow* for a power law according to (17)), and the fourth entry denotes whether this function depends on time since cluster initiation, $t_c$ (here simply denoted as $t$) or the number of undergone cell divisions, $m$. In total, this gives 8 model versions, each with two fit parameters, $\alpha$ and $\beta$:

- For $C.\delta.exp.t$, parameter $\delta$ varies as $\delta(t) = \beta e^{\alpha t_c} - 1$

- For $C.\delta.exp.m$, parameter $\delta$ varies as $\delta(m) = \beta e^{\alpha m} - 1$

- For $C.\delta.pow.t$, parameter $\delta$ varies as $\delta(t) = \beta t_c^{\alpha} - 1$

- For $C.\delta.pow.m$, parameter $\delta$ varies as $\delta(m) = \beta m^{\alpha} - 1$

- For $C.\gamma.exp.t$, parameter $\gamma$ varies as $\gamma(t) = \beta e^{\alpha t_c}$

- For $C.\gamma.exp.m$, parameter $\gamma$ varies as $\gamma(m) = \beta e^{\alpha m}$

- For $C.\gamma.pow.t$, parameter $\gamma$ varies as $\gamma(t) = \beta t_c^{\alpha}$

- For $C.\gamma.pow.m$, parameter $\gamma$ varies as $\gamma(t) = \beta m^{\alpha}$

Furthermore, we fit versions for different, fixed values of $r$.

In Figures. 3-5, top panels, we show the log-likelihood functions of those model versions as a heatmap, as obtained from kinetic Monte Carlo simulations, where 'hotter' colours, according to the shown colour range, represent higher likelihoods, ($r = 0.5$ and $r = 0.3$ are shown separately for model version $C.\delta$, in Figures 3 and 4 respectively). It is notable that most of those functions do not show a single global maximum, but rather a curve along which the log-likelihood remains maximal. We therefore tested the models at two sets of parameter values, each, chosen as close as possible to the maximal likelihood, and we plotted predicted cK-cell distributions in clusters along the measured distributions, in the panels below the respective log-likelihood functions.

We see from the predicted distributions that for $r = 0.3$ substantial qualitative deviations between predicted and measured distributions remain, even for best fit parameter values. Hence we reject model versions $C.\delta$ with $r = 0.3$. For $r = 0.5$, predicted cK-cell distributions show a reasonable, qualitative fit in all four cases, but some data points still deviate significantly (by more than two standard deviations). We do not show versions of $C.\delta$ with $r < 0.3$ since the quality of fits decreases with decreasing $r$. Hence, we can exclude any significant contribution of asymmetric divisions, as their probability is $1 - 2r = 0$ for $r = 0.5$.

We also see reasonable fits for $C.\gamma$. In particular, for $C.\gamma.pow.t$ we see the best fit with highest likelihood. Furthermore, this model version is the only one for which the likelihood function has a unique maximum. We therefore identify this model version as the best fitting one, without strictly rejecting other versions. We note that this model implies that all divisions are symmetric, $cK \to cK + cK$ and differentiation occurs independently of cell division, $cK \to cKD$.

While those models of type $C$ yield reasonable fits for cK-cell distributions, we require a model that matches as accurately as possible the cK-cell distributions in order to go ahead and also fit the cKD-cell distributions. Hence, in the following we will seek to further improve the fit as a basis for modelling cKD cells.

##### *4.1.4. Cluster-age dependent differentiation rates with sharp cut-off*

Given that the tails of the distribution in model $C.\gamma.pow.t$ are slightly over-estimated in previous fits, we hypothesised whether a combination of the deterministic differentiation, version A, and varying differentiation rates

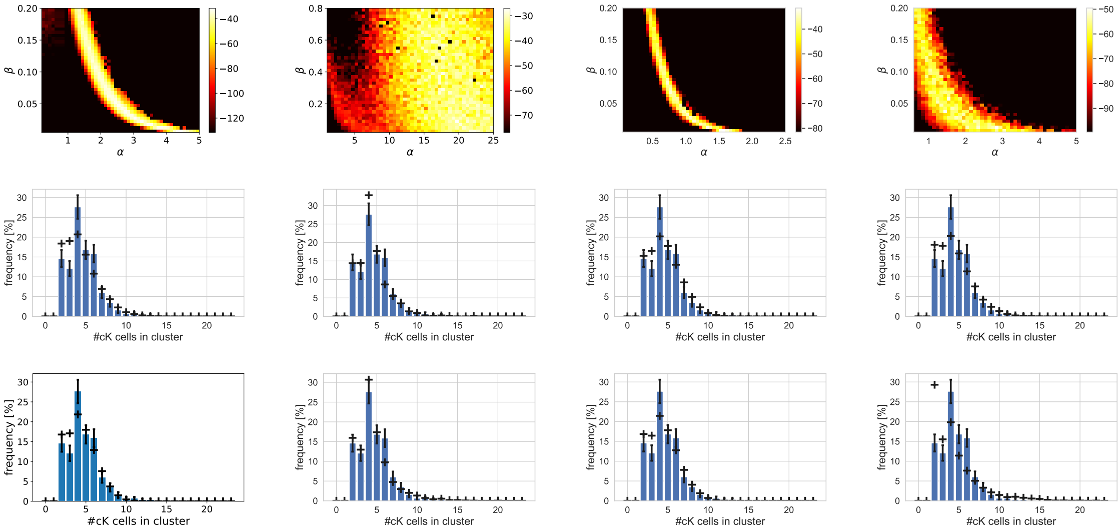


Figure 3: Fitting model versions $C.\delta$ with $r = 0.5$ to distributions of cK cells in clusters. The heatmaps show the log-likelihoods according to the adjacent colour scale bar. (Top row) log-likelihood as function of fit parameters $\alpha$ and $\beta$ for model subversions $C.\delta.pow.m$ (left column), $C.\delta.pow.t$ (left middle column), $C.\delta.exp.m$ (right middle column), $C.\delta.exp.t$ (right column). (Middle and bottom rows) Plots of cK-cell distributions in clusters for the parameter values with maximum likelihood of corresponding plots above in same column. Crosses are simulation results and bars are experimental data. The parameters of simulated cluster size distributions in the two bottom rows are: (in the order 'top', 'bottom', and in units of $cc^{-1}$): (left) $\alpha = 1.6, \beta = 0.135$ and $\alpha = 4.4, \beta = 0.005$, (middle left) $\alpha = 16.0, \beta = 0.58$ and $\alpha = 20.0, \beta = 0.005$, (middle right) $\alpha = 1.3, \beta = 0.01$ and $\alpha = 1.55, \beta = 0.005$, (right) $\alpha = 0.7, \beta = 0.08$ and $\alpha = 2.8, \beta = 0.01$.

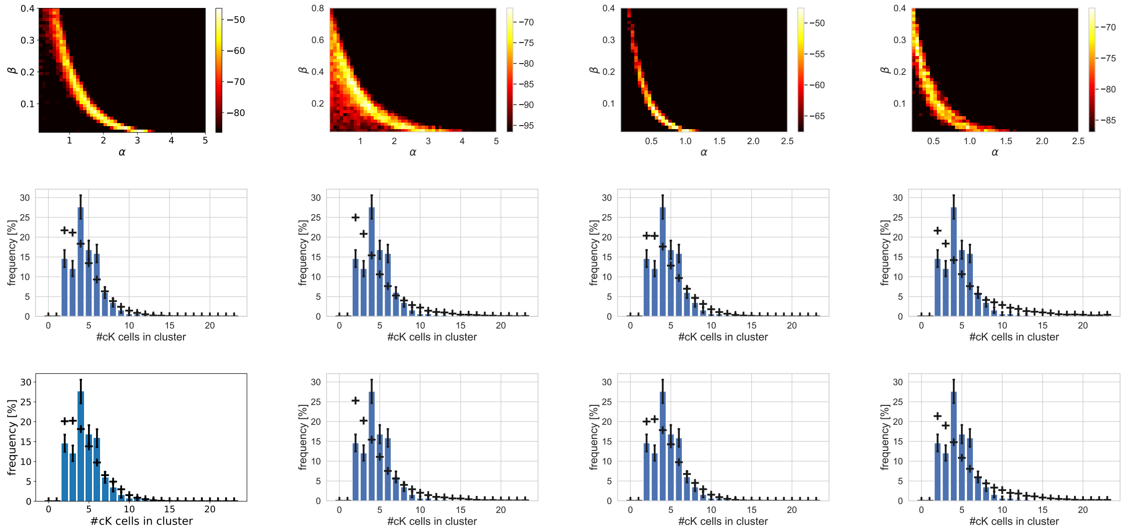


Figure 4: Fitting model versions $C.\delta$ with $r = 0.3$ to distributions of cK cells in clusters. The heatmaps show the log-likelihoods according to the adjacent colour scale bar. (Top row) log-likelihood as function of fit parameters $\alpha$ and $\beta$ for model subversions $C.\delta.pow.m$ (left column), $C.\delta.pow.t$ (left middle column), $C.\delta.exp.m$ (right middle column), $C.\delta.exp.t$ (right column). (Middle and bottom rows) Plots of cK-cell distributions in clusters for the parameter values with maximum likelihood of corresponding plots above in same column. Crosses are simulation results and bars are experimental data. The parameters of simulated cluster size distributions in the two bottom rows are: (in the order 'top', 'bottom', and in units of $cc^{-1}$ where relevant): (left) $\alpha = 1.7, \beta = 0.07$ and $\alpha = 3.0, \beta = 0.01$, (middle left) $\alpha = 0.7, \beta = 0.34$ and $\alpha = 1.2, \beta = 0.19$, (middle right) $\alpha = 0.5, \beta = 0.08$ and $\alpha = 1.0, \beta = 0.01$, (right) $\alpha = 0.95, \beta = 0.02$ and $\alpha = 0.25, \beta = 0.27$.

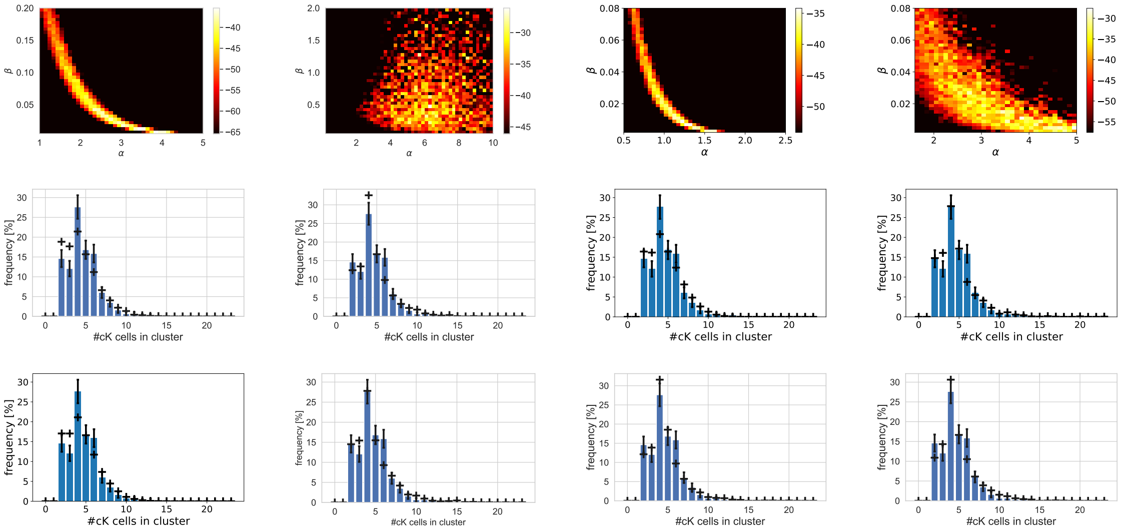


Figure 5: Fitting model versions $C.\gamma$ to distributions of cK cells in clusters. (Top row) log-likelihood as function of fit parameters $\alpha$ and $\beta$ for model subversions $C.\gamma.pow.m$ (left column), $C.\gamma.pow.t$ (left middle column), $C.\gamma.exp.m$ (right middle column), $C.\gamma.exp.t$ (right column). The heatmaps show the log-likelihoods according to the adjacent colour scale bar. (Middle and bottom rows) Plots of cK-cell distributions in clusters for the parameter values with maximum likelihood of corresponding plots above in same column. Crosses are simulation results and bars are experimental data. The parameters of simulated cluster size distributions in the two bottom rows are: (in the order 'top', 'bottom', and in units of $cc^{-1}$ where relevant): (left) $\alpha = 3.0, \beta = 0.0175$ and $\alpha = 3.9, \beta = 0.005$, (middle left) $\alpha = 6.1, \beta = 0.45$ and $\alpha = 4.0, \beta = 0.3$, (middle right) $\alpha = 1.35, \beta = 0.004$ and $\alpha = 1.6, \beta = 0.002$, (right) $\alpha = 3.8, \beta = 0.008$ and $\alpha = 4.7, \beta = 0.002$.

could define a model that matches the data better. We therefore now check the performance of model version $C$, but with the additional constraint that once the number of undergone cell divisions reaches $m_0$, cK cells always differentiate, to become cKD cells. We call this model version $C{-}A$. Here, we use $m_0 = 5$ as we do not observe any clusters with more than $2^4 = 16$ cK cells.

As in the previous section, we fitted the model for each of the 12 subversions (distinguished for variated parameter, type of function, independent variable, and $r = 0.3, 0.5$ for $C$-$A.\delta$). The result is shown in Figure 6, for all 12 model versions. Here, we only show the best-fit cK-cell distributions for model version $C$-$A.\gamma$, since the other versions show a substantially lower likelihood (The best fit distributions shown in the lowest panels are according to the best fits from the log-likelihood function immediately above, respectively). We see that, again, model version $C$-$A.\gamma$-$pow$-$t$ shows the best fit – which is better than the fit without cutoff – and has also a unique maximum in its likelihood function.

#### *4.2. Fitting cKD-cells*

We now take the best fitting model version and parameters from fitting cK cells, and include the distributions of cKD cells and associated parameters, for computing model likelihoods. Hence, we take model $C$-$A.\gamma.pow.t$, i.e. setting $\gamma_K(t) = \beta_K t^{\alpha_K}$ with $\alpha_K = 5.0$ and $\beta_K = 0.02cc^{-1}$, and we assume a differentiation cutoff at $m_0 = 5$. Based on these fixed parameters, we will now use the same model versions as used in the previous section for cK cell dynamics, but apply them to cKD cell dynamics parameters $\gamma_D$, $\delta_D$, $r_D$ instead.

We note that we do not use the joint cluster size distribution $f_{n_K,n_D}$ for comparison with the data, since this distribution is too sparse and thus the fitting is prone to noise. Instead, we use both the cluster size distributions of cK-cells, only, $D_1 = \{f_{n_K}\}$, and that of cKD-cells, only, $D_2 = \{f_{n_D}\}$, and compute the likelihoods as a Bayesian update: we use the Bayesian posterior coming from $D_1$,

$$P(\theta|D_1) = \frac{L(\theta|D_1)}{N(D_1)} \tag{18}$$

(where $N(D_1)$ is the Bayesian normalisation factor which depends only on the data, and where we assume a flat prior) as prior for the Bayesian posterior

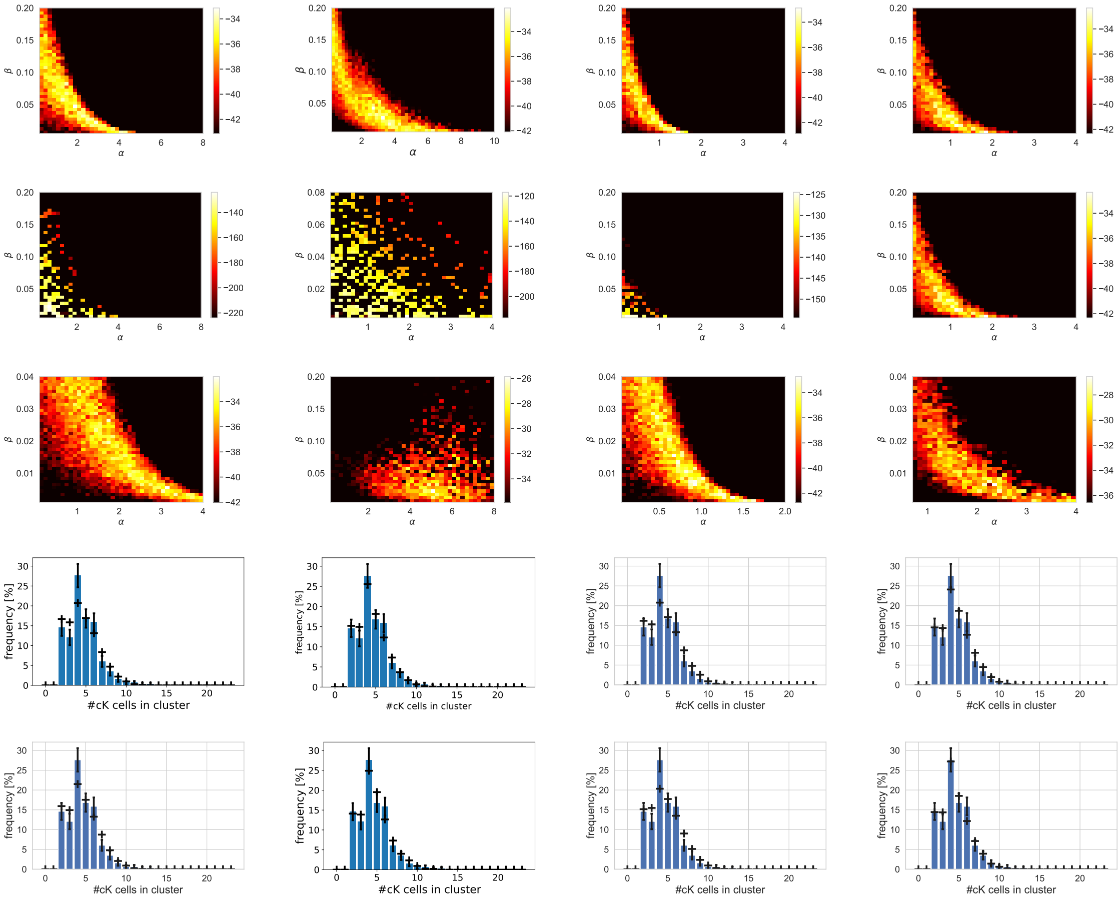

Figure 6: Fitting model versions $C\text{-}A.\delta$ with $r = 0.5$ (top row) and $r = 0.3$ (top middle row) and model version $C.\gamma$ (lower middle row) to distributions of cK cells in clusters. The *differentiation cutoff* is upon 5 cell divisions, when cK cells differentiate for sure, cK $\rightarrow$ cKD. The top three rows show the log-likelihood as function of fit parameters $\alpha$ and $\beta$ for the corresponding model subversions. The heatmaps show the log-likelihoods according to the adjacent colour scale bar. (Bottom rows) Plots of cK-cell distributions in clusters for the model and the parameter values with maximum likelihood immediately above in the same column (i.e. only for model $C\text{-}A.\gamma$). Crosses are simulation results and bars are experimental data. The parameters of simulated cluster size distributions in the two bottom rows are: (in the order 'top', 'bottom', and in units of $cc^{-1}$ where relevant): (left) $\alpha = 1.7, \beta = 0.019$ and $\alpha = 3.1, \beta = 0.004$, (middle left) $\alpha = 4.0, \beta = 0.03$ and $\alpha = 5.0, \beta = 0.02$, (middle right) $\alpha = 0.7, \beta = 0.013$ and $\alpha = 1.25, \beta = 0.002$, (right) $\alpha = 1.6, \beta = 0.009$ and $\alpha = 2.3, \beta = 0.006$. Plots of cK-cell distributions for other models are not shown as the likelihoods are substantially smaller.

of $D_2$, i.e.

$$P(\theta|D_2, D_1) = \frac{L(\theta|D_2)P(\theta)}{N(D_1)} = \frac{L(\theta|D_2)L(\theta|D_1)}{N(D_1)N(D_2)} \tag{19}$$

Since $N(D_1)N(D_2)$ only depends on the data, we can maximise the Bayesian posterior by maximising the product of likelihoods, $L(\theta|D_1, D_2) = L(\theta, D_1) \times L(\theta, D_2)$. This is the log-likelihood function shown in Figure 7, top three panels.

We note that model version $C$ includes model versions $A$ and $B$ as special cases:

- versions $A.m$ and $A.t$ corresponds to model version $C.\gamma.exp.m$ and $C.\gamma.exp.t$, respectively, when $\alpha \to \infty$ on the line $\beta = e^{-\alpha m}$ and $\beta = e^{-\alpha t_c}$, respectively.

- versions $B.\delta$ and $B.\gamma$ correspond to model versions $C.\delta.exp.t$ and $C.\gamma.exp.t$, respectively, when $\alpha = 0$.

Hence, it is sufficient to fit model version $C$, applied to cKD cells. We thus assume that $\delta_D(t_c, m), \gamma_D(t_c, m)$ follow the same functional form as $\delta_K(t_c, m), \gamma_K(t_c, m)$, yet with possibly different parameters $\alpha_D, \beta_D$. For cKD cells, also the cell division rate $\lambda_D$ could be different to that of cK cells. The BrdU labelling experiments displayed in Table 2 (labelling frequency measured 2 hours after BrdU labelling) show that there is a significant difference in labelling frequency of the two cell types, and that the labelling frequency of cKD cells is 0.73 times that of cK cells. Thus, a best estimate for the cKD cell division rate is $\lambda_D = 0.73 \times \lambda_K = 0.73cc^{-1}$, which we use in the following.

| cell type | cK | cKD |
|---|---|---|
| % BrdU+ | 57.5 | 42.1 |
| n | 262 | 361 |

Table 2: BrdU labelling frequency (% BrdU+) of cK and cKD cells (percentage), together with sample sizes $n$. The p-value for the difference in BrdU+ percentage between cell types is $1.18 \times 10^{-7}$, thus the difference is significant. The ratio of the labelling frequency of cKD vs. cK cells, 57.5/41.1 = 0.73, provides a best estimate for the ratio of cell division rates, as labelling frequencies are not saturated, thus $\lambda_D/\lambda_K = 0.73$.

We see that – when applied to cKD cell distributions – model versions $C.\delta$ does not yield reasonable fits, as the likelihoods are greatly lower than those of model version $C.\gamma$. Both model version $C.\gamma.pow.t$ and $C.\gamma.exp.t$ yield good fits, whereby $C.\gamma.pow.t$ is a slightly better match. For displaying the clone size distributions we chose two values with highest likelihood, although there is a complete line of equally well fitting parameters as visible in the plot, hence the best fit parameters are not unique. Nonetheless, for the following analysis we chose manually the **best fit parameters**,

$$\begin{aligned} \alpha_K &= 5.0cc^{-1} \\ \beta_K &= 0.02cc^{-1} \\ \alpha_D &= 5.0cc^{-1} \\ \beta_D &= 0.01cc^{-1} \end{aligned} \tag{20}$$

(21)

The majority of the distribution is matched well. However, deviations can be seen in the very tail of the distribution and for small cKD-cell numbers. This is generally expected for Markov models (see discussion in section 2 and a similar discussion in [4]). Deviations at small cKD cell numbers can be attributed to the tendency of clusters to dissolve at the end of their lifetime, upon which they are not counted for the data (bars): we observed that clusters which consist only of cKD and cD cells tend to appear as non-cohesive. These may still be cells in close proximity, and they do contain cycling cells, but were not counted due to the lack of cell-cell contact. The model, on the other hand, does not include the synchronous dissolution of clusters. While it would in principle be possible to model such a situation, this would require additional parameters which would render the fitting of the model infeasible.

In Figure 9, we also see that model version $C.\gamma.pow.t$ with the best fit parameters produces an excellent fit of the expected proportions of cluster types cK-only (TAP clusters), mixed cK-cKD clusters (TAP-pNB clusters), and cKD-only clusters (pNB clusters).

Notably, the best fits do not appear in the regions that correspond to model versions $A$ and $B$ as described above, hence we can again reject those model versions.

Given the high coherence of model and data in the bulk of the distribution, in particular for model $C.\gamma.pow.t$, we conclude that the differentiation propensity increases with cluster age and differentiation is independent of cell division ($cKD \to cKD + cKD$ and $cKD \to cD$). The deviations of the best model at small cell numbers, together with the experimental observation of disperse 'clusters' which were not scored as clusters due to the lack of cell-cell adhesion, suggests that this is due to a synchronous dissolution of clusters which is missed by the model.

### 5. Estimation of the average cell cycle time

In the previous sections, we have expressed all times in terms of average cell cycle times, "cc", that is, the inverse of the cell division rate. This is sufficient for simulations, as the time unit is arbitrary and can be chosen freely in simulations, and the experimental situation is in a steady state, as argued in section 1. Given the best fit parameters, we can now make predictions for the mean lifetime of clusters – defined as the time after which all its cells have exited the cell cycle – and by comparison with BrdU-tracing data we can determine an estimate for the cell cycle time and thus the cell division rate.

Simulating the best fit model, *C-A.γ.pow.t* for both cK and cKD cell dynamics with the best fit parameters according to 20, we obtain a mean lifetime of clusters of $t_l = 3.74cc$. Figure 10 shows the corresponding predicted lifetime distribution. To estimate the real mean lifetime from the data, we make use of the BrdU labelling experiments, where cells are labelled with BrdU at time point $t_0 = t_h - \Delta t$ where $\Delta t = 4\,\text{days}$ and $t_h$ is the harvesting time (Figure 3F in the main text). Here we measured the proportions $p_{B+} \approx 0.632$ of clusters that contain at least one BrdU+ cell when harvested, averaged over 3 harvesting times $t_h$. The fact that the variation between harvesting times is not large consolidates that the system is in a steady state with constant cluster birth rates, $r$ (see also section 1). When labelled 8h and 2h before harvesting we checked that almost all active proliferative clusters become labelled with BrdU for at least one cell (see Figure 3F in the main text), hence any cluster being initiated before $t_0$ is BrdU+ while any cluster initiated after that time point is BrdU-. Furthermore, only clusters that have been actively proliferating until $t_h$ can be measured, that is, any cluster initiated after the time point $t_b = t_h - t_l$ can be measured when harvested, where $t_l$ is the lifetime of that cluster. Thus, the total number of BrdU+ clusters, $N_{B+}$ is the number of clusters initiated between $t_b$ and $t_0$, and the total number of BrdU- clusters, $N_{B-}$, is the number of clusters initiated between $t_0$ and $t_h$. Denoting by $\bar{t}_l$ the mean lifetime of a cluster, we

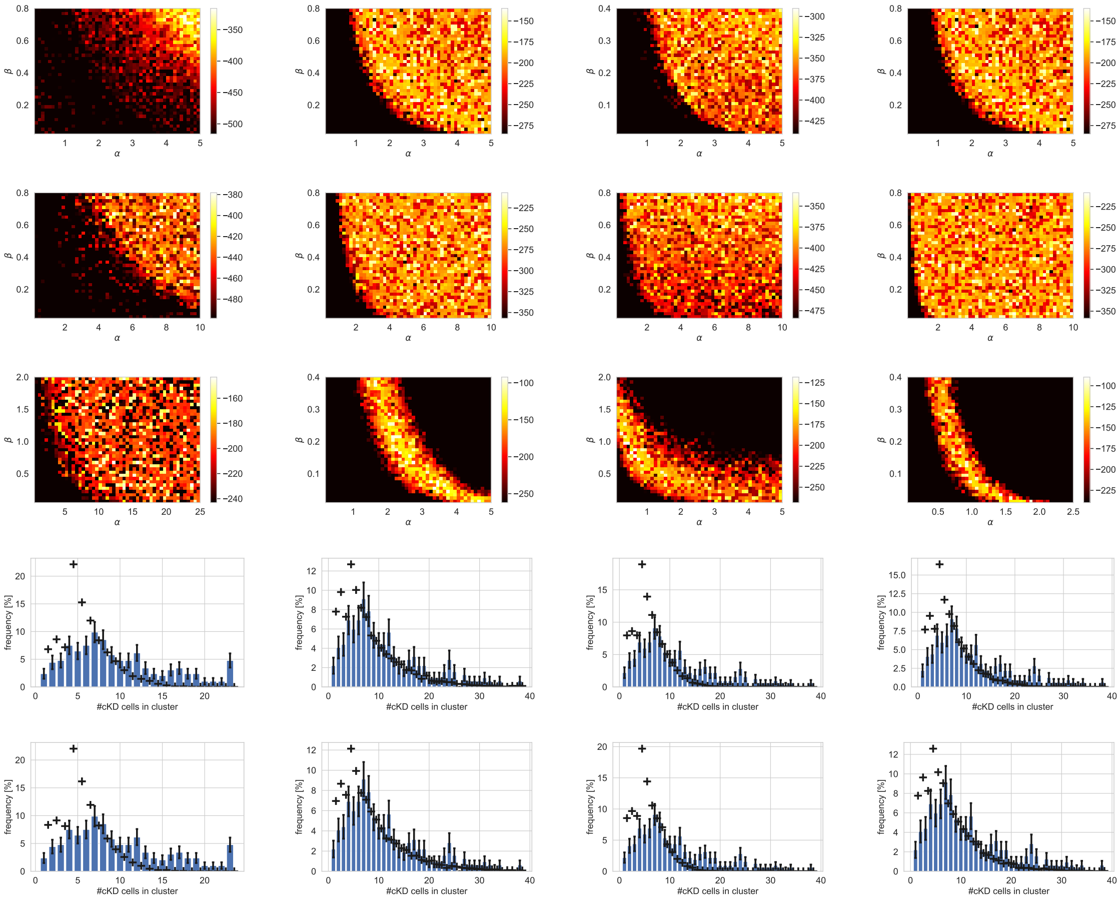


Figure 7: Fitting model versions $C.\delta$ and $C.\gamma$ – applied to cKD cell dynamics – to distributions of cK and cKD cells in clusters. All models displayed here assume best fit parameters for cK cell parameters, according to model $C$-$A.\gamma.pow.t$, i.e. $\delta_K = -1, \gamma_K = \beta_K t_c^{\alpha_K}$ with $\alpha_K = 5.0, \beta_K = 0.02$ (as in Figure 6.). Top three rows show log-likelihoods from joining the information from cK-cell and cKD cell distributions as in section 4.2, as function of $\alpha$ and $\beta$. The heatmaps show the log-likelihoods according to the adjacent colour scale bar. The model subversions are applied to cKD cell parameters $\delta_D$ and $\gamma_D$ only. (Top row) model $C.\delta$ and $r = 0.5$,(Top middle row) model $C.\delta$ and $r = 0.3$, (Lower middle row) model $C.\gamma$. (Bottom rows) Plots of cKD-cell distributions in clusters for various best fit parameter values of model $C.\gamma$ (Plots of cKD-cell distributions for other models are not shown as the likelihoods are substantially smaller). Crosses are simulation results and bars are experimental data. The parameters of simulated cluster size distributions in the two bottom rows are: (in the order 'top', 'bottom', and in units of $cc^{-1}$ where relevant): (left) $\alpha_D = 11.0, \beta_D = 0.65$, $\alpha_D = 23.0, \beta_D = 1.15$, (middle left) $\alpha_D = 5.0, \beta_D = 0.01$, $\alpha_D = 6.0, \beta_D = 0.003$, (middle right) $\alpha_D = 0.3, \beta_D = 2.0$, $\alpha_D = 1.5, \beta_D = 0.5$, (right) $\alpha_D = 0.65, \beta_D = 0.22$, $\alpha_D = 1.5, \beta_D = 0.02$.

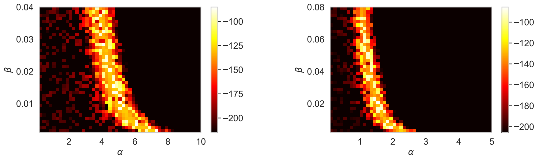


Figure 8: Log-likelihood landscapes, with model and parameters as in Figure 7 third row, for model *C-A.γ.power.t* (left) and *C-A.γ.exp.t* (right), but with higher parameter space resolution.

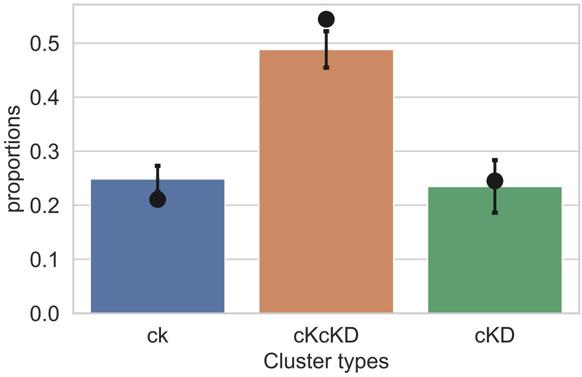


Figure 9: Proportions of cluster types (cK cells only (cK), mixed clusters (cKcKD), cKD cells only (cKD), as found in the data (bars), and as simulated by model *C.γ.pow.t* applied for both cK and cKD cells, for the best fit parameters according to (20) (points). Error bars are standard error of mean of cluster counts in the data.

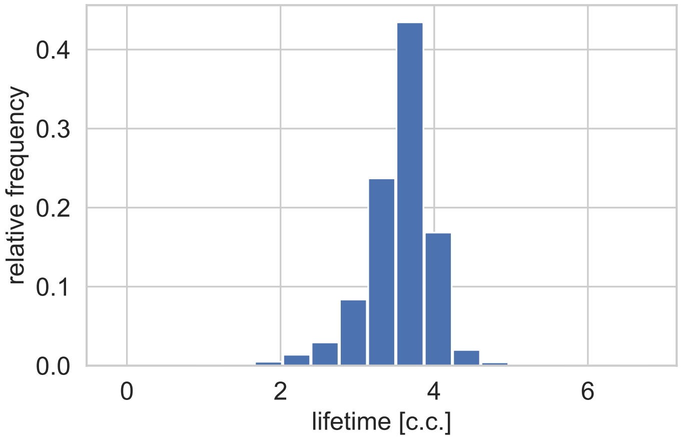


Figure 10: Distribution of cluster lifetimes. The horizontal axis shows cluster lifetimes, while the vertical axis shows frequencies.

have then

$$N_{B+} = r(t_0 - \bar{t}_b) = r(\bar{t}_l - \Delta t), \qquad N_{B-} = r(t_h - t_0) = r\Delta t \tag{22}$$

$$\implies \frac{N_{B+}}{N_{B-}} = \frac{p_{B+}}{p_{B-}} = \frac{\bar{t}_l - \Delta t}{\Delta t} = \frac{\bar{t}_l}{\Delta t} - 1 \tag{23}$$

$$\implies \bar{t}_l = \Delta t \left( \frac{p_{B+}}{1 - p_{B+}} + 1 \right) = 4\,\text{days} \times (\frac{0.623}{0.377} + 1) = 10.6\,\text{days} \tag{24}$$

From this follows a mean cell cycle time of around $t_{cc} = \bar{t}_l[\text{days}]/\bar{t}_l[cc] \approx 2.83\,\text{days}$, and hence, the cell division rate $\lambda = 1/t_{cc} = 0.353\,\text{day}^{-1}$.
