## Supplementary material for "Dynamic spatiotemporal activation of a pervasive neurogenic competence in striatal astrocytes supports continuous neurogenesis following injury": Table S1 - Statistical analyses

***Table S1. Results of all statistical analyses presented in the study***
* Indicates that a post hoc test was performed, please see below

| Graph | Statistical test | Result | P Value | Post-hoc |
|---|---|---|---|---|
| Fig. 1F | Linear Regression | AdjR^2= 0.8895 | 9.04E-4 | |
| Fig. 2D (Total cells) | Kruskal-Wallis Test | K-W chi squared = 212.4, df=4 | 8.24E-45 | * |
| Fig. 2D (TAPs) | Kruskal-Wallis Test | K-W chi squared = 120.5, df=3 | 6.12E-26 | * |
| Fig. 2D (pNBs) | Kruskal-Wallis Test | K-W chi squared = 125.4, df=3 | 5.30E-27 | * |
| Related to Fig. 2B (N of cells vs association with pmNBs) | Logistic Regression | AdjR^2= 0.31 | 5.96E-21 | |
| Related to Fig. 2B (% of pNBs vs association with pmNBs) | Logistic Regression | AdjR^2= 0.44 | 2.19E-25 | |
| Fig. 2H | Permutation Test using Monte Carlo Simulations | Moran's I coefficients | * | |
| Fig. S2F | Fisher's Exact Test | | 4.99E-4 | * |
| Fig. 3E | One-Way ANOVA | F(4,13)= 1.98 | 0.157 | |
| Fig. 3G | Mann–Whitney Test | Mann–Whitney U = 22678 | 6.907E-11 | |
| Fig. 3H | Mann–Whitney Test | Mann–Whitney U = 22711 | 1.955E-11 | |
| Fig. 3I | Fisher's Exact Test | | 2.932E-12 | |
| Fig. 3J | Independent samples t-test | t(6.6704)= 1.0764 | 0.3191 | |
| Fig. S4C (% of TAPs-only) | One-Way ANOVA | F(3,9)= 3.883 | 0.049 | |
| Fig. S4C (% of TAPs+pNBs) | One-Way ANOVA | F(3,9)= 0.23 | 0.873 | |
| Fig. S4C (% of pNBs-only) | One-Way ANOVA | F(3,9)= 1.824 | 0.213 | |
| Fig. S4D (N of cells in TAPs-only) | Kruskal-Wallis Test | K-W chi squared = 2.5423, df=3 | 0.4677 | |
| Fig. S4D (N of cells in TAPs+pNBs) | Kruskal-Wallis Test | K-W chi squared = 2.2104, df=3 | 0.5299 | |
| Fig. S4D (N of cells in pNBs-only) | Kruskal-Wallis Test | K-W chi squared = 6.9651, df=3 | 0.07302 | |
| Fig. S5A | Independent samples t-test | t(7.0858)= -5.9575 | 0.0005403 | |
| Fig. S5B (TAPs vs pNBs) | Paired samples t-test | t(3) = 3.4728 | 0.04026 | |
| Fig. S5B (TAPs in TAPs-only vs TAPs in TAPs+pNBs) | Paired samples t-test | t(3) = -2.1131 | 0.125 | |
| Fig. S5B (pNBs in TAPs+pNBs vs pNBs in pNBs-only) | Paired samples t-test | t(3) = 2.0584 | 0.1317 | |
| Fig. S5C | One-Way ANOVA | F(3,9)= 0.826 | 0.512 | |
| Fig. S5D | Anderson-Darling k-sample T | | 0.35636 | |
| Fig. S5E | Kolmogorov-Smirnov Test | D = 0.3427 | 1.51E-14 | |
| Fig. S5F | Kolmogorov-Smirnov Test | D = 0.093482 | 0.247 | |
| Fig. 4F | Linear Regression | AdjR^2= 0.88 | 0.042 | |
| Fig. 4H | Linear Regression | AdjR^2= 0.53 | 0.004 | |
| Fig. 4I | Monte Carlo Permutation Test | Moran's I coefficients | * | |
| Fig. S6E | One-Way ANOVA | F(2,9)= 1.571 | 0.26 | |
| Fig. S6F | One-Way ANOVA | F(2,9)= 0.156 | 0.858 | |
| Fig. S6G | One-Way ANOVA | F(2,9)= 0.834 | 0.512 | |
| Fig. S6H | One-Way ANOVA | F(2,9)= 1.834 | 0.219 | |
| Fig. S6K (TAPs-only) | Fisher's Exact Test | | 1 | |
| Fig. S6K (TAPs+pNBs) | Fisher's Exact Test | | 0.454 | |
| Fig. S6K (pNBs-only) | Fisher's Exact Test | | 0.301 | |
| Fig. 5D | One-Way ANOVA | F(3,10)= 14.550 | 0.00056 | * |
| Fig. 5E (TAPs-only) | One-Way ANOVA | F(3,10)= 4.486 | 0.03056 | * |
| Fig. 5E (TAPs+pNBs) | One-Way ANOVA | F(3,10)= 7.520 | 0.00638 | * |
| Fig. 5E (pNBs-only) | One-Way ANOVA | F(3,10)= 17.801 | 0.00025 | * |
| Fig. 5J | Fisher's Exact Test | | 6.428E-05 | * |
| Fig. 6K | Mann–Whitney Test | Mann-Whitney U = 1161 | 0.01312 | |

| | | | | |
|---|---|---|---|---|
| Fig. 6N | One-Way ANOVA | F(2,6)= 68.52 | 7.38E-08 | * |
| Fig. 6O | One-Way ANOVA | F(2,6)= 2.435 | 0.168 | |
| Fig. 6P | One-Way ANOVA | F(2,6)= 0.73 | 0.52 | |
| Fig. S7A (Total) | Kruskal-Wallis Test | K-W chi squared = 37.595, df=3 | 3.443E-08 | * |
| Fig. S7A (100%) | Kruskal-Wallis Test | K-W chi squared = 52.062, df=3 | 2.905E-11 | * |
| Fig. S7B (Total) | Kruskal-Wallis Test | K-W chi squared = 43.687, df=3 | 1.759E-09 | * |
| Fig. S7B (100%) | Kruskal-Wallis Test | K-W chi squared = 31.305, df=3 | 7.334E-07 | * |
| Fig. S7F (including the 5 orange) | Wilcoxon signed-rank Test | V = 63.5 | 0.0414 | |
| Fig. S7F (excluding the 5 orange) | Wilcoxon signed-rank Test | V = 52 | 0.2512 | |
| Fig. S7G (including the 5 orange) | Wilcoxon signed-rank Test | V = 12 | 0.03756 | |
| Fig. S7G (excluding the 5 orange) | Wilcoxon signed-rank Test | V = 9 | 0.4469 | |
| Fig. S7H (SOX9+Ascl1- cells) | Fisher's Exact Test | | 1.014E-11 | * |
| Fig. S7H (SOX9+Ascl1+ cells) | Fisher's Exact Test | | <2.2E-16 | * |
| Fig. S7H (SOX9-Ascl1+ cells) | Fisher's Exact Test | | 4.492E-10 | * |

### Post hoc analyses

| **Figures 2D and S2F. Post hoc** | Figure 2D Dunn's Test for multiple comparison | | | Figure S2F |
|---|---|---|---|---|
| Comparison | Total cells | TAPs | pNBs | Pairwise Fisher's Exact Test |
| TAPs-only - TAPs+pNBs_Low | 0.0412478 | 0.179955 | | 0.0303 |
| TAPs-only - TAPs+pNBs_Med | 8.74E-13 | 0.064751 | | 7.44E-09 |
| TAPs-only - TAPs+pNBs_High | 1.21E-44 | 9.51E-23 | | 1.85E-38 |
| TAPs-only - pNBs-only | 7.18E-22 | | | 1.31E-41 |
| TAPs+pNBs_Low - TAPs+pNBs_Med | 0.1065108 | 0.034045 | 0.024782 | 0.542 |
| TAPs+pNBs_Low - TAPs+pNBs_High | 6.74E-05 | 1.43E-08 | 3.47E-12 | 0.000136 |
| TAPs+pNBs_Low - pNBs-only | 0.0265778 | | 6.50E-11 | 1.64E-05 |
| TAPs+pNBs_Med - TAPs+pNBs_High | 7.84E-06 | 1.26E-10 | 1.72E-18 | 2.31E-09 |
| TAPs+pNBs_Med - pNBs-only | 0.2312437 | | 5.97E-15 | 1.36E-11 |
| TAPs+pNBs_High - pNBs-only | 0.0001211 | | 0.362679 | 0.542 |

| **Figure 2H. Moran's I analysis** | AnimalID | Moran's I Index | P value |
|---|---|---|---|
| Total cells | G14.3 | -0.003 | 0.34 |
| Total cells | G14.4 | -0.058 | 0.821 |
| Total cells | N2 | -0.033 | 0.683 |
| Total cells | N3 | -0.005 | 0.422 |
| Total cells | N1 | 0.013 | 0.388 |
| % of pNBs | G14.3 | -0.029 | 0.553 |
| % of pNBs | G14.4 | -0.085 | 0.516 |
| % of pNBs | N2 | -0.014 | 0.984 |
| % of pNBs | N3 | -0.019 | 0.727 |
| % of pNBs | N1 | 0.001 | 0.724 |
| Association w/ pmNB | G14.3 | -0.038 | 0.247 |
| Association w/ pmNB | G14.4 | -0.057 | 0.929 |
| Association w/ pmNB | N2 | 0.033 | 0.745 |
| Association w/ pmNB | N3 | -0.042 | 0.63 |
| Association w/ pmNB | N1 | 0.034 | 0.222 |

| Figure S4C (TAPs-only). Post hoc | Comparison | P value |
|---|---|---|
| | 3wpl-4wpl | 0.920 |
| | 3wpl-5wpl | 0.807 |
| | 3wpl-8wpl | 0.216 |
| | 4wpl-5wpl | 0.996 |
| | 4wpl-8wpl | 0.087 |
| | 5wpl-8wpl | 0.047 |

| Figure 4I. Moran's I analysis | Specimen | Moran's I Index | P value |
|---|---|---|---|
| | #C1 | -0.005 | 0.622 |
| | #C2 | -0.025 | 0.649 |
| | #C3 | -0.005 | 0.991 |
| | #C4 | -0.002 | 0.519 |

| Figures 5D and 5E. Tukey Post hoc | Total clusters | TAPs-only | TAPs+pNBs | pNBs-only |
|---|---|---|---|---|
| T-4 vs T-7 | 0.4186 | 0.9744 | 0.1374 | 0.7795 |
| T-4 vs T-14 | 0.0021 | 0.3591 | 0.0092 | 0.0006 |
| T-4 vs T-bQA | 0.0016 | 0.0369 | 0.0098 | 0.0053 |
| T-7 vs T-14 | 0.0150 | 0.5167 | 0.3027 | 0.0012 |
| T-7 vs T-bQA | 0.0096 | 0.0501 | 0.2642 | 0.0142 |
| T-14 vs T-bQA | 0.9383 | 0.3520 | 0.9949 | 0.5948 |

| Figure 5J. Pairwise Fisher's Exact Test | P value |
|---|---|
| T-4 vs T-7 | 1 |
| T-4 vs T-14 | 0.0039 |
| T-4 vs REF | 0.0033 |
| T-7 vs T-14 | 0.00984 |
| T-7 vs REF | 0.00984 |
| T-14 vs REF | 1 |

| Figure 6N. Tukey Post hoc | P value |
|---|---|
| N_Area vs N_Core | 0.989 |
| N_Area vs hSVZ | 0.00013 |
| N_Core vs hSVZ | 0.00014 |

| Figures S7A-S7B. Dunn's Test or multiple comparisons | Figure S6A (N of cells) | | Figure S6B (% of pNBs) | |
|---|---|---|---|---|
| Comparison | Total | 100% | Total | 100% |
| T-4 vs T-7 | 0.9090 | 0.0288 | 0.0740 | 0.0611 |
| T-4 vs T-14 | 0.0027 | 6.085E-05 | 7.069E-06 | 0.0006 |
| T-4 vs REF | 0.00001 | 4.558E-08 | 9.117E-09 | 1.621E-05 |
| T-7 vs T-14 | 0.0055 | 0.0265 | 0.0267 | 0.0613 |
| T-7 vs REF | 0.0001 | 3.658E-05 | 0.0030 | 0.0022 |
| T-14 vs REF | 0.2486 | 0.0293 | 0.5487 | 0.2340 |

| Figure S7H. Pairwise Fisher's Exact Test | SOX9+Ascl1- | SOX9+Ascl1+ | SOX9-Ascl1+ |
|---|---|---|---|
| Single VS Pair | 0.116 | 0.284 | 0.556 |
| Single VS Trio | 0.116 | 1 | 0.237 |
| Single VS Cluster | 4.36E-07 | 2.8E-03 | 1.05E-03 |
| Pair VS Trio | 1 | 0.106 | 0.237 |
| Pair VS Cluster | 1.68E-04 | 1.11E-19 | 1.36E-06 |
| Trio VS Cluster | 6.08E-03 | 2.1E-04 | 0.237 |
